# Cardiomyocytes in Persistent Cancer Therapy-related Cardiac Dysfunction Show Contractile and Natriuretic Peptide Signaling Signatures

**DOI:** 10.64898/2026.11.13.766492

**Authors:** Angel Chan, Jens Hansen, Ravi Iyengar

**Author notes:** Address for correspondence: Ravi Iyengar, PhD, Department of Pharmacological Sciences, Icahn School of Medicine at Mount Sinai, One Gustave L. Levy Place, Box 1215, New York, NY 10029, USA., E-mail: Ravi Iyengar, PhD, Angel Chan, MD, PhD, Jens Hansen, MD. Drs Chan and Hansen contributed equally to this work and are joint first authors.

## Abstract

**BACKGROUND:** Anthracycline and trastuzumab therapy of HER2-positive breast cancer causes cancer therapy-related cardiac dysfunction (CTRCD) in a subset of treated patients. Some recover cardiac function, others do not. Whether persistence is a cardiomyocyte-intrinsic property is unknown.

**OBJECTIVES:** We sought to determine whether induced pluripotent stem cell-derived cardiomyocytes (iPSC-CMs) from patients with persistent CTRCD respond to these drugs differently from those of patients with reversible or no CTRCD, and to identify the functional pathways involved.

**METHODS:** iPSCs were derived from 23 women with normal baseline cardiac function treated with doxorubicin-based chemotherapy followed by trastuzumab: 5, 11 and 7 with no, reversible, and persistent CTRCD, respectively. iPSC-CMs were exposed to 7 drug regimens, generating 161 transcriptomic profiles that we resolved by singular value decomposition and subjected to pathway enrichment analysis. Reliability was tested by 100 clinical label permutations and leave-one-patient-out downsampling.

**RESULTS:** Persistent CTRCD responses clustered more tightly and separated from reversible and no-CTRCD responses to a higher degree than expected by chance (P = 0.04). Reversible and no-CTRCD responses did not resolve from one another. Genes encoding guideline-recommended biomarkers, including natriuretic peptides and cardiac troponins, were more strongly induced or less repressed in persistent CTRCD cell lines, while natriuretic peptide receptor signaling was lower. Contraction, calcium handling, cytoskeletal and glycolytic pathways differed between CTRCD groups. Comparison with signatures induced by cardiotoxic kinase inhibitors in healthy donor cardiomyocytes documented high functional similarities.

**CONCLUSIONS:** Cardiomyocytes display a cell-intrinsic signature of persistent CTRCD that centers on contractile homeostasis and natriuretic peptide receptor signaling, supporting patient group-selective cardioprotective intervention strategies.

## INTRODUCTION

Anthracycline therapy followed by trastuzumab remains a standard regimen for high-risk HER2-positive breast cancer, and improvements in survival have made long-term cardiovascular outcomes a central concern for survivors.^1,2^ Cancer therapy-related cardiac dysfunction (CTRCD) still occurs in approximately 9% of treated patients, most of whom have no preexisting cardiovascular disease and a normal ejection fraction before treatment.^3^ Anthracycline-associated cardiac injury has been recognized since the 1970s,^4^ and a decline in cardiac function during trastuzumab therapy was evident in the earliest trials.^1,5^

Most patients with CTRCD recover left ventricular ejection fraction (LVEF) once cancer therapy is completed or halted and heart failure therapy is initiated, but a subset does not, and these patients carry the burden of chronic heart failure on top of their cancer diagnosis. Because there is currently no way to identify this subset before or early during treatment, oncologists and cardiologists must weigh interruption of effective cancer therapy against uncertain cardiac risk. Existing risk models rely on cumulative anthracycline dose, baseline LVEF, global longitudinal strain and comorbidity burden, and perform only modestly at the level of the individual patient.^3,6^

Patient-specific induced pluripotent stem cell-derived cardiomyocytes (iPSC-CMs) provide a way to ask whether susceptibility is an intrinsic property of the patient’s cardiomyocytes. Using iPSC-CMs from healthy donors, we previously identified transcriptomic signatures shared by cardiotoxic tyrosine kinase inhibitors (TKIs) that map to sarcomere dynamics, calcium handling and energy metabolism.^7,8^ Outlier analysis linked a distinct response to the RARG variant rs2229774, a known genetic risk allele for anthracycline cardiotoxicity.^9^

We therefore hypothesized that iPSC-CMs derived from patients who developed persistent CTRCD would respond to doxorubicin and trastuzumab differently from cardiomyocytes derived from patients who recovered or who never developed CTRCD and that the pathways distinguishing these groups would point toward measurable clinical correlates. We tested this hypothesis in cardiomyocytes derived from 23 women with HER2-positive breast cancer and known long-term cardiac outcomes.

## METHODS

### Study approval

The protocol was approved by the Memorial Sloan Kettering Cancer Center Institutional Review Board, and all procedures were HIPAA compliant. Written informed consent was obtained from all participants. Samples were deidentified before transfer to the Icahn School of Medicine at Mount Sinai, where all experiments and computational analyses were performed.

### Patients

Women treated with doxorubicin, cyclophosphamide and paclitaxel followed by trastuzumab at Memorial Sloan Kettering Cancer Center were identified retrospectively by chart review. Eligible patients were adults with histologically confirmed HER2-positive breast cancer, normal LVEF (≥53%) by echocardiography before treatment, and serial imaging during therapy. CTRCD was defined as a fall in LVEF to <53% with a decrement of >10% from baseline, corresponding to moderate to severe CTRCD in current classifications.^6^ Patients with obstructive coronary artery disease, prior LV dysfunction or heart failure, significant valvular disease, or significant arrhythmia were excluded. Patients with CTRCD were classified as reversible (recovery to LVEF ≥53%) or persistent based on recovery status at 48 months after the last dose of trastuzumab; the persistent group was followed for a median of 71.2 months.

### Cardiomyocyte derivation and drug exposure

Peripheral blood mononuclear cells were reprogrammed with Sendai virus vectors, and 2 fully characterized iPSC clones per patient were differentiated to cardiomyocytes by monolayer Wnt modulation followed by lactate selection, as described previously.^7^ Purity and cardiac identity were confirmed by immunostaining, reverse transcription quantitative polymerase chain reaction and flow cytometry. Cardiomyocytes were used 30 to 45 days after differentiation and exposed to 7 regimens: doxorubicin 0.1 µmol/L for 24 hours; low- or high-dose trastuzumab (200 or 400 µg/mL, respectively) for 48 hours; doxorubicin for 24 hours followed by addition of either trastuzumab dose for another 48 hours; or doxorubicin combined simultaneously with either trastuzumab dose for 48 hours. The doxorubicin concentrations correspond to clinically observed serum concentrations.^7^ The low trastuzumab concentration approximates peak serum concentration sat steady state during 3-weekly dosing, and the high concentration is twice that level.^8^ Untreated cells served as controls. All exposures were performed in duplicate for each of 2 clones per patient.

### Transcriptomics and identification of group-selective responses

Bulk RNA sequencing was performed on all conditions and analyzed as described previously.^7,8^ Briefly, bulk transcriptomic data were aligned to the human reference genome hg38 using STAR,^10^ read counts were mapped to genes using the RefGene reference annotation and feature counts.^11^ We used the negative-binomial test implemented in the ‘exactTest’ functionality of edgeR ^12^ to identify differentially expressed genes between treated and control cell lines, yielding 161 response profiles. P values were transformed into -log_10_ P values and defined as positive or negative for up- or downregulated genes, respectively. Before the transformation, we replaced P values that were zero with 10^-64^, since this is the lowest P value that is generated by our edgeR pipeline.

### Definition of sets that combine multiple cell line/treatment combinations

We defined sets of DEG vectors for which we searched for common vector components, using six different set annotations (Suppl. Fig. 1C). The first annotation grouped all DEG vectors induced by the same in vitro treatment (T annotation), the second all DEG vectors induced in cell lines of the same CTRCD group (3 CT groups), the third all DEG vectors induced in either no- or reversible and persistent CTRCD lines (NvsCT) and the fourth all DEG vectors induced in either persistent or non-persistent cell lines (PvsNP). The fifth and sixth annotations separated the 3 and 2 sets of the 3 CT groups and PvsNP annotations into smaller sets that contained all cell lines with the relevant CTRCD and were treated with the same regimen (T & 3 CT group, T & PvsNP, respectively). Since our algorithm will split existing sets into new sets (see below), we refer to those sets as initial sets.

### Pairwise correlation analysis and clustering

The following data processing and analysis steps are identical to the ones applied in our previous publication,^7^ except that we applied them to the defined sets of the six different annotations. We calculated the Pearson correlation coefficients between all pairs of the 161 DEG vectors and subjected the results to hierarchical clustering (distance function: (1-cor(x))/2). To quantify the clustering efficiency for each set, our algorithm cut the cluster dendrogram at any potential height that gives clusters containing at least two DEG vectors. For each set and cluster, we calculated the fraction of vectors in that cluster that belonged to that set (precision) and the fraction of vectors of that set that were in that cluster (recall). Both values allowed calculation of the F1 score (beta = 1). For each set, we defined the highest F1 score as the set-selective clustering efficiency and the related cluster the set-selective cluster. We additionally calculated a DEG-vector selective F1 score. This is the highest F1 score calculated for any cluster that contained the set-selective cluster and the DEG vector in question.

Since the 3 CT groups, NvsCT and PvsNP annotation grouped all 161 DEG vectors into only 3, 2 or 2 sets, respectively, we introduced an additional constraint. We only considered set-selective clusters that contained at most 60% of all DEG vectors.

### Identification of set-selective subspaces and gene expression profiles

To identify set-selective DEG vectors we used an updated version of our published pipeline (Suppl. Fig. 1B).^7^ Briefly, our published pipeline subjects the matrix consisting of the 161 complete DEG vectors to singular value decomposition (SVD). SVD factorizes a matrix into the product of a matrix of orthonormal left singular vectors, a diagonal matrix containing singular values, and a matrix of orthonormal right singular vectors. If applied to a matrix where rows represent genes and columns treatment/cell line combinations, the left singular vectors are often labeled eigenarrays,^13^ since each complete DEG vector is a linear combination of all eigenarrays. Vector-selective coefficients for this linear combination are given by the right singular vectors, while the singular values are additional coefficients identical for all vectors that document the initial variance explained by the related eigenarray. Using the vector-selective coefficients and a two-tailed t-test, our published pipeline tests each of the 161 eigenarrays for its ability to separate DEG responses of a set in question from all other DEG responses. For each set, all eigenarrays are ranked by significance, before generating 159 subspaces spanned by the top 3 to 161 eigenarrays. Within each subspace, our published pipeline calculated set- and DEG-vector selective F1 scores as described above. To document the amount of initial information that is still preserved in the identified subspace, we calculated the cosine similarities between the complete DEG vectors of that set and their projections into the identified subspace. To identify potential set-selective subspaces, we calculated the weighted average between the median cosine similarity and the set-selective F1 score, where we changed the weight of the F1 score (i.e., its relative contribution) from 0.00 to 0.95 in 0.05 steps. In consequence, we identified up to 20 potential set-selective subspaces for each set.

To identify outlier responses within these potential set-selective subspaces, we here updated our pipeline. For each potential set-selective subspace, we analyzed for each DEG vector that was not part of the set-selective clusters if its vector-selective clustering efficiency (F1 score) was significantly different from the vector-selective clustering efficiencies of the DEG vectors within the set-selective cluster. In addition to the already implemented Dixon’s test, we used Grubbs’ test if the number of supplied F1 scores exceeded the maximum number of 29 values that can be analyzed with Dixon’s test. If the identified vector-selective cluster contained more than 60% of all DEG vectors (our cutoff defined above), it was disqualified and its F1 score was set to zero before testing. Results were adjusted by Bonferroni correction. In addition to a significant adjusted P value (≤ 0.01), we introduced a second cutoff, i.e., that the related vector-selective cluster must contain at least 33% of all DEG vectors. This ensured that vectors that are close to set-selective clusters with high F1 scores are not labeled as outliers, which would contradict intuitive annotation after manual investigation of the cluster dendrogram. DEG vectors passing both cutoffs were labeled as outliers, all other DEG vectors of the set in question as regular responses.

To identify the set-selective subspace from the 20 potential subspaces we updated our published pipeline as well. We first considered only subspaces that gave a minimal set-selective clustering efficiency (F1 score) of 33%. If no subspace fulfilled this requirement, we selected the subspace with the highest set-selective clustering efficiency. Otherwise, we consecutively ranked the identified subspaces by the number of regular DEG responses, set-selective clustering efficiencies and decreasing F1 score weights. The top ranked subspace was defined as the set-selective subspace and the DEG vectors within that subspace that belonged to the set in question as the set-selective DEG vectors. Identified 161 set-selective DEG vectors of all sets were combined into a new matrix that was subjected to pairwise correlation analysis, followed by hierarchical clustering.

### Separation of regular and outlier responses and iterative application of the pipeline

If set-selective DEG vectors contained at least four outlier and regular responses that constituted at least 20% of all responses annotated to the same set, the outlier and regular responses were separated into two new sets. We selected the second cutoff, because in the 3 CT groups, NvsCT and PvsNP annotations all seven treatments of each cell line are part of the same initial set (i.e., no-, reversible, persistent, combined reversible and persistent or combined persistent and reversible CTRCD). Since the transcriptional response in the complete data is dominated by cell line-selectivity, our algorithm could group the DEG vectors into sets that only contain single or a few cell lines otherwise. In such a case, the identified set-selective DEGs would again be cell-line-selective. Each iteration contained the described calculation of the clustering efficiency for the cardiotoxicity groups. Our pipeline proceeded with the iterations until no set contained the required numbers of outlier and regular responses. Results of the last iteration were defined as final.

Note that our algorithm splits sets into new sets but does not split the CTRCD groups into new groups. In the CTRCD-group based initial set annotations (3 CT groups, NvsCT, PvsNP), the cardiotoxicity groups only overlap with the initial sets. At the end of each iteration, the sets can be split, while the CTRCD group annotations are unaffected.

### Calculation of the clustering efficiency for each cardiotoxicity group

Within the final matrix of merged set-selective DEG vectors, we calculated the clustering efficiency for each CTRCD group (no-, reversible, persistent, combined persistent and reversible and combined reversible and no-CTRCD) using the same algorithm described above. In the case of the T & 3 CT groups and T & PvsNP annotations, we only considered cell lines that were either treated with high-dose single or simultaneous or sequential regimens.

### Robustness analysis after cardiotoxicity label shuffling

In the case of the 3 CT group, NvsCT, PvsNP, T & 3 CT group and T & PvsNP annotations, we repeated the pipeline 100 times after shuffling of the cardiotoxicity labels. Each time, we updated the sets and calculated the cardiotoxicity group clustering efficiencies as described above. We calculated Monte Carlo P values by comparing the final clustering efficiencies for the cardiotoxicity groups without and with group label shuffling.

### Identification of marker genes with differential induction between any two cardiotoxicity groups

We compared the induced final set-selective expression levels of literature-curated serum marker genes^14–17^ between all treatments of persistent and non-persistent, reversible or no-CTRCD cell lines based on the PvsNP, 3 CT groups and T & PvsNP annotation, using the Wilcoxon rank-sum test. For the T & PvsNP annotation, we only considered high-dose single or simultaneous or sequential treatments, since we could resolve persistent and non-persistent CTRCD cell lines, if considering only those treatments. Genes with an adjusted P value (Benjamini-Hochberg correction) of at most 0.05 were labeled as significant.

### Identification of pathways with differential induction between any two cardiotoxicity groups

For the identification of cardiotoxicity group-selective pathways, we compared pathway expression between the persistent and the non-persistent, reversible or no-CTRCD groups using the PvsNP, 3 CT groups, and T&PvsNP annotations. We focused on these comparisons, since the persistent samples were clearly separated from those two groups, as indicated by either a significant empirical P value (PvsNP) or a trend towards significance (3 CT groups and T&PvsNP). For the T&PvsNP annotation, we only considered cell lines treated with high-dose single or simultaneous or sequential treatment combinations.

Up- and downregulated genes among the top 1000 genes of each final set-selective DEG vector were submitted to pathway enrichment analysis using MBC PathNet^18,19^ and the Molecular Biology of the Cell Ontologyv2 (MBCOv2).^19,20^ log_10_ P values of downregulated pathways were subtracted from those of upregulated pathways to calculate overall pathway changes. log_10_ P values of not identified pathways were set to 0 before this calculation. If a pathway was neither up- nor downregulated the difference was consequently zero as well. To focus on pathways with relevant expression changes, we subjected only pathways to the following analysis that were significantly (|log_10_ P value difference| ≥ log_10_ 0.05) induced in the same direction (i.e., up or down) in at least 10% of all cell line/treatment combinations annotated to at least one of the two compared cardiotoxicity groups. -log_10_ P value differences between the persistent and the non-persistent, reversible or no-CTRCD groups were then compared using the Wilcoxon rank-sum test.

MBCO v2 pathways are annotated at four different levels, where pathways of the same level show minimal redundancies and lower-level pathways (with higher level numbers) describe subfunctions of higher-level pathways. Consequently, we used the Benjamini-Hochberg correction separately for all pathways of the same level. Pathways with an adjusted Wilcoxon P value of at most 0.05 were considered overrepresented in the group with the higher mean value.

The identification of pathways was repeated using set-selective DEGs that our pipeline identified after removal of each of the 23 cell lines. For each pathway predicted from the complete dataset, we counted in how many downsampled datasets of the same set annotation the pathway was also predicted with significance.

We finally selected pathways for discussion and network visualization that were either predicted based on the PvsNP annotation and in at least 70% of the related downsampled datasets, or on the basis of two different annotations and in at least 50% of the related downsampled datasets for both annotations.

## RESULTS

### Patient cohort

We studied 23 women with HER2-positive breast cancer who received doxorubicin (approximately 250 mg/m²), cyclophosphamide with paclitaxel, followed by trastuzumab, all with normal cardiac function before treatment (Table 1, Fig. 1). Five patients maintained normal LVEF (61 ± 2%) throughout treatment and through twelve months of trastuzumab therapy (no CTRCD). Eighteen patients developed CTRCD, with LVEF falling to 30 ± 9% from a baseline of 61 ± 5% (P < 0.001). Of these, eleven recovered LVEF ≥ 53% within a median of 10.4 months (IQR: 6.3 to 22.1 months) (reversible CTRCD), whereas seven did not recover over a median follow-up of 71.2 months (IQR: 53.8 to 92.4 months) (persistent CTRCD). iPSC-derived cardiomyocytes from each patient were exposed to seven drug regimens: (1) doxorubicin for 24 h, (2,3) simultaneous treatment with doxorubicin and low- or high-dose trastuzumab for 48 h, (4,5) 24 h doxorubicin followed by addition of low- or high-dose trastuzumab for another 48 h and (6,7) low- or high-dose trastuzumab alone for 48 h. We selected these time periods to ensure equal exposure to trastuzumab for 48 h within each treatment regimen.

**Figure 1.**
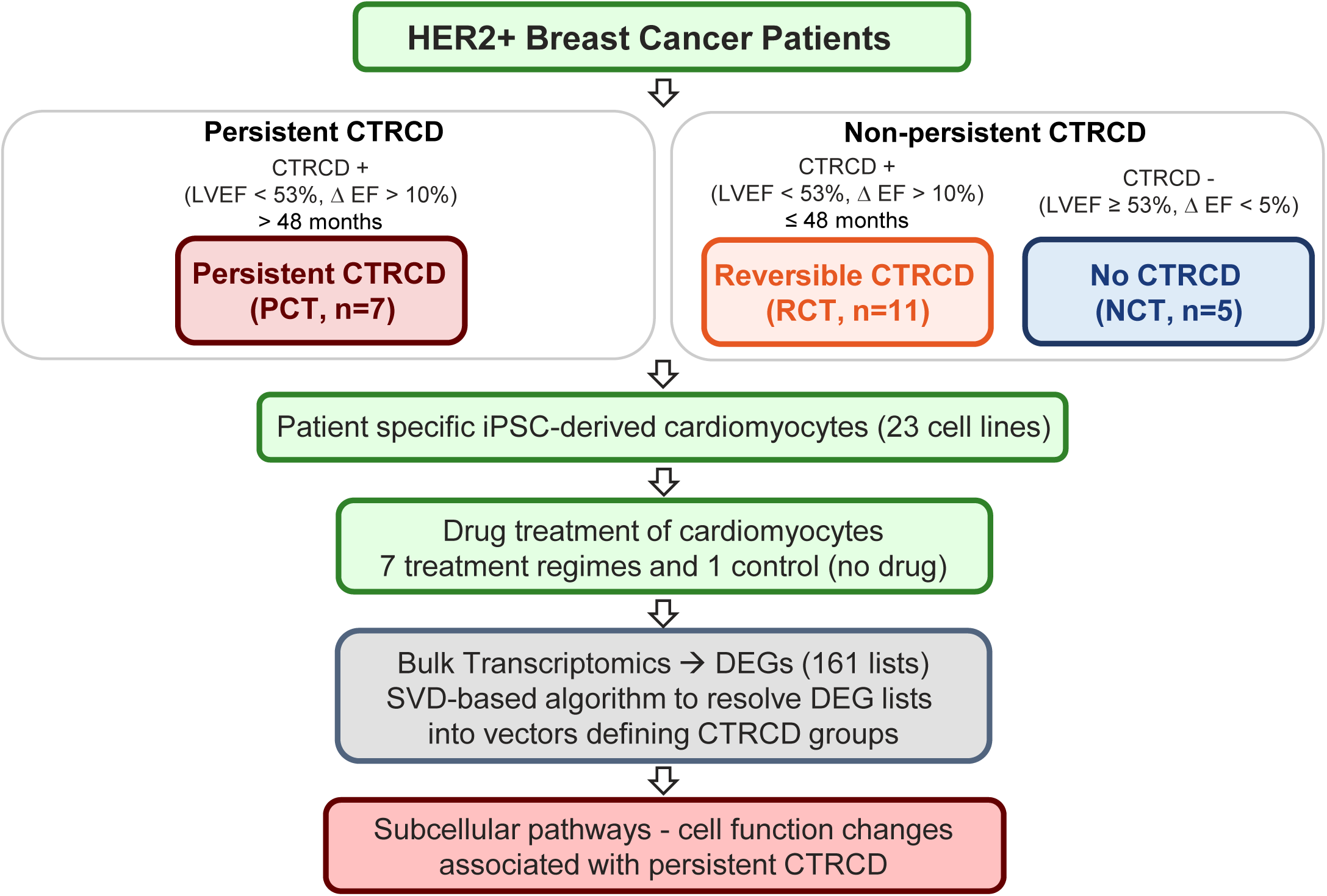
Study design and patient groups. Patients were grouped into the persistent CTRCD, reversible CTRCD and no-CTRCD groups based on indicated left ventricular ejection fraction (LVEF) criteria. Patient-specific induced pluripotent stem cells (iPSCs) derived cardiomyocytes were exposed to seven doxorubicin and trastuzumab regimens, followed by transcriptomic profiling and computational resolution of clinical group-selective responses. CTRCD = cancer therapy-related cardiac dysfunction; DEGs = differentially expressed genes; EF = ejection fraction; NCT = no CTRCD; PCT = persistent CTRCD; RCT = reversible CTRCD; SVD = singular value decomposition.

**TABLE 1.**
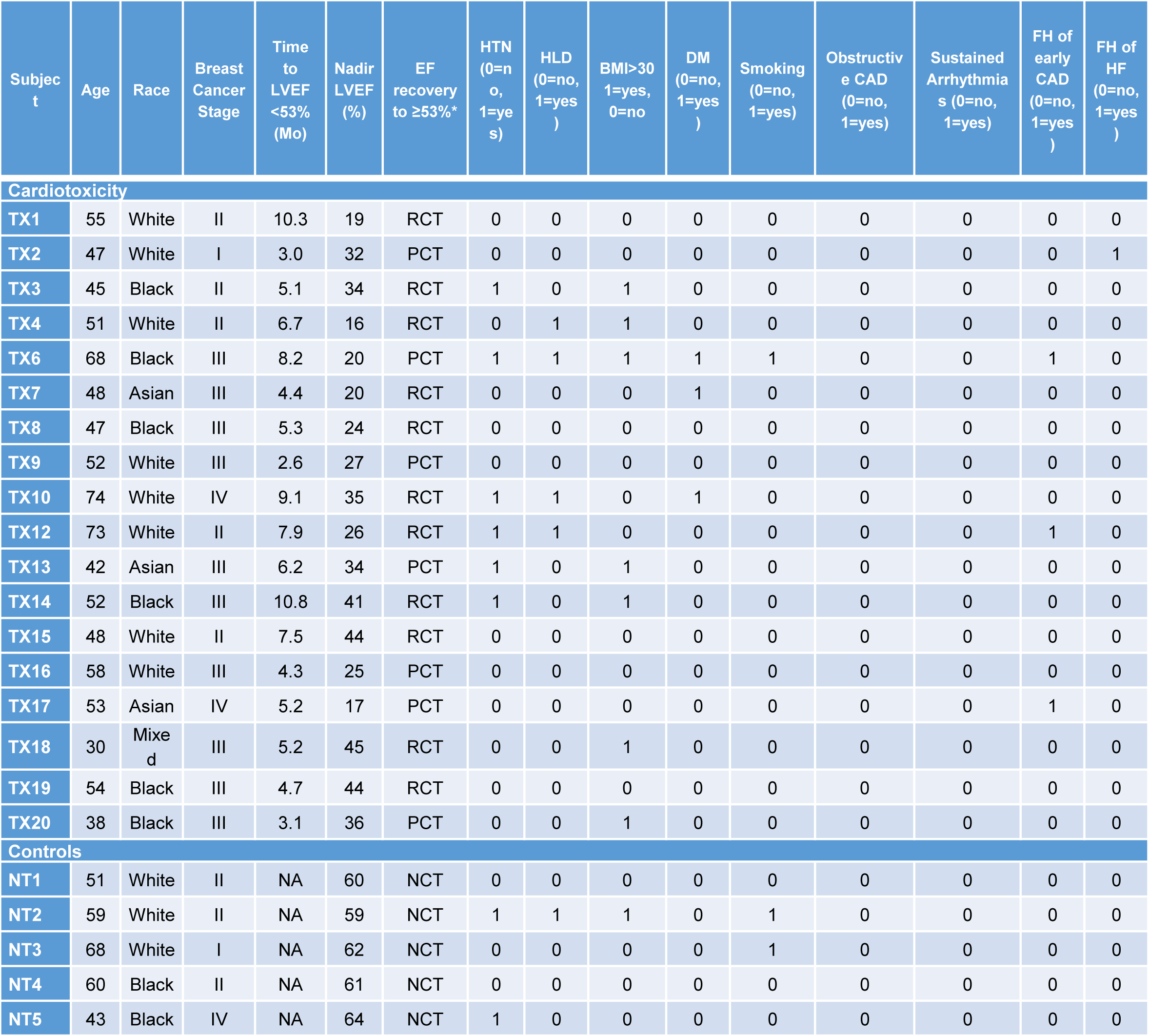
Characteristics of patients with HER2-positive breast cancer. Values are n (%), mean ± SD or median (IQR). BMI = body mass index; CAD = coronary artery disease; CTRCD = cancer therapy-related cardiac dysfunction; HLD = hyperlipidemia; HTN = hypertension; LVEF = left ventricular ejection fraction; other abbreviations as in Figure 1.

| **Subject** | **Age** | **Race** | **Breast Cancer Stage** | **Time to LVEF <53% (Mo)** | **Nadir LVEF (%)** | **EF recovery to ≥53%\*** | **HTN (0=no, 1=yes)** | **HLD (0=no, 1=yes)** | **BMI>30 1=yes, 0=no** | **DM (0=no, 1=yes)** | **Smoking (0=no, 1=yes)** | **Obstructive CAD (0=no, 1=yes)** | **Sustained Arrhythmias (0=no, 1=yes)** | **FH of early CAD (0=no, 1=yes)** | **FH of HF (0=no, 1=yes)** |
|---|---|---|---|---|---|---|---|---|---|---|---|---|---|---|---|
| **Cardiotoxicity** |  |  |  |  |  |  |  |  |  |  |  |  |  |  |  |
| **TX1** | 55 | White | II | 10.3 | 19 | RCT | 0 | 0 | 0 | 0 | 0 | 0 | 0 | 0 | 0 |
| **TX2** | 47 | White | I | 3.0 | 32 | PCT | 0 | 0 | 0 | 0 | 0 | 0 | 0 | 0 | 1 |
| **TX3** | 45 | Black | II | 5.1 | 34 | RCT | 1 | 0 | 1 | 0 | 0 | 0 | 0 | 0 | 0 |
| **TX4** | 51 | White | II | 6.7 | 16 | RCT | 0 | 1 | 1 | 0 | 0 | 0 | 0 | 0 | 0 |
| **TX6** | 68 | Black | III | 8.2 | 20 | PCT | 1 | 1 | 1 | 1 | 1 | 0 | 0 | 1 | 0 |
| **TX7** | 48 | Asian | III | 4.4 | 20 | RCT | 0 | 0 | 0 | 1 | 0 | 0 | 0 | 0 | 0 |
| **TX8** | 47 | Black | III | 5.3 | 24 | RCT | 0 | 0 | 0 | 0 | 0 | 0 | 0 | 0 | 0 |
| **TX9** | 52 | White | III | 2.6 | 27 | PCT | 0 | 0 | 0 | 0 | 0 | 0 | 0 | 0 | 0 |
| **TX10** | 74 | White | IV | 9.1 | 35 | RCT | 1 | 1 | 0 | 1 | 0 | 0 | 0 | 0 | 0 |
| **TX12** | 73 | White | II | 7.9 | 26 | RCT | 1 | 1 | 0 | 0 | 0 | 0 | 0 | 1 | 0 |
| **TX13** | 42 | Asian | III | 6.2 | 34 | PCT | 1 | 0 | 1 | 0 | 0 | 0 | 0 | 0 | 0 |
| **TX14** | 52 | Black | III | 10.8 | 41 | RCT | 1 | 0 | 1 | 0 | 0 | 0 | 0 | 0 | 0 |
| **TX15** | 48 | White | II | 7.5 | 44 | RCT | 0 | 0 | 0 | 0 | 0 | 0 | 0 | 0 | 0 |
| **TX16** | 58 | White | III | 4.3 | 25 | PCT | 0 | 0 | 0 | 0 | 0 | 0 | 0 | 0 | 0 |
| **TX17** | 53 | Asian | IV | 5.2 | 17 | PCT | 0 | 0 | 0 | 0 | 0 | 0 | 0 | 1 | 0 |
| **TX18** | 30 | Mixed | III | 5.2 | 45 | RCT | 0 | 0 | 1 | 0 | 0 | 0 | 0 | 0 | 0 |
| **TX19** | 54 | Black | III | 4.7 | 44 | RCT | 0 | 0 | 0 | 0 | 0 | 0 | 0 | 0 | 0 |
| **TX20** | 38 | Black | III | 3.1 | 36 | PCT | 0 | 0 | 1 | 0 | 0 | 0 | 0 | 0 | 0 |
| **Controls** |  |  |  |  |  |  |  |  |  |  |  |  |  |  |  |
| **NT1** | 51 | White | II | NA | 60 | NCT | 0 | 0 | 0 | 0 | 0 | 0 | 0 | 0 | 0 |
| **NT2** | 59 | White | II | NA | 59 | NCT | 1 | 1 | 1 | 0 | 1 | 0 | 0 | 0 | 0 |
| **NT3** | 68 | White | I | NA | 62 | NCT | 0 | 0 | 0 | 0 | 1 | 0 | 0 | 0 | 0 |
| **NT4** | 60 | Black | II | NA | 61 | NCT | 0 | 0 | 0 | 0 | 0 | 0 | 0 | 0 | 0 |
| **NT5** | 43 | Black | IV | NA | 64 | NCT | 1 | 0 | 0 | 0 | 0 | 0 | 0 | 0 | 0 |

### Cardiomyocyte responses identify patients with persistent CTRCD

Unprocessed transcriptomic responses to drug treatments clustered predominantly by cell line rather than by clinical group (Fig. 2A, Suppl. Fig. 1A, Suppl. Table 1), indicating that the identity of the individual patient’s cardiomyocytes dominates the drug response, as we had observed in healthy donor lines before.^7^ Clinical-outcome-related signals are therefore superimposed on a much larger patient-specific response and have to be resolved from it, before any clinical comparison can be made.

**Figure 2.**
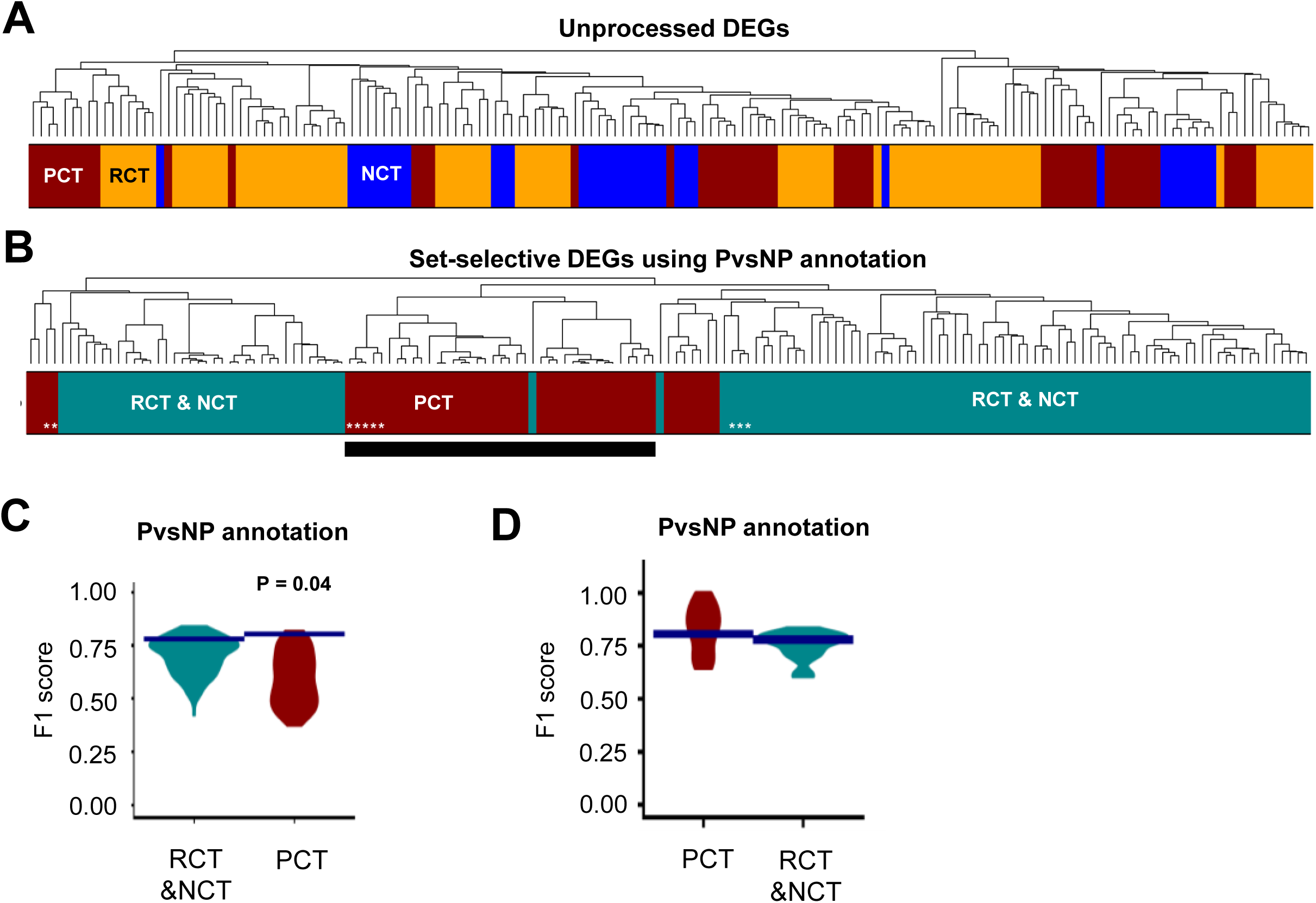
Responses resolve patients with persistent CTRCD. **(A)** Unprocessed response profiles cluster by cell lines rather than CTRCD groups. **(B)** The same analysis after SVD-based resolution using the PvsNP set definition generates a core cluster of persistent CTRCD responses (black bar; white asterisks indicate transcriptomic outlier responses in the set-selective subspaces). **(C)** Clustering efficiencies (F1 scores) of each CTRCD group in the true data (blue lines) and after 100 label permutations through the full pipeline (violins) with the empirical P value. **(D)** Clustering efficiencies in the complete data (blue lines) compared with those obtained in the 23 leave-one-patient-out datasets (violin). PvsNP = persistent versus non-persistent CTRCD; other abbreviations as in Figure 1.

We applied our updated SVD-based pipeline for the identification of set-selective response profiles (Suppl. Fig. 1B). For six independent analyses, we defined six different set annotations (Suppl. Table 2) that grouped DEG vectors induced by the same treatment in any cell line (1 in Suppl. Fig. 1C) and induced by any or the same treatment in cell lines with matching CTRCD responses (2,3,4 or 5,6, respectively, in Suppl. Fig. 1C). For technical results obtained during these analyses, see the Results section in the Supplemental Appendix.

After SVD-based resolution, responses induced in persistent CTRCD lines enriched within a single cluster with low merging heights (Fig. 2B). This was observed whenever the algorithm actively searched persistent CTRCD-selective responses, i.e., within the persistent versus non-persistent comparison (PvsNP) (Fig. 2B, Suppl. Fig. 2A) and within the comparison that kept all three clinical groups separately (3 CT groups) (Suppl. Fig. 2B). When additionally stratifying responses within the PvsNP annotation by treatment regimen (T & PvsNP), our algorithm was still able to resolve persistent CTRCD-selective responses, at least those induced by sequential or high-dose regimens (Suppl. Fig. 2C). Comparing responses in cell lines with no CTRCD versus CTRCD (NvsCT), we observed a cluster enriched for cell lines without CTRCD (Suppl. Fig. 2D). The search for treatment-selective responses across all cell lines (T) (Suppl. Fig. 2E) and across cell lines with the same CTRCD (T & 3 CT groups) (Suppl. Fig. 2F) did not sufficiently resolve any CTRCD group.

Shuffling of the clinical labels 100 times through the full pipeline showed that close clustering of persistent CTRCD lines exceeded chance for PvsNP comparison (P = 0.04; minimum possible p ≈ 0.01) (Fig. 2C). Concordant results were obtained for the 3 CT groups (P = 0.069) and the T & PvsNP comparisons (P = 0.089) (Suppl. Fig. 3A). A tendency toward significant enrichment of no-CTRCD cell lines within the same cluster was further documented for the NvsCT comparison (P = 0.079). No significance was documented for the T & 3 CT groups comparison.

Since the empirical P values lie near, rather than far below, conventional thresholds, we interpret them as convergent evidence for a reproducible grouping rather than individually decisive tests. Label shuffling with identical group sizes and pipeline indicates that the algorithm does not simply extract shared components from any arbitrary set.

Significance or a tendency toward significance was observed for both annotations that grouped all persistent CTRCD lines into one set (i.e., PvsNP, 3 CT groups). Additionally, we observed a tendency for significance for persistent CTRCD lines (treated with high-dose or sequential regimens) after additional stratification of the PvsNP sets by treatment regimen (T & PvsNP), where the algorithm does not enforce similarity across different treatments within the persistent CTRCD lines. As negative controls, apart from the tendency for no-CTRCD cell lines in the NvsCT comparison, we did not document empirically significant enrichment for the reversible or no CTRCD groups in any other annotation.

In leave-one-patient-out analysis, clustering efficiencies of persistent CTRCD responses were similar to those obtained with the complete dataset (Fig. 2D, Suppl. Fig. 3B) and the pathway-level differences reported below were identified in most downsampled datasets (Suppl. Figs. 7 and 8).

Taken together, these analyses establish that responses of cardiomyocytes from patients with persistent CTRCD form a reproducible group that is not an artifact of the resolution procedure or of individual patients. They do not establish that the transcriptomic response alone would classify an individual patient prospectively, which would require an independent validation cohort. Subsequent comparisons are presented as descriptions of group-level potential physiological responses.

### Genes encoding guideline-recommended cardiac biomarkers

We next examined genes coding for established and suggested circulating markers of dilated, hypertrophic cardiomyopathy ^14,15^ and general cardiac injury ^17^ (Fig. 3A, Suppl. Fig. 4). Of 44 genes encoding 40 circulating markers, 30 were differentially induced between persistent CTRCD lines and the comparison groups (Fig. 3B). For 17 of these, including the genes encoding the biomarkers recommended for CTRCD surveillance by the 2022 European Society of Cardiology cardio-oncology guideline (natriuretic peptides: NPPA, NPPB, cardiac troponins: TNNT2, TNNI3 and cardiac/skeletal troponin C: TNNC1),^6^ the direction of induction in cardiomyocytes matched the direction of the reported clinical association: genes whose circulating products rise with increased risk were more strongly induced in persistent CTRCD lines. For 10 genes, including those encoding myoglobin, endostatin, big endothelin-1, GDF15 and apelin, the relationship was reversed, and results were discordant across annotations for 3 genes (TGFB1, TIMP1 and VEGFA). Transcript levels may reflect protein abundance or compensatory mechanisms that counteract it and circulating marker levels also depend on leakage rates, which can increase with cellular damage even when intracellular protein abundance stays constant. This re-identification of potential and recommended clinical circulating risk markers provides external validation of the gene expression profiles identified by our algorithm.

**Figure 3.**
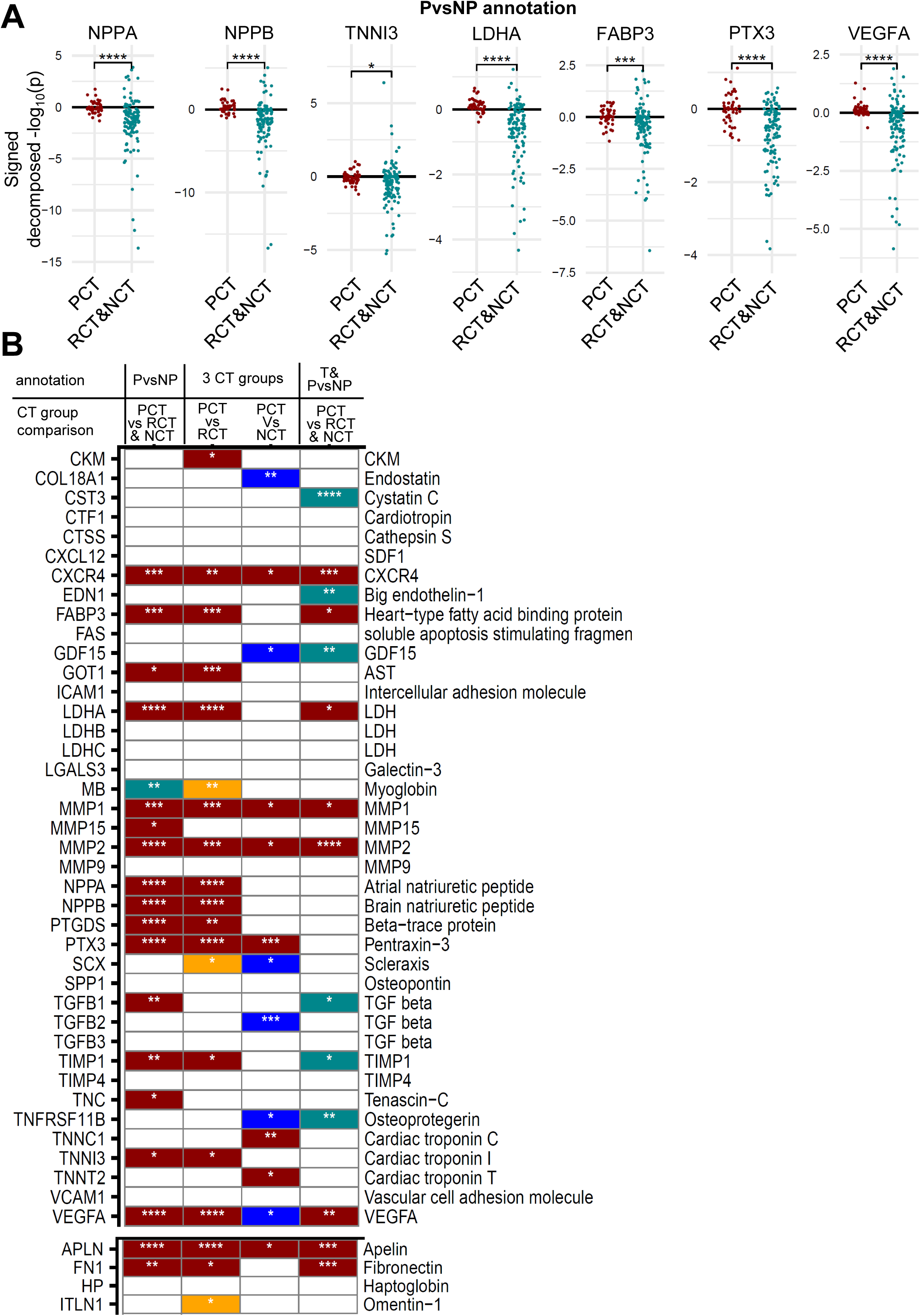
Induction of genes encoding circulating cardiac biomarkers. **(A)** Set-selective induction of representative marker genes in persistent versus non-persistent CTRCD cardiomyocytes. **(B)** Literature-curated marker genes (left labels) for cardiomyopathy and cardiac injury, grouped by whether higher (top) or lower (bottom) circulating levels of the encoded proteins (right labels) are associated with increased risk. *, **, ***, ****: adjusted Wilcoxon P ≤ 0.05, ≤ 0.01, ≤ 0.001, ≤ 0.0001. Colors indicate the group with higher expression: dark red: persistent CTRCD, turquoise: non-persistent CTRCD, orange: reversible CTRCD, blue: no-CTRCD. 3 CT groups = 3 CTRCD groups annotations, T & PvsNP: PvsNP annotation stratified by treatment annotations, respectively. Other abbreviations as in Figures 1 and 2.

### Contractile, calcium and cytoskeletal pathways separate persistent CTRCD

To predict functional responses selective for persistent CTRCD lines (Suppl. Fig. 5A), we subjected set-selective response profiles to pathway enrichment analysis with the Molecular Biology of the Cell Ontology version 2 ^19,21^ (Suppl. Fig. 5B). Using the PvsNP, the 3 CT groups and the T & PvsNP comparisons, we identified pathway activity changes selective for cell lines with persistent CTRCD (Fig. 4A, Suppl. Fig. 6A to C, respectively). Multiple identified differences mapped to cellular functions that converge on contractile homeostasis, when comparing responses in persistent CTRCD lines against all lines, reversible CTRCD (Fig. 4B, Suppl. Fig. 7) and no CTRCD lines (Suppl. Fig. 8). Predictions were concordant across hierarchical parent-child levels for the former (Fig. 4C, Supp. Fig. 9) and latter comparisons (Suppl. Fig. 10).

**Figure 4.**
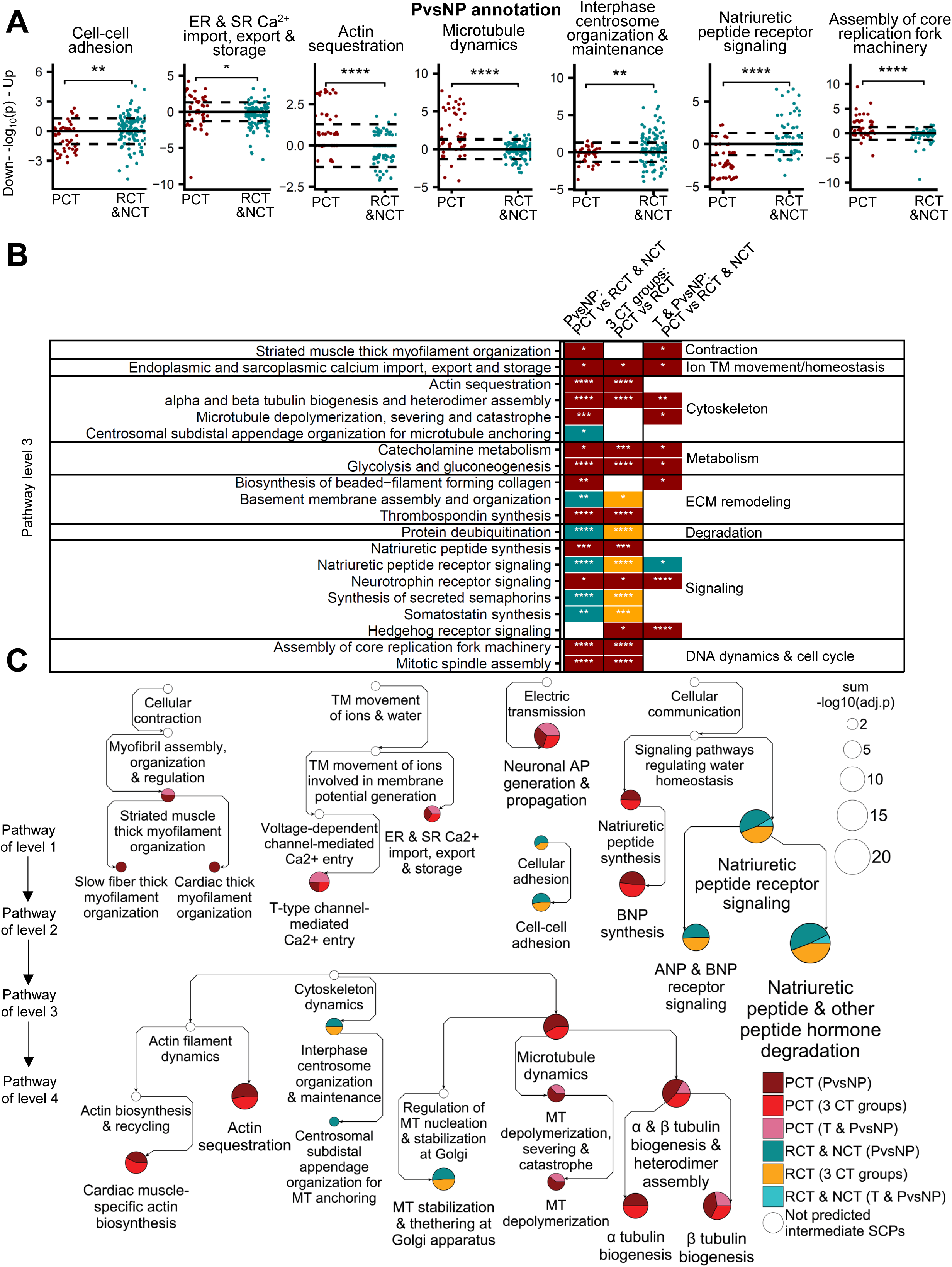
Cellular pathways distinguishing persistent CTRCD. **(A)** Net induction of representative pathways in each cell line and treatment combination. **(B)** Consistently predicted MBCO v2 level 3 pathways with different net induction grouped by cellular function. Colors indicate the group with higher induction (PCT: dark red; RCT and NCT: turquoise; RCT: orange). **(C)** Selected pathways mapped to the MBCO parent-child hierarchy. Arrows connect parent and child pathways. Pie slice areas are proportional to the summed -log_10_(adjusted P value) of all shown conditions. MBCO = Molecular Biology of the Cell Ontology; v = version; other abbreviations as in Figures 1 to 3.

Pathways describing striated muscle and cardiac thick myofilament organization, action potential generation, endoplasmic and sarcoplasmic calcium import, export and storage, and T-type channel-mediated calcium entry were more strongly induced in persistent CTRCD cardiomyocytes. Cytoskeletal pathways diverged: persistent CTRCD lines showed increased tubulin heterodimer supply, microtubule depolymerization, and cardiac actin biosynthesis and sequestration, whereas reversible lines showed increased centrosomal and Golgi-associated microtubule anchoring, functions that can maintain the ordered microtubule architecture of the cardiomyocyte.^22^ Increased free actin and tubulin monomer supply with increased depolymerization could be the profile of a cytoskeleton undergoing remodeling, and an increase in microtubule density is an established contributor to increased myocyte stiffness in human heart failure.^23^ Cell-cell adhesion pathways and basement membrane assembly,^24^ both of which support mechanical coupling and mechanotransduction, were higher in reversible and no-CTRCD lines.

### Pathways reflecting doxorubicin’s mechanism of action

Pathways reflecting doxorubicin’s mechanism of action, i.e., topoisomerase II inhibition and induction of double-strand breaks ^5^ are upregulated to higher levels in persistent CTRCD lines. Among these are replication fork assembly, homologous recombination-mediated DNA double-strand break repair, and interstrand crosslink repair, the latter comprising Fanconi anemia genes. Besides their initially identified function,^25^ some of the Fanconi anemia genes can support replication fork protection,^26^ replication-stress response ^27^ and homologous recombination-mediated double-strand break repair.^28^

### Natriuretic peptide resistance

Natriuretic peptide receptor signaling, and its child pathways ANP and BNP receptor signaling and Natriuretic peptide and other peptide hormone degradation, were consistently lower in persistent CTRCD lines (Fig. 4B, Suppl. Figs. 7C, D). The pathway (and its children) reached significance in the complete dataset and in 83% (65% and 96%) and 100% (100% and 100%) of the downsampled datasets, if compared with all other or the reversible CTRCD cell lines, respectively (Suppl. Figs. 7C, D).

While natriuretic peptide receptor signaling was down- and upregulated in most persistent and non-persistent CTRCD lines, respectively (Fig. 4A, Suppl. Fig. 6A, B, C), pathways and genes indicating ligand synthesis displayed the opposite relationship. NPPA and NPPB were downregulated in most non-persistent CTRCD lines and showed inconsistent expression changes in persistent CTRCD lines (Fig. 3A, Suppl. Fig. 4A, B, C, D). Our results could reflect a cardiomyocyte manifestation of the multi-organ natriuretic peptide resistance classically described in chronic heart failure ^29,30^ and are consistent with earlier observations in the murine heart.^31^

### Glycolysis

Glycolysis and gluconeogenesis were among the pathways most strongly and reproducibly associated with persistent CTRCD (Fig. 4, Suppl. Fig. 7-10).

### Extracellular matrix dynamics

Thrombospondin family members that are induced in pressure overload models ^32^ and collagen II, VI and XXIV syntheses were predicted at higher levels for the persistent CTRCD lines (Suppl. Figs. 7-10).

### Convergence with the cardiotoxic kinase inhibitor signature

To determine whether identified pathways represent more general mechanisms of drug-induced cardiotoxicity, we mapped them onto pathway signatures that distinguish cardiotoxic from non-cardiotoxic TKIs in cardiomyocytes from healthy donors,^7^ recomputed here with MBCO v2 (Fig. 5, Suppl. Fig. 11).

**Figure 5.**
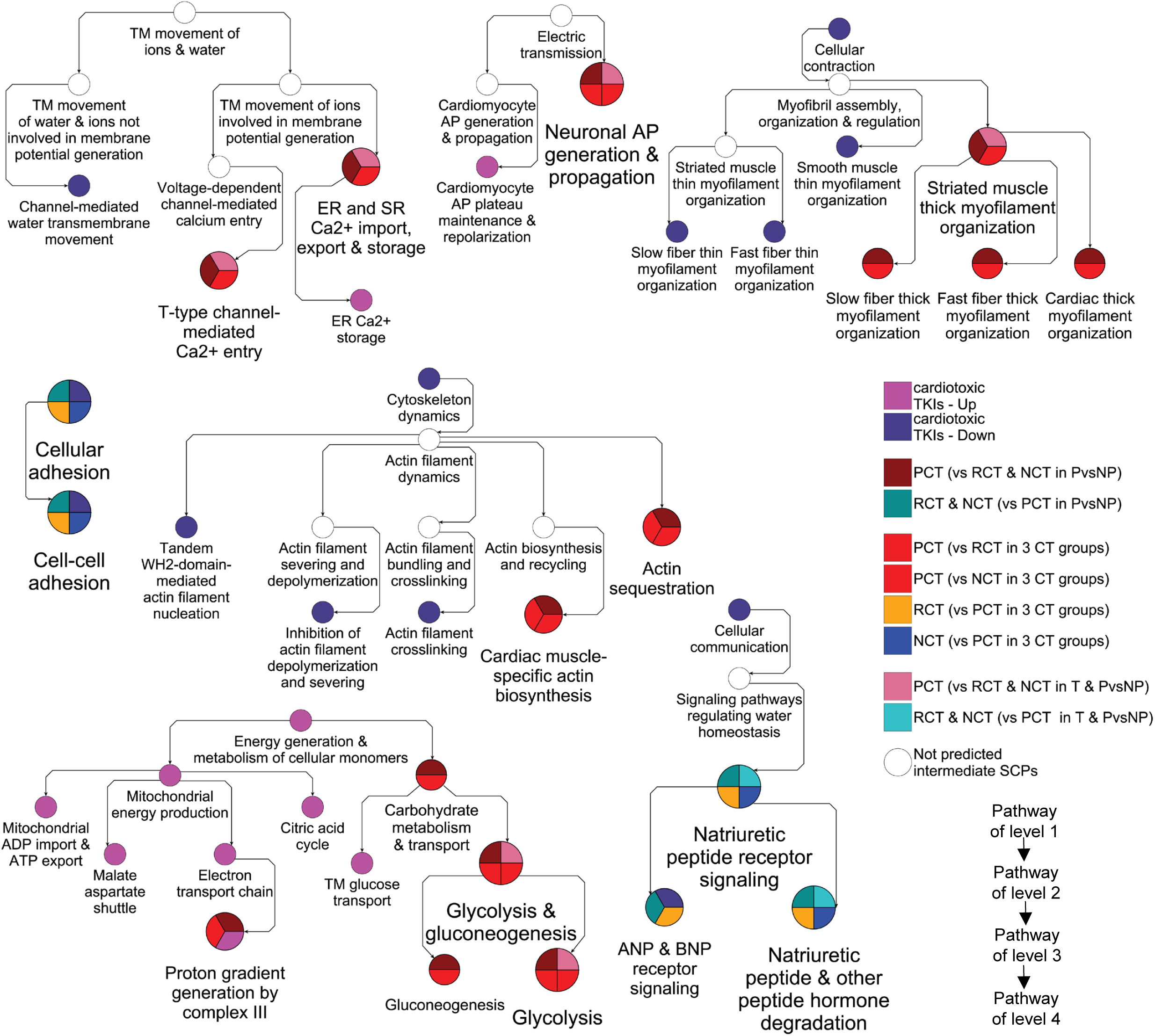
Cardiotoxic kinase inhibitor signature pathways. Pathways differentially induced by cardiotoxic relative to non-cardiotoxic tyrosine kinase inhibitors (TKIs) in healthy donor cardiomyocytes, were mapped onto pathways identified in this study within the MBCO v2 hierarchy. Selected branches are shown. Abbreviations as in Figures 1 to 4.

Overlapping and taxonomically related pathways (Suppl. Fig. 12) responded either in the same direction (i.e., with higher or lower induction in the more toxic condition) or in opposite directions. Pathways responding in the same direction include electrical transmission and calcium homeostasis, cellular adhesion, mitochondrial energy generation, natriuretic peptide receptor signaling, glucose metabolism and transport, and cell cycle dynamics. Pathways responding in opposite directions map to cellular contraction, DNA replication and actin dynamics. Notably, the actin pathways that increase in persistent CTRCD lines describe monomer supply and sequestration, whereas those that decrease after cardiotoxic TKI exposure describe nucleation, crosslinking and inhibition of depolymerization. Cardiotoxic TKIs additionally induced a broader set of lipid, fatty acid and amino acid metabolic pathways not identified in our current study. Recurrent identification of contractile, calcium-handling and natriuretic peptide receptor signaling pathways across two independent cohorts, drug classes and study designs, despite opposite-direction changes in the contraction pathway, supports disrupted contractile homeostasis and natriuretic peptide signaling as cellular functions central to drug-induced cardiac dysfunction.

## DISCUSSION

We documented different responses to anthracycline and trastuzumab treatment in cardiomyocytes derived from patients who developed persistent CTRCD after therapy with those same drugs and in cardiomyocytes derived from patients who recovered or who never developed dysfunction (Central illustration). Our results suggest that susceptibility to persistent dysfunction is at least in part a cell-intrinsic property of the patient’s cardiomyocytes rather than solely a consequence of cumulative exposure or comorbidity. Three features of the identified signatures have potential clinical implications.

### Contractile homeostasis

Multiple pathways that consistently separate persistent CTRCD cardiomyocytes from the other groups describe cellular functions that converge on contraction, i.e., force generation, calcium handling, energy supply, mechanical coupling, actin dynamics and the microtubule system that determines passive stiffness.^23^ The same functional categories distinguished cardiotoxic TKIs, suggesting that disturbed contractile homeostasis is a shared cellular feature of drug-induced cardiac dysfunction.

### Natriuretic peptide resistance

Monitoring natriuretic peptides and troponins is recommended for CTRCD screening. The corresponding genes were downregulated in most cardiomyocytes from patients with reversible or no CTRCD and displayed inconsistent changes in those with persistent CTRCD, providing cellular-level support for this clinical practice. We additionally found natriuretic peptide receptor signaling as well as the peptidase responsible for natriuretic peptide degradation to be upregulated in cardiomyocytes from patients with reversible or no CTRCD and downregulated in those with persistent CTRCD. Sacubitril-valsartan acts in part by raising the availability of natriuretic peptides through inhibition of their degradation ^33^ and has shown benefit in heart failure ^34^ and, in early studies, in cancer therapy-related dysfunction.^35^ If cardiomyocytes in patients at highest risk exhibit natriuretic peptide insensitivity due to receptor downregulation, increasing ligand levels alone may be insufficient to restore beneficial signaling effects in this patient subgroup. Mechanistically, the observed receptor downregulation could be the consequence of chronically elevated ligands or an intrinsic property within susceptible cardiomyocytes. In the latter case, natriuretic peptide receptor expression, rather than circulating peptide levels, would be a more relevant biomarker. Future experiments could investigate this question and potentially allow definition of patient subgroups for selective cardioprotection.

### Glucose handling

Increased glycolytic pathway activity in persistent CTRCD cardiomyocytes and increased glucose transmembrane transport induced by cardiotoxic TKIs are consistent with reports of increased myocardial glucose uptake during cardiotoxic therapy,^36^ although this association has not been confirmed uniformly.^37^ SGLT1 is a sodium-glucose transporter that is closely related to SGLT2, the target of a drug class with established heart failure benefit ^38^ and growing interest in cardio-oncology.^39,40^ While SGLT2 is not detected in healthy or diseased human heart tissue,^41^ SGLT1 is overexpressed in diseased human myocardium.^42^ Our finding provides a cardiomyocyte-level, patient-stratified rationale for evaluating dual sodium-glucose cotransporter inhibition ^43,44^ specifically in patients at risk for persistent rather than reversible dysfunction.

### Study limitations

Our study has several limitations. The number of patients in each group is small, particularly in the group without CTRCD, which comprised only 5 patients. Being unable to resolve reversible from no-CTRCD cell lines, we cannot state whether these two groups show generally similar responses at the cellular level or are simply underpowered. The empirical P values supporting the persistent CTRCD grouping are close to conventional thresholds, although they are reproducible across three different annotations and are accompanied by internal negative controls. Nevertheless, they are derived from seven patients and require confirmation in an independent cohort. All donors were women, as determined by the breast cancer population studied, so sex-related differences could not be assessed. Patients also received cyclophosphamide and paclitaxel, which were not included in our in-vitro regimes. We measured transcriptional responses to each treatment regimen at a single time point and therefore did not distinguish between transient and sustained transcriptional responses. Identified gene expression signatures need functional validation, e.g., calcium transient, force or stiffness measurements. Additionally, iPSC-derived cardiomyocytes are immature relative to adult myocardium and our experimental design lacks other heart cells, e.g., fibroblasts, endothelial and immune cells, as well as three-dimensional tissue structure and mechanical forces.

However, cardiomyocytes are the major cell type underlying cardiac contractility and our cardiomyocyte cultures exhibited robust, spontaneous contractile activity, suggesting suitability for studying contractility-related phenotypes.

## CONCLUSIONS

We documented a distinct, reproducible response to anthracycline and trastuzumab treatment in cardiomyocytes from patients who developed persistent CTRCD after therapy with the same drugs. Our findings suggest that susceptibility to irreversible cardiac dysfunction is, at least in part, a cell-autonomous property of the patient’s cardiomyocytes, characterized by disturbed contractile homeostasis and cardiomyocyte natriuretic peptide resistance. Natriuretic peptide receptor signaling and glucose handling could help define patient subgroups for selective cardioprotective intervention, allowing treatment strategies to be adapted to patient-selective biology.

## ABBREVIATIONS AND ACRONYMS

3 CT groups: 3 CTRCD groups
CTRCD: cancer therapy-related cardiac dysfunction
DEG: differentially expressed gene
iPSC-CM: induced pluripotent stem cell-derived cardiomyocyte
LVEF: left ventricular ejection fraction
MBCO: Molecular Biology of the Cell Ontology
NCT: no-CTRCD group
NvsCT: no-CTRCD vs CTRCD
PCT: persistent CTRCD group
PvsNP: persistent CTRCD vs non-persistent
CTRCD RCT: reversible CTRCD group
SGLT: sodium-glucose cotransporter
SVD: singular value decomposition
T: treatment
T & 3 CT groups: treatment & 3 CTRCD groups
T & PvsNP: treatment & persistent CTRCD vs non-persistent
CTRCD TKI: tyrosine kinase inhibitor

## PERSPECTIVES

### COMPETENCY IN MEDICAL KNOWLEDGE

Patients who develop cancer therapy-related cardiac dysfunction (CTRCD) during anthracycline and trastuzumab therapy and never recover to normal ejection fractions have cardiomyocytes that show different early responses to these drugs than cardiomyocytes from patients who recover or do not develop CTRCD. Our results indicate that persistence of dysfunction is partly determined by intrinsic properties of the patient’s cardiomyocytes rather than by cumulative exposure and comorbidities alone.

### TRANSLATIONAL OUTLOOK

An independent validation cohort and functional validation are needed before clinical translation. Prospective studies should test whether natriuretic peptide receptor signaling, contractile and glycolytic activity, measured in patient-derived iPSC-CMs that require only a blood sample, identify patients destined for persistent dysfunction can be used to select patients for targeted angiotensin receptor-neprilysin inhibition or sodium-glucose cotransporter inhibition.

## ACKNOWLEDGMENTS

The authors thank Gomathi Jayaraman, Debbie Certa, Tina Bin Hu and Som Prabha for assistance with cell culture, drug treatment and sample preparation; Drs Richard Steingart, Jennifer Liu and Anthony Yu for assistance with patient recruitment; and the staff of the Mount Sinai Genomics Core for library preparation and sequencing.

## AUTHOR CONTRIBUTIONS

Drs Chan and Iyengar conceived the study and obtained funding. Dr Chan recruited patients and designed and conducted clinical and experimental work. Dr Hansen performed all computational analyses. Dr Iyengar provided overall project management and has overall responsibility for the project. All authors wrote and revised the manuscript.

## DECLERATION OF GENERATIVE AI AND AI-ASSISTED TECHNOLOGIES IN THE MANUSCRIPT PREPARTION PROCESS

The manuscript was initially written and edited by all three authors. During the preparation of this work, we used Claude (Opus 5 and Opus 5.5, Anthropic) to correct typos, grammar and style to optimize the flow. After using this tool, the authors rechecked and rewrote the content as needed. Claude’s suggestion for the central illustration was slightly modified.

Code for preparation of pathway enrichment results for follow-up analyses and subtraction of down-from upregulated pathway activities was optimized for faster execution using Claude. Results obtained for code written by the authors and optimized by Claude were compared by the authors.

The authors take full responsibility for all content generated by AI.

## CODE AVAILABILITY

Analysis code has been deposited at github.com/DToxS/SVD_Cardiomyocyte_Signatures_of_Persistent_CTRCD/.

## SUPPLEMENTARY TABLES

**Supplementary Table 1**. Unprocessed differentially expressed genes.

**Supplementary Table 2**. Set annotations.

**Supplementary Table 3**. Final set-selective DEGs for the (A) PvsNP, (B) 3 CT groups, (C) T & PvsNP, (D) NvsCT, (E) T and (F) T & 3 CT groups set annotations.

**Supplementary Table 4**. Initial and final cluster efficiencies for cardiotoxicity groups within the (A) PvsNP, (B) 3 CT groups, (C) T & PvsNP, (D) NvsCT, (E) T and (F) T & 3 CT groups set annotations.

**Supplementary Table 5**. Comparative expression of clinical cardiotoxicity markers within the (A) PvsNP, (B) 3 CT groups, (C) T & PvsNP set annotations.

**Supplementary Table 6**. Pathway enrichment analysis of final DEGs for the (A) PvsNP, (B) 3 CT groups and (C) T & PvsNP annotations.

**Supplementary Table 7**. Pathways with significantly different induction levels between (A) PCT and RCT&NCT cell lines based on the PvsNP annotation, (B) PCT and RCT and (C) PCT and NCT based on the 3CT groups annotation as well as PCT and RCT&NCT based on the T&PvsNP annotation.

**Supplementary Table 8**. Pathway enrichment analysis of DEGs induced in healthy cardiomyocytes by cardiotoxic and non-cardiotoxic TKIs.

**Supplementary Table 9**. Pathways with significantly different induction levels in healthy cardiomyocytes after treatment with cardiotoxic and non-cardiotoxic TKIs.

**Supplementary Table 10**. Separation of regular and outlier DEG responses within each iteration for the (A) T, (B) 3 CT groups, (C) NvsCT, (D) PvsNP, (E) T & 3 CT groups and (F) T & PvsNP annotations.

**Supplementary Table 11**. Initial and final clustering set-selective efficiencies and cosine similarities of set-selective DEGs for the (A) T, (B) 3 CT groups, (C) NvsCT, (D) PvsNP, (E) T & 3 CT groups and (F) T & PvsNP annotations.

## SUPPLEMENTARY APPENDIX

Pairwise correlation of the unprocessed DEG vectors, followed by hierarchical clustering grouped DEG vectors induced in the same cell line predominantly into the same clusters (Suppl. Fig. 1). DEG vectors induced by the same drug treatment, e.g., sequential doxorubicin and low dose trastuzumab (Suppl. Fig. 13A) or induced in cardiomyocyte lines obtained from patients belonging to the same cardiotoxicity group (Fig. 2B) were distributed across different clusters.

To extract the relevant information from the complete DEG responses, we applied a modified version of our published SVD-based pipeline for the identification of set-selective DEG vectors (Suppl. Fig. 1B). In our previous study,^7^ a set contained all DEG responses induced by the same drug. For each set of unprocessed responses, our pipeline searches for vector components with high similarity to each other. Briefly, our SVD-algorithm searched for set-selective subspaces (spanned by set-selective eigenarrays, also known as principal components), in which all DEG vectors of the same set cluster closely together (after projection of the complete data into that subspace). This projection contains the DEGs selective for that set, while set-selective DEGs for other sets were obtained from other subspaces selective for those sets. Using this logic, we defined six different set annotations (Suppl. Fig. 1C, Suppl. Table 2): 1) all DEG responses induced by the same drug treatment regimen (annotation T), 2) all DEG responses induced in cell lines of the same CTRCD group (3 CT groups), 3) all DEG responses induced in no-CTRCD and CTRCD lines (NvsCT), 4) all DEG responses induced in persistent or non-persistent CTRCD lines (PvsNP), 5) all DEG responses induced by the same drug treatment in cell lines of the same CTRCD group (T & 3 CT groups) and 6) all DEG responses induced by the same drug treatment in either persistent or non-persistent CTRCD lines (T & PvsNP annotation). We selected these six set annotations, since we were testing whether our algorithm could identify gene expression profiles indicative of any CTRCD groups. For each of the six set-annotations our pipeline identified set-selective subspaces. Investigation of the clustering results within those set-selective subspaces identified multiple outlier DEG vectors that did not cluster with the other vectors of the same set (Suppl. Fig. 13B). Consequently, we updated our algorithm so that it can now identify multiple outlier responses rather than only one (see Methods for details). Outlier responses label responses that differ from the set-characteristic responses and could be defined as an additional independent set. For the next iteration, we instructed the algorithm to separate regular and outlier responses into two new sets, if each new set contained at least four DEG responses that additionally represent at least 20% of all DEG responses in the parent set. We selected the second cutoff, since within the 3 CT groups, NvsCT and PvsNP annotations all seven treatments of each cell line are assigned to the same set (i.e., persistent, non-persistent, reversible and no-CTRCD).

DEG vectors labeled as regular in one iteration can become outliers in the next iteration. Consequently, the algorithm could continue to split the sets until each was dominated by a single cell line or a few similar cell lines. CTRCD group-selective responses would be obscured again, as they are in the complete dataset. Iterative application of our pipeline after separation of regular and outlier responses based on the T, 3 CT groups, NvsCT, PvsNP, T & 3 CT groups and T & PvsNP annotation (Suppl. Tables 10A-F, respectively), as described above, improved set-selective clustering, e.g., of sequential doxorubicin and high-dose trastuzumab treatment of reversible CTRCD lines (Suppl. Fig. 13C). Set-associated clustering improved with each iteration for the T, 3 CT groups, NvsCT, PvsNP, T & 3 CT groups and T & PvsNP annotations (Suppl. Fig. 14A to F, Suppl. Tables 11A to F, respectively). As expected, the amount of information from the unprocessed DEG responses preserved within each subspace varied between set annotations for the sets of the T, 3 CT groups, NvsCT, PvsNP, T & 3 groups and T & PvsNP annotations (Suppl. Figs. 14A-F, Suppl. Tables 11A-F, respectively). Our updated pipeline increased the number of sets for the T, 3 CT groups, NvsCT, PvsNP, T & 3 CT groups and T & PvsNP annotations from 7, 3, 2, 2, 21 and 14 initial sets to 21, 8, 5, 4, 26 and 20 final sets (Suppl. Fig. 1C). These final sets were reached within 4, 3, 3, 2, 2 and 3 iterations, respectively.

## Central Illustration. Cardiomyocytes carry a cell-intrinsic signature of persistent cardiotoxicity

Cardiomyocytes derived from induced pluripotent stem cells of 23 women with HER2-positive breast cancer and no (n = 5), reversible (n = 11) or persistent (n = 7) CTRCD were exposed to seven doxorubicin and trastuzumab regimens and profiled by transcriptomics. Responses of persistent CTRCD lines were resolved from those of the other groups and differed in contraction, calcium handling, cytoskeletal, glycolytic and natriuretic peptide receptor signaling pathways, with implications for selection of cardioprotective therapy. ARNI = angiotensin receptor/neprilysin inhibitor; CTRCD = cancer therapy-related cardiac dysfunction.

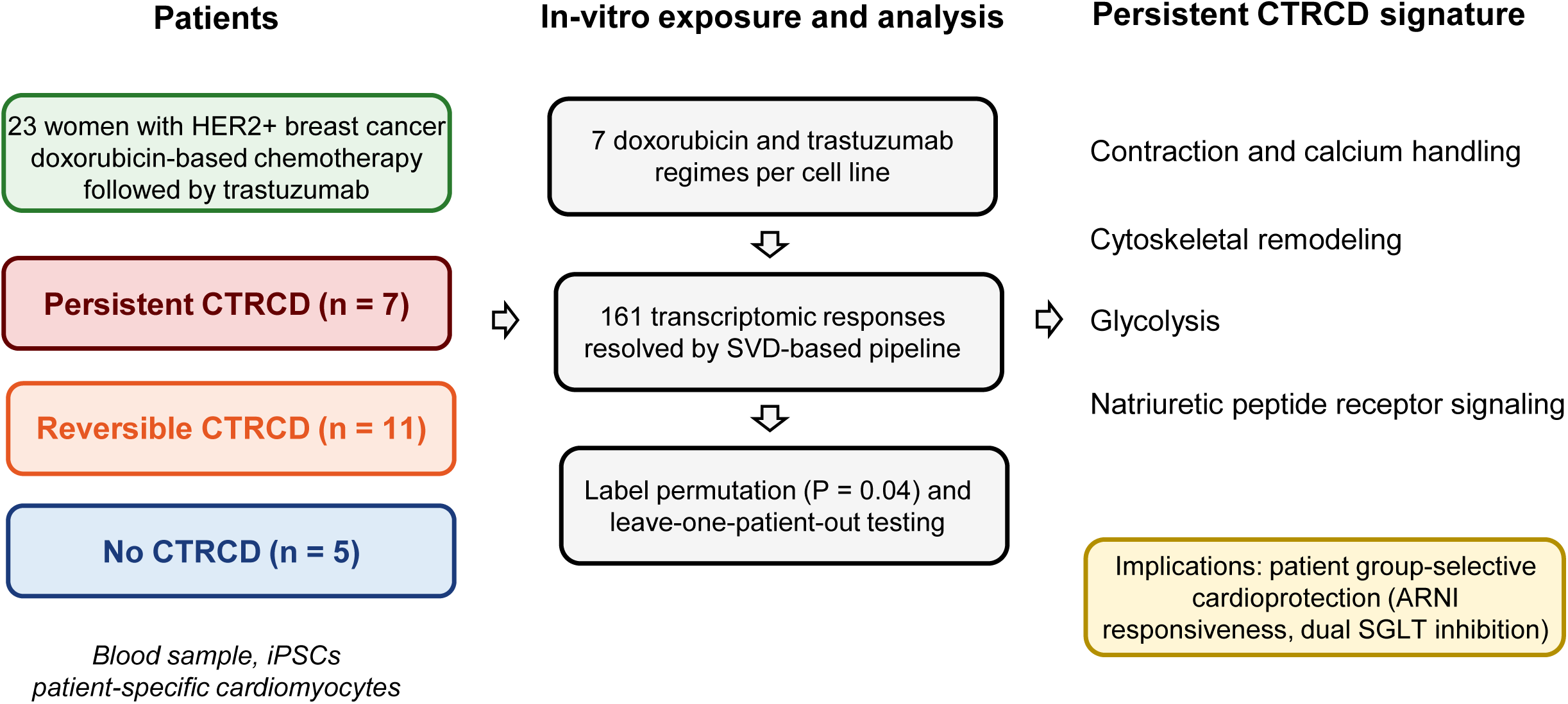

---

**Supplementary Figure 1.**
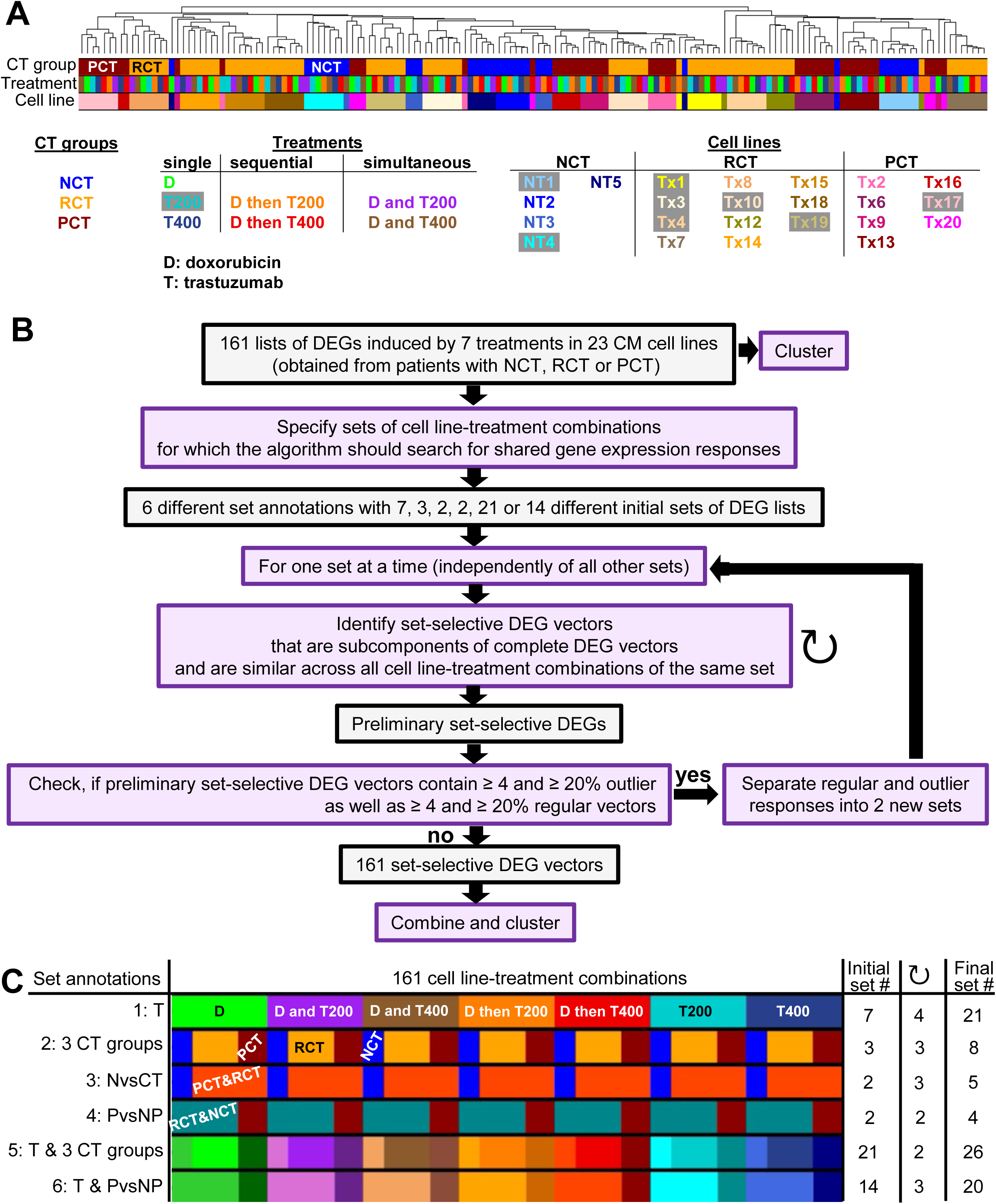
Clustering resolution, SVD-pipeline and set annotations. **(A)** 161 complete DEG lists were subjected to pairwise correlation analysis, followed by hierarchical clustering. Grouping of DEG lists is primarily driven by the treated cell lines, suggesting that the response vectors are cell-line selective and that drug- or CTRCD group-selective effects are obscured. The dendrogram is also shown in Figure 2A and Supplemental Figures 2A to F. **(B)** We updated our SVD-based algorithm for the identification of set-selective gene expression profiles and implemented iterative separation of outlier and regular responses. **(C)** Treatment regimen-induced responses were assigned to sets that contained all responses induced (1) by the same treatment (T), (2) in cell lines of the same CTRCD group (3 CT groups), (3) in cell lines with no-CTRCD or with reversible and persistent CTRCD (NvsCT), (4) in cell lines with persistent or non-persistent CTRCD (PvsNP), (5) by the same treatment in cell lines of the same CTRCD group (T & 3 CT groups) and (6) by the same treatment in cell lines with persistent or non-persistent CTRCD (T & PvsNP). Relationships between the six set annotations can be extracted from color overlaps. ↻: iterations. D: doxorubicin, T200, T400: trastuzumab 200 μg/mL, 400 μg/mL.

**Supplementary Figure 2.**
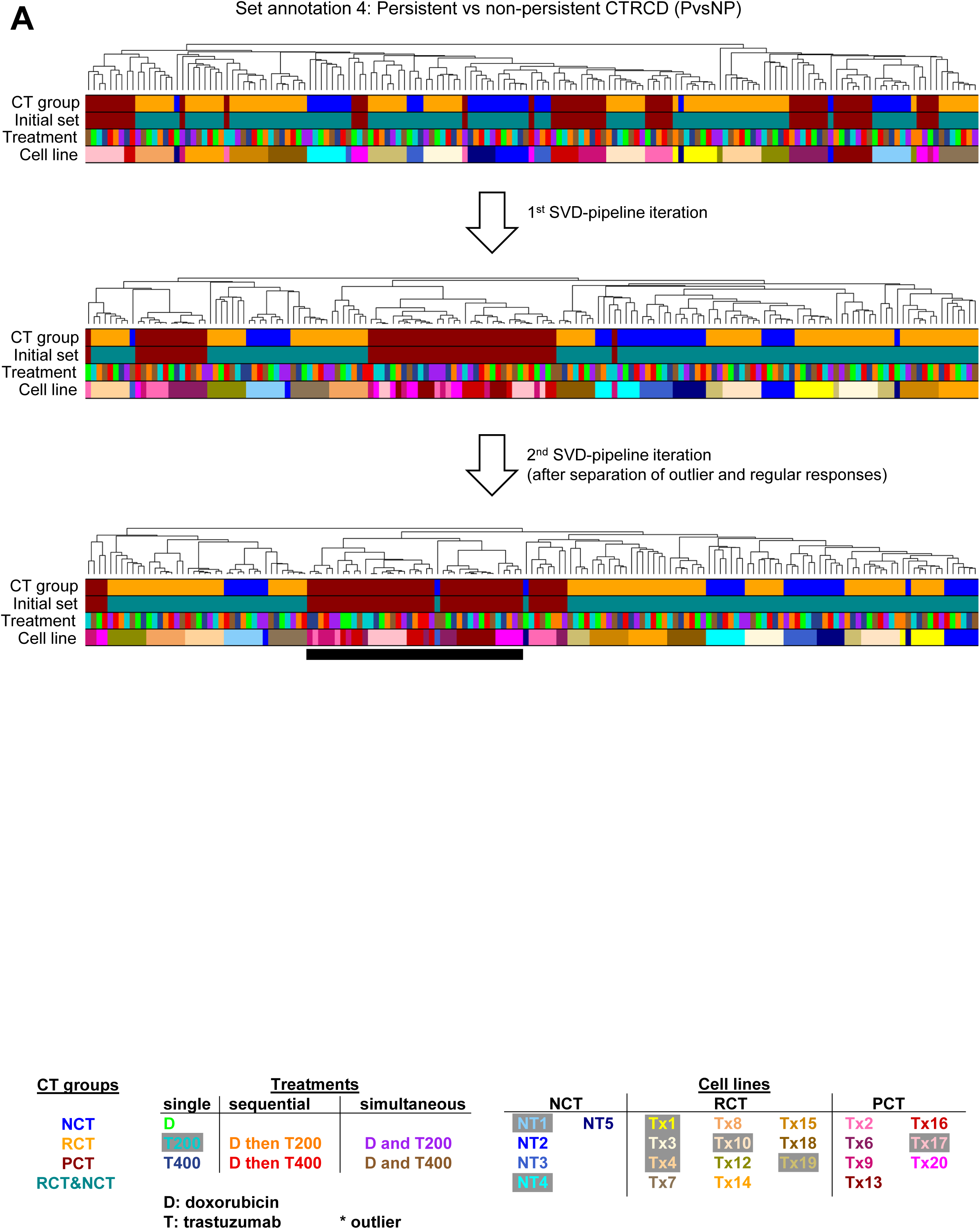
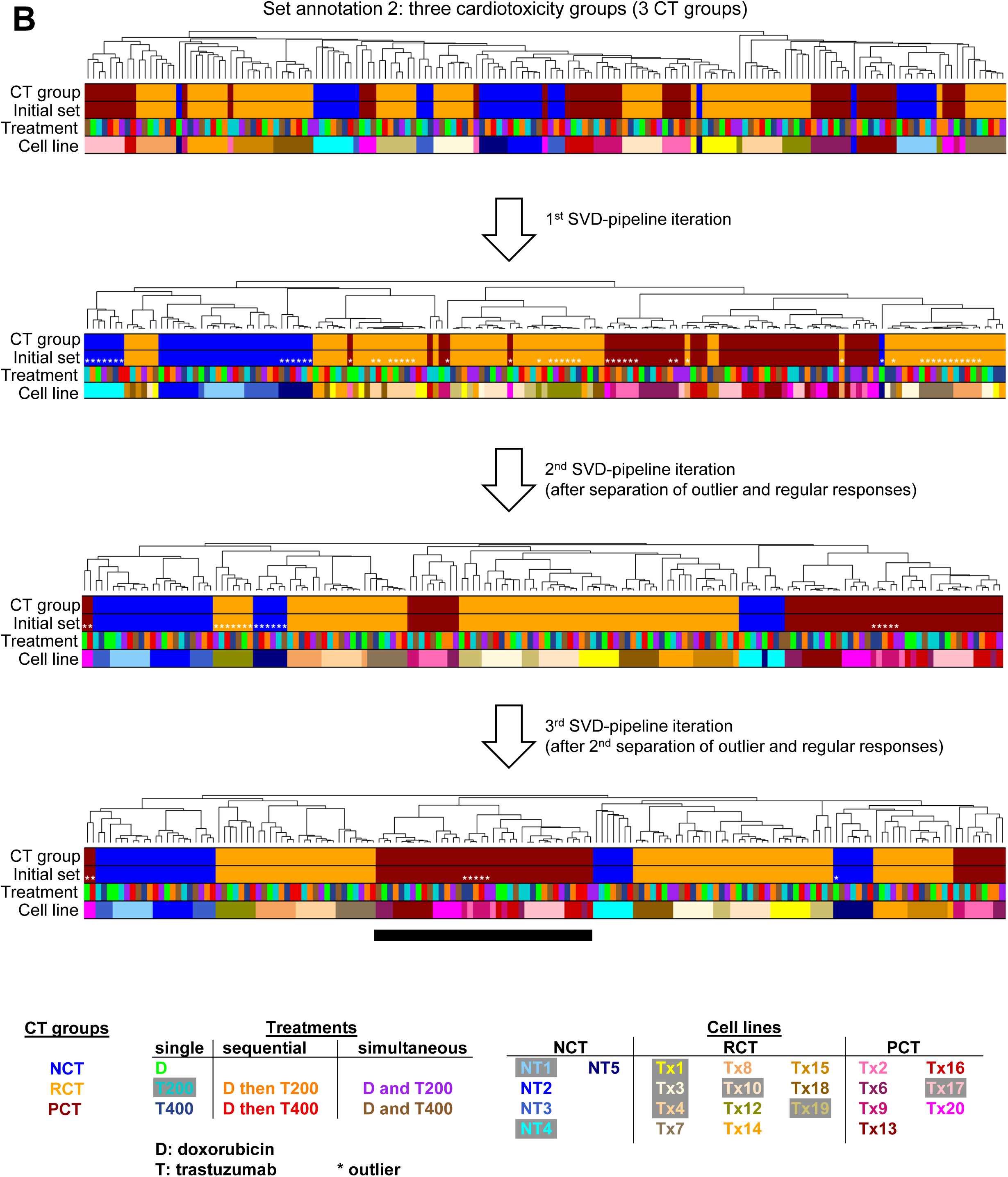
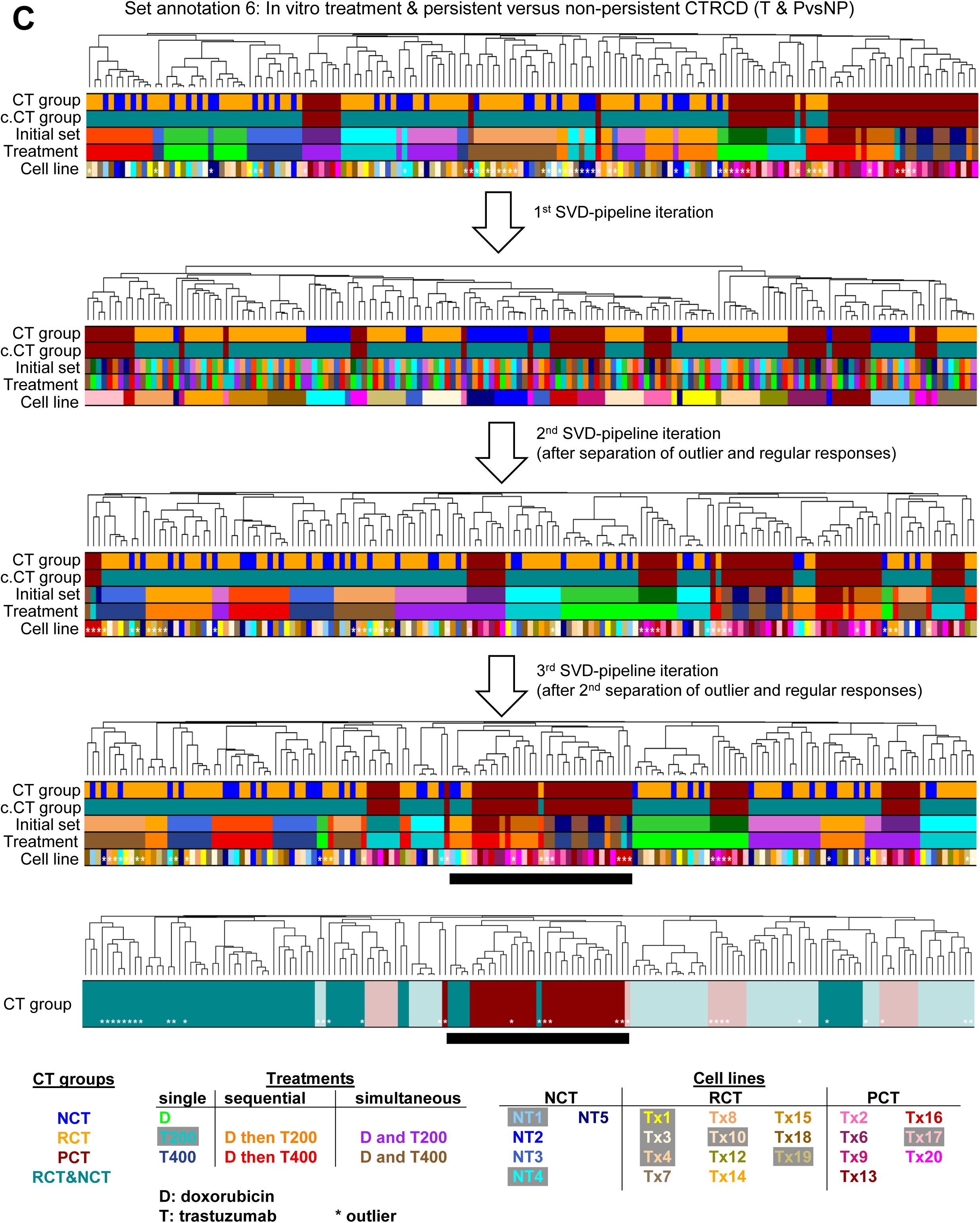
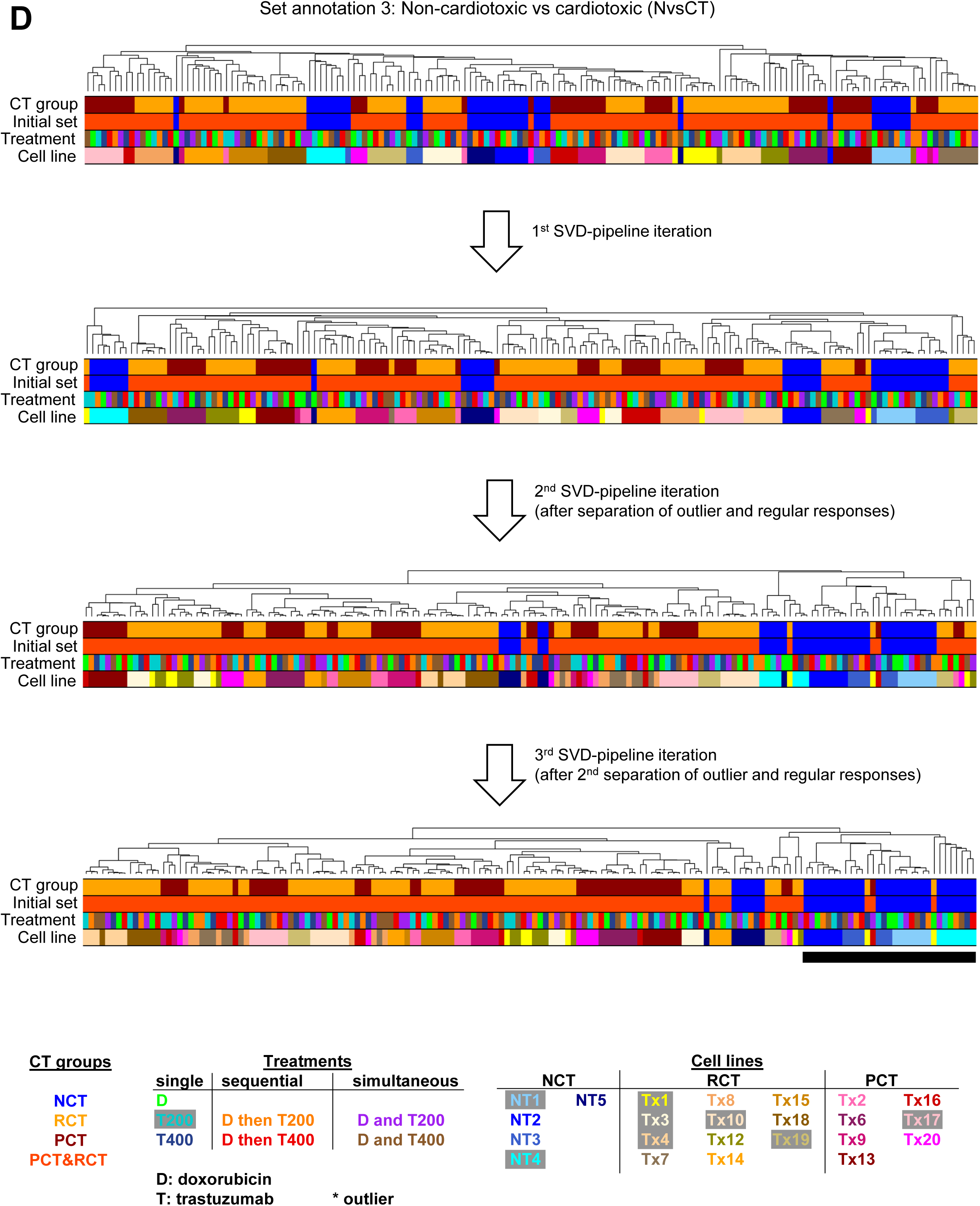
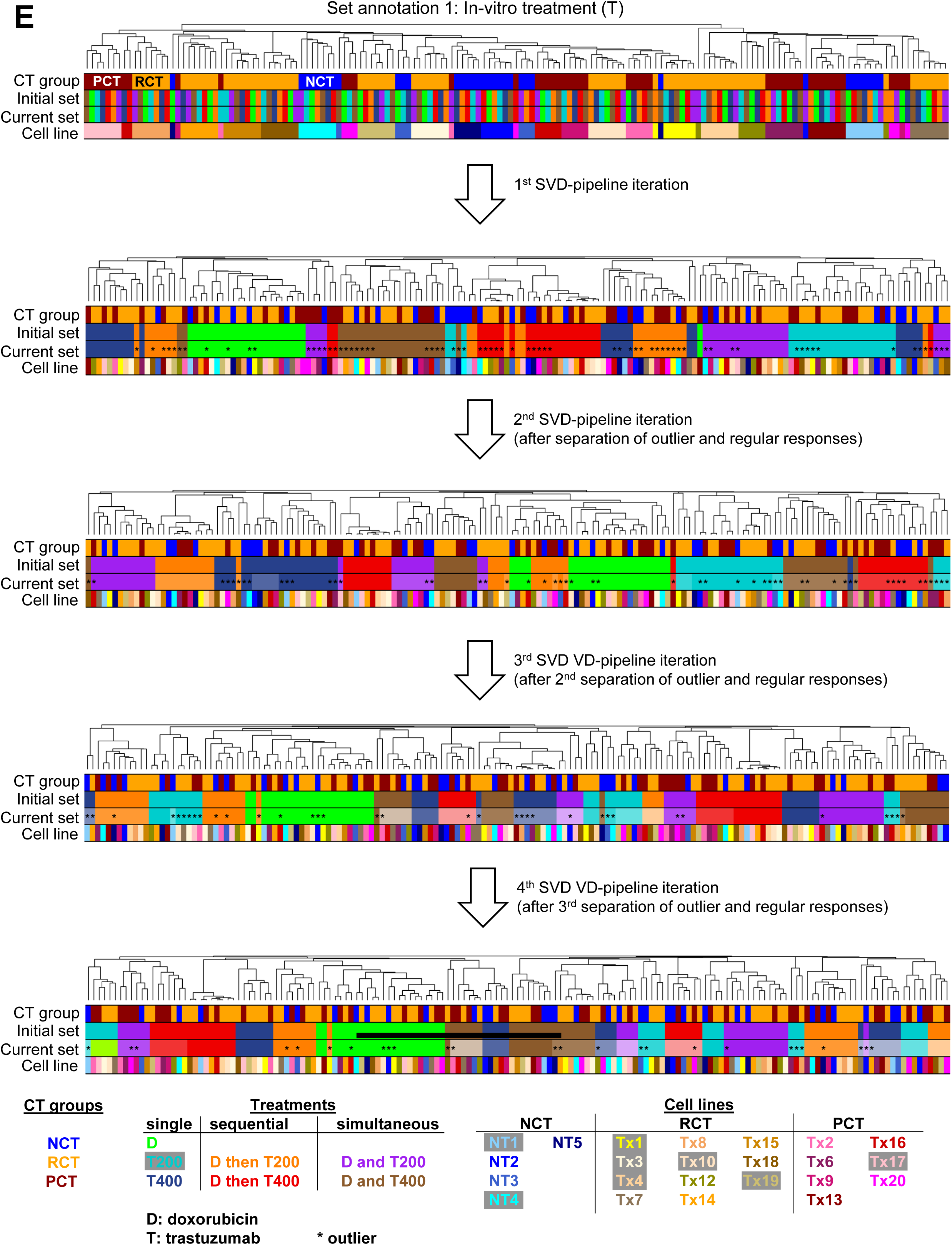
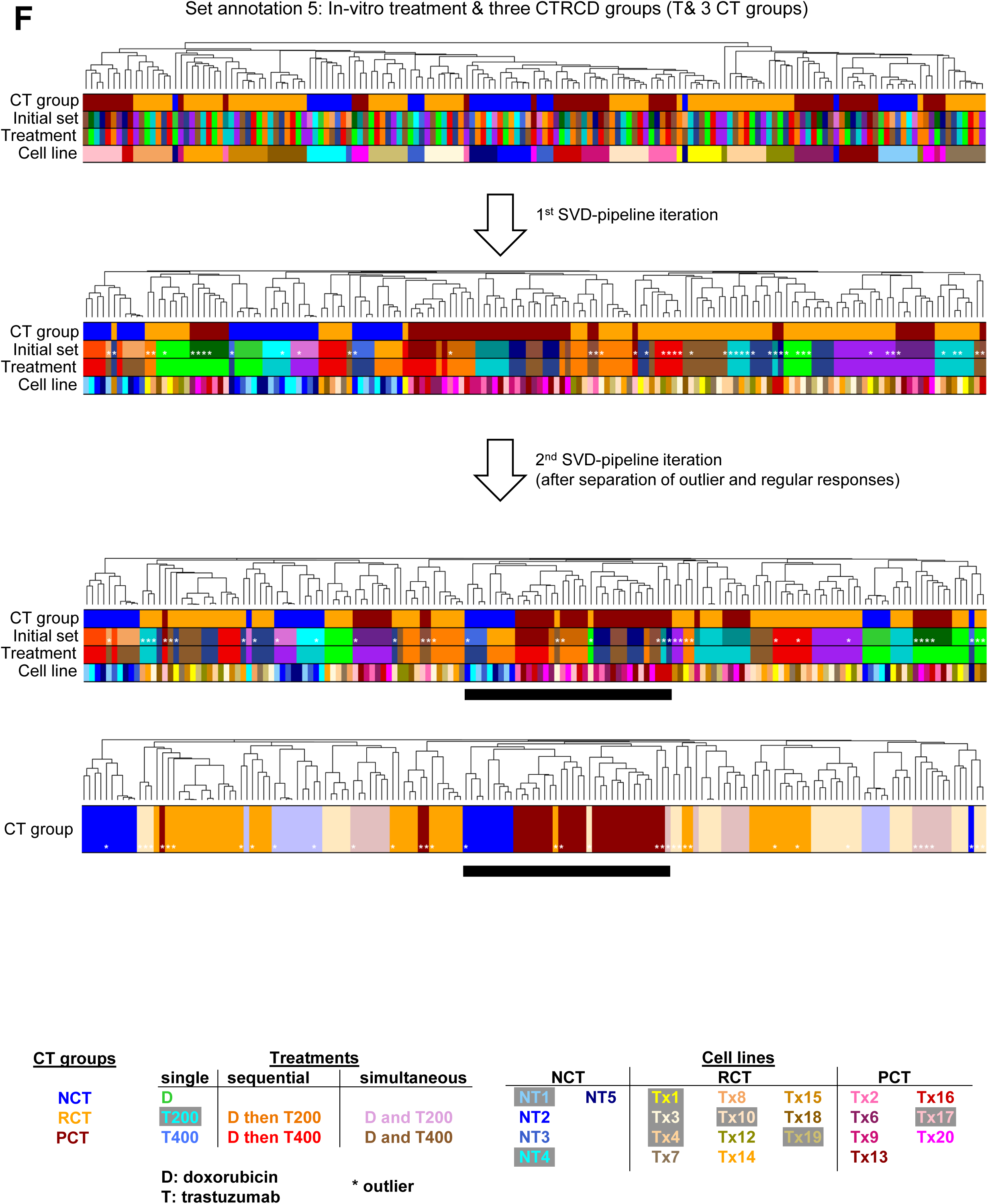
Unprocessed and combined set-selective DEGs across iterations of the SVD-pipeline. Unprocessed and set-selective DEGs obtained after each iteration using the **(A)** PvsNP, **(B)** 3 CT groups, **(C)** T & PvsNP, **(D)** NvsCT, **(E)** T and **(F)** T & 3 groups annotations were combined into a new matrix and subjected to pairwise correlation analysis, followed by hierarchical clustering. Asterisks label outlier responses in the set-selective subspaces (see methods and supplementary appendix for detail). Black horizontal bars label clusters with an enrichment of DEG vectors of the same CTRCD group. The clustering dendrogram obtained for the complete data is the same for each annotation and additionally shown in Figure 2A and Supplemental Figure 1A. The final clustering dendrogram of the PvsNP annotation is also shown in Figure 2B. The last dendrogram shown for the (C) T & PvsNP and (F) T & 3 CT groups annotations reflects the final clustering results, but marks only high-dose single or simultaneous and any sequential treatment regimens with bright colors. The remaining treatment regimens are shown in whitened colors. c.CTG: combined CTG.

**Supplementary Figure 3.**
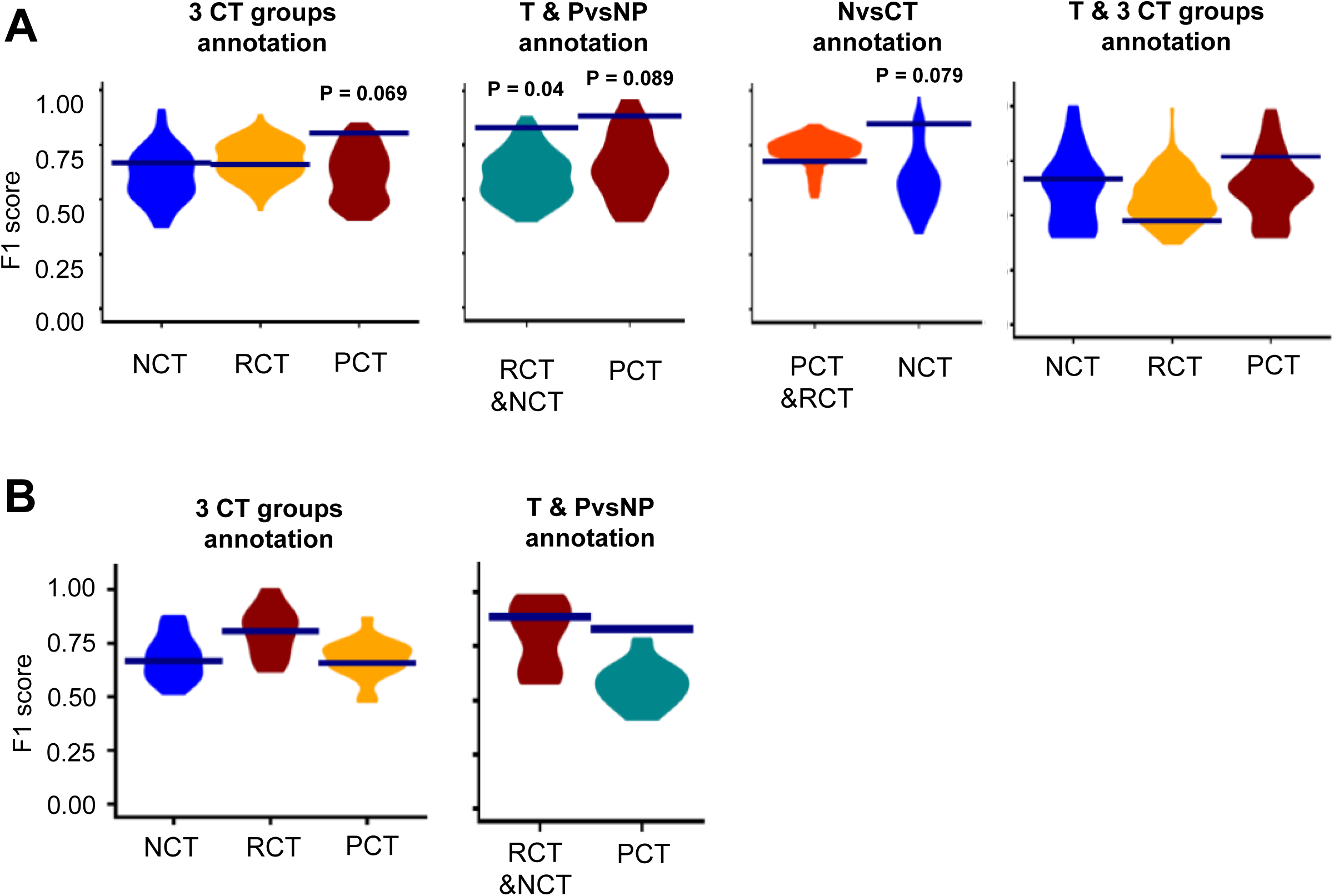
Clustering efficiencies of CTRCD groups in additional annotations. **(A)** Final clustering results of merged set-selective responses obtained for the 3 CT groups, T & PvsNP, NvsCT and T & 3 CT groups annotations were analyzed to quantify the clustering efficiency (F1 score) for each CTRCD group (blue lines). For the T & PvsNP and T & 3 CT groups annotations, only high-dose single or simultaneous as well as any sequential treatments were considered. CTRCD group labels were shuffled 100 times, followed by set reassignment and submission to the iterative SVD pipeline. Clustering efficiencies obtained with true (blue lines) and shuffled labels (violins) were compared to calculate Monte Carlo P values (no label: P > 0.1). **(B)** Clustering efficiencies in the complete data (blue lines) compared with those obtained in the 23 leave-one-patient-out datasets (violin). Abbreviations as in Suppl. Fig. 1.

**Supplementary Figure 4.**
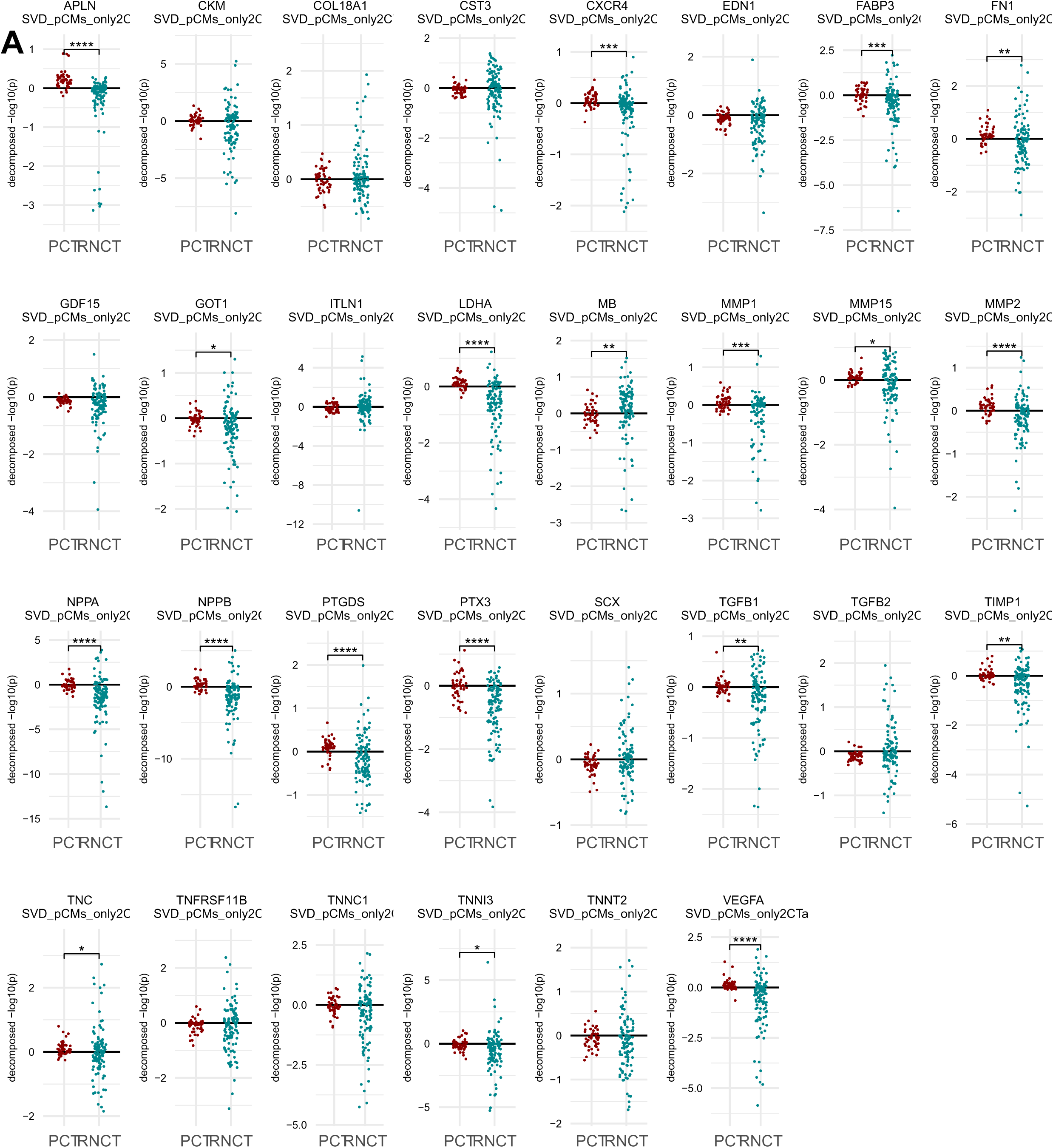
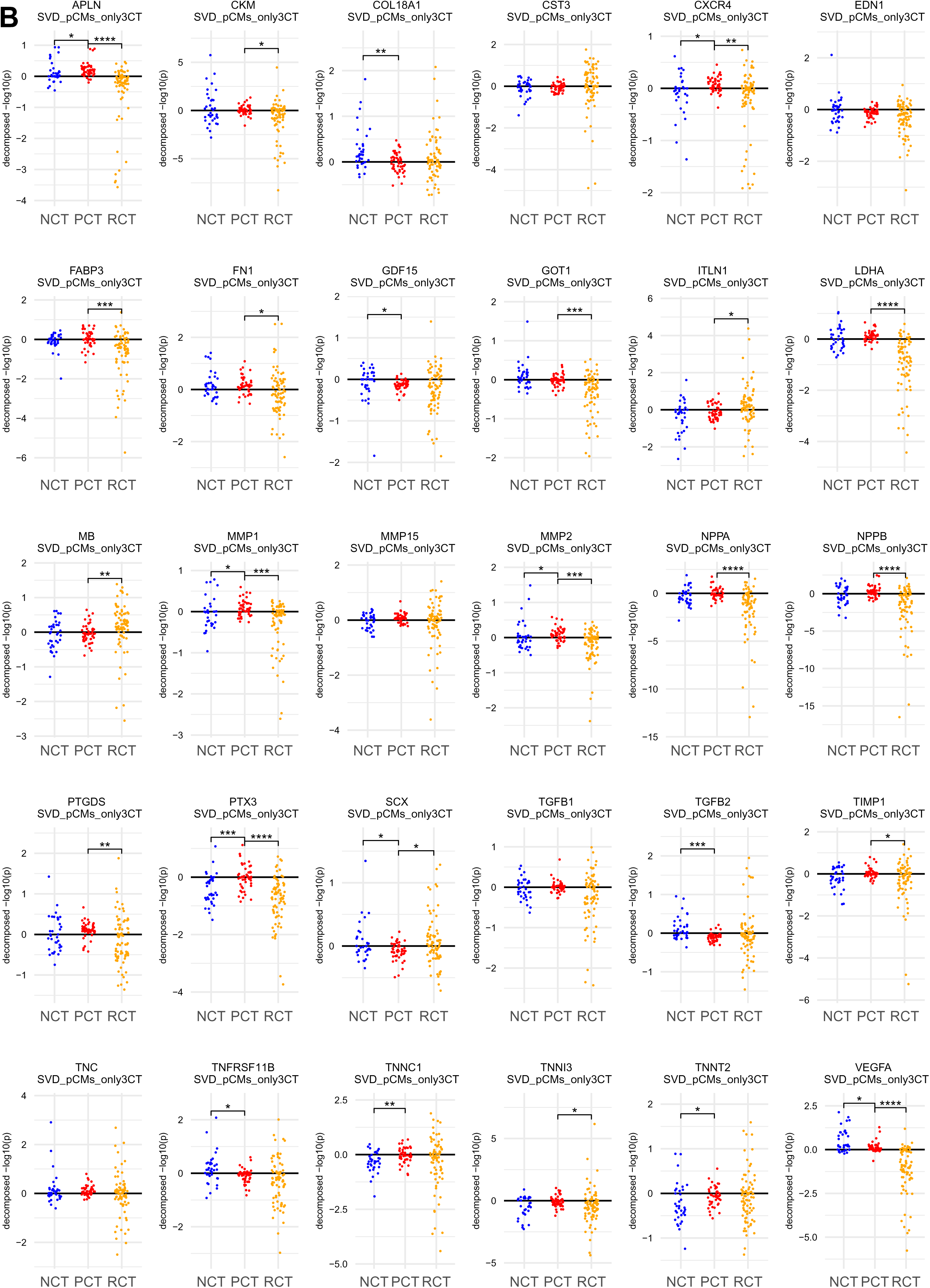
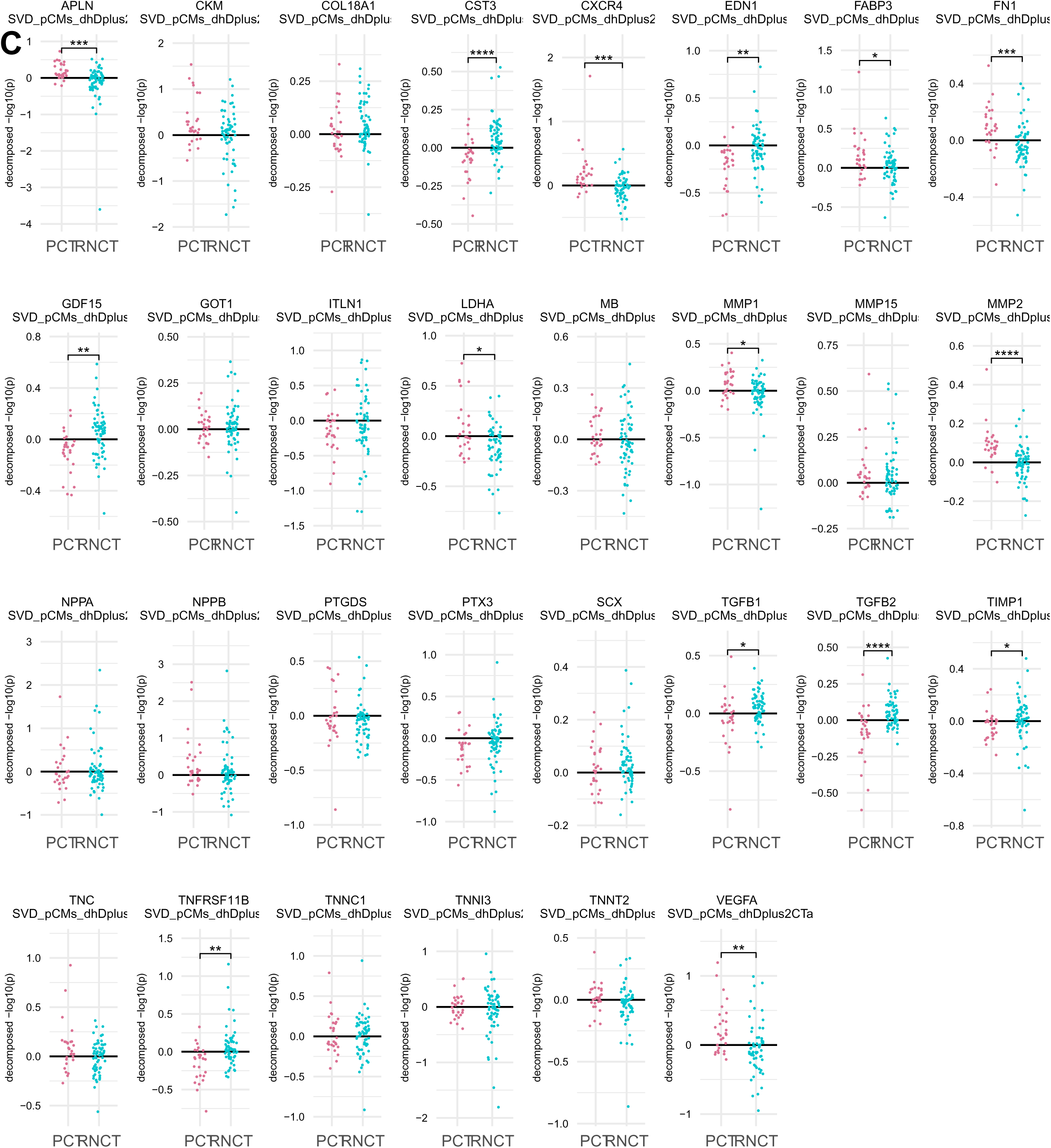
Levels of genes coding for suggested serum biomarkers. Set-selective induction levels for literature-curated genes coding for suggested serum markers of dilated cardiomyopathy, hypertrophic cardiomyopathy or cardiac damage were compared between given CTRCD groups based on **(A)** the PvsNP, **(B)** 3 CT groups and **(C)** T & PvsNP annotations, using the Wilcoxon rank sum test. *, **, ***, ****: adjusted Wilcoxon P value ≤ 0.05, ≤0.01, ≤0.001, ≤0.0001. n.: no significance. Abbreviations as in Figures 1 to 3. Figure 3A shows the same data for selected genes and the PvsNP annotation.

**Supplementary Figure 5.**
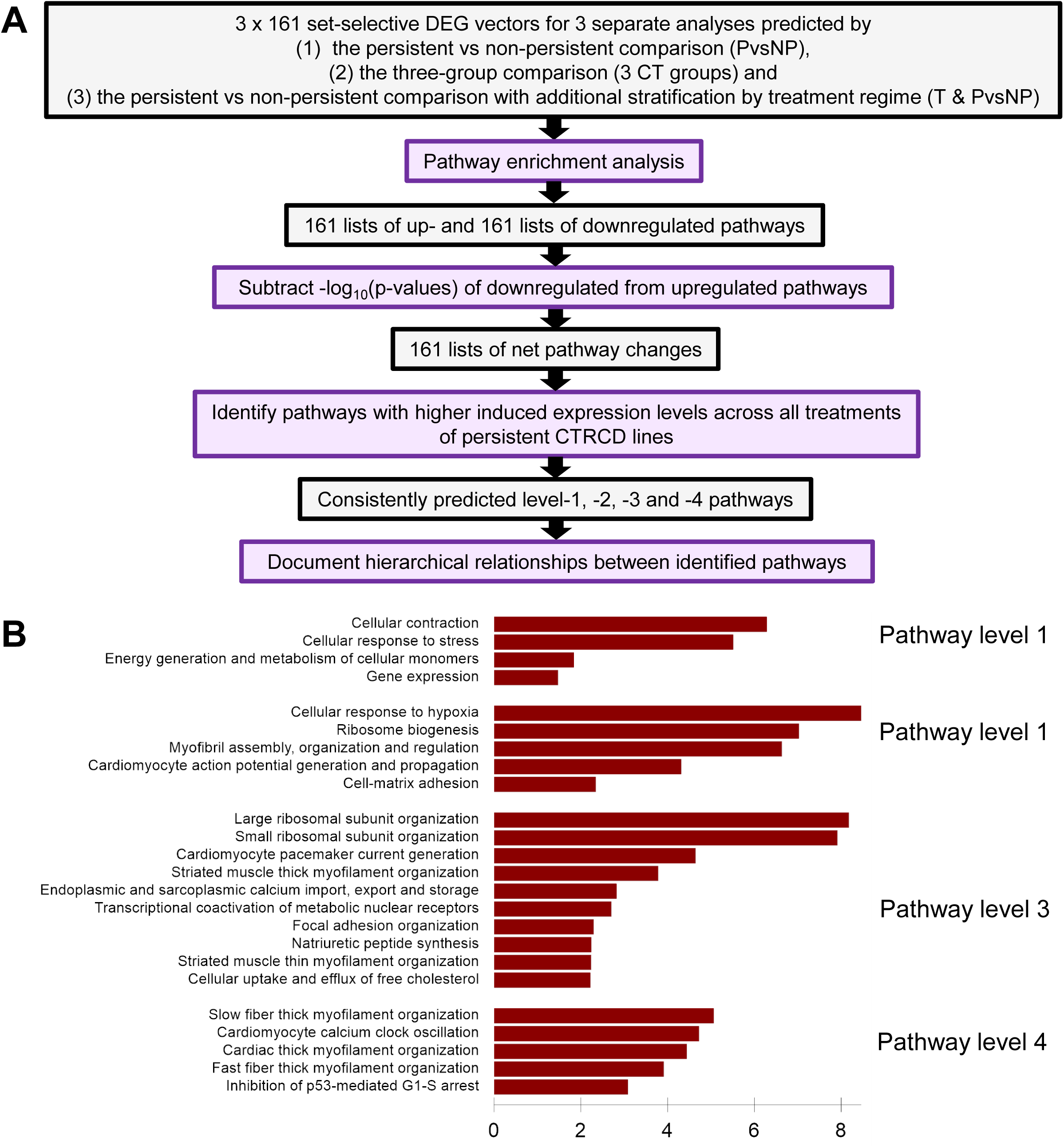
Pathway analysis workflow. **(A)** Up- and downregulated set-selective DEGs identified for the PvsNP, 3 CT groups and T & PvsNP annotations were subjected to pathway enrichment analysis using Fisher’s exact test and MBCO v2. Predictions were used to identify pathways with a significantly different induced induction between the persistent CTRCD and the non-persistent or reversible CTRCD groups. **(B)** Muscle contractility and electric transmission pathways were among the top significant 5 level 1, 5 level 2, 10 level 3 and 5 level 4 pathways predicted for the persistent CTRCD cell line Tx9 after sequential treatment with doxorubicin and low-dose trastuzumab.

**Supplementary Figure 6.**
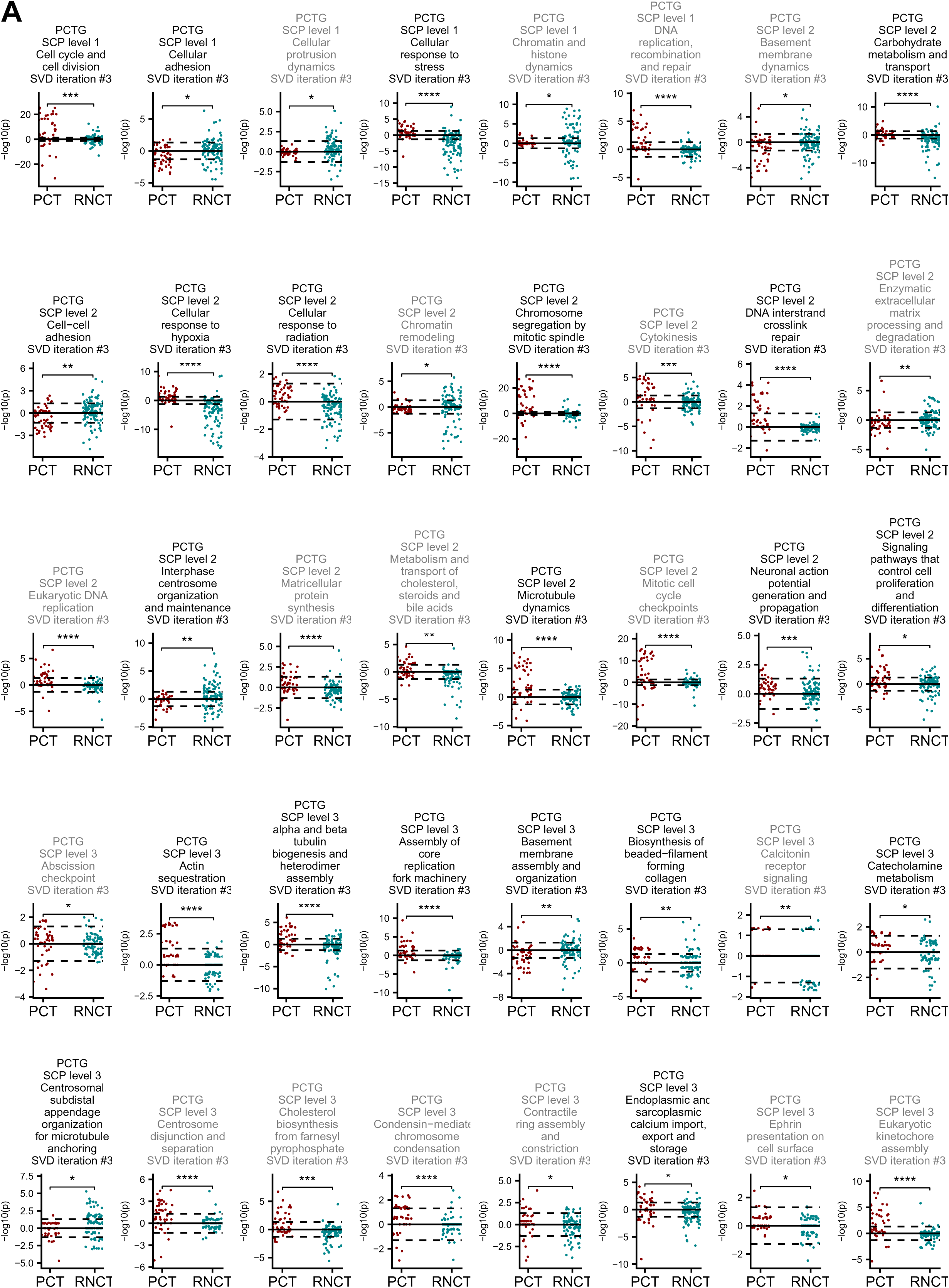
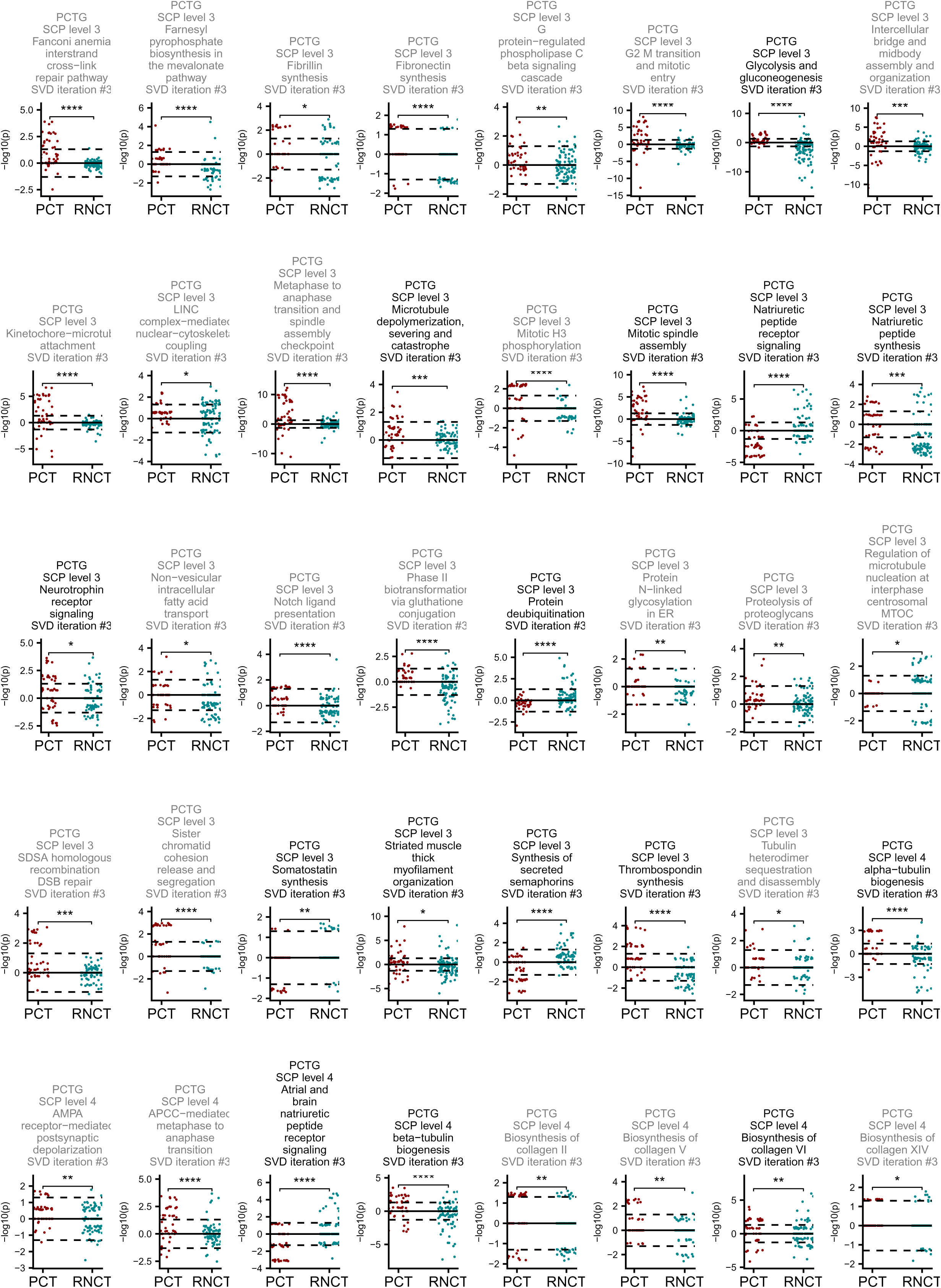
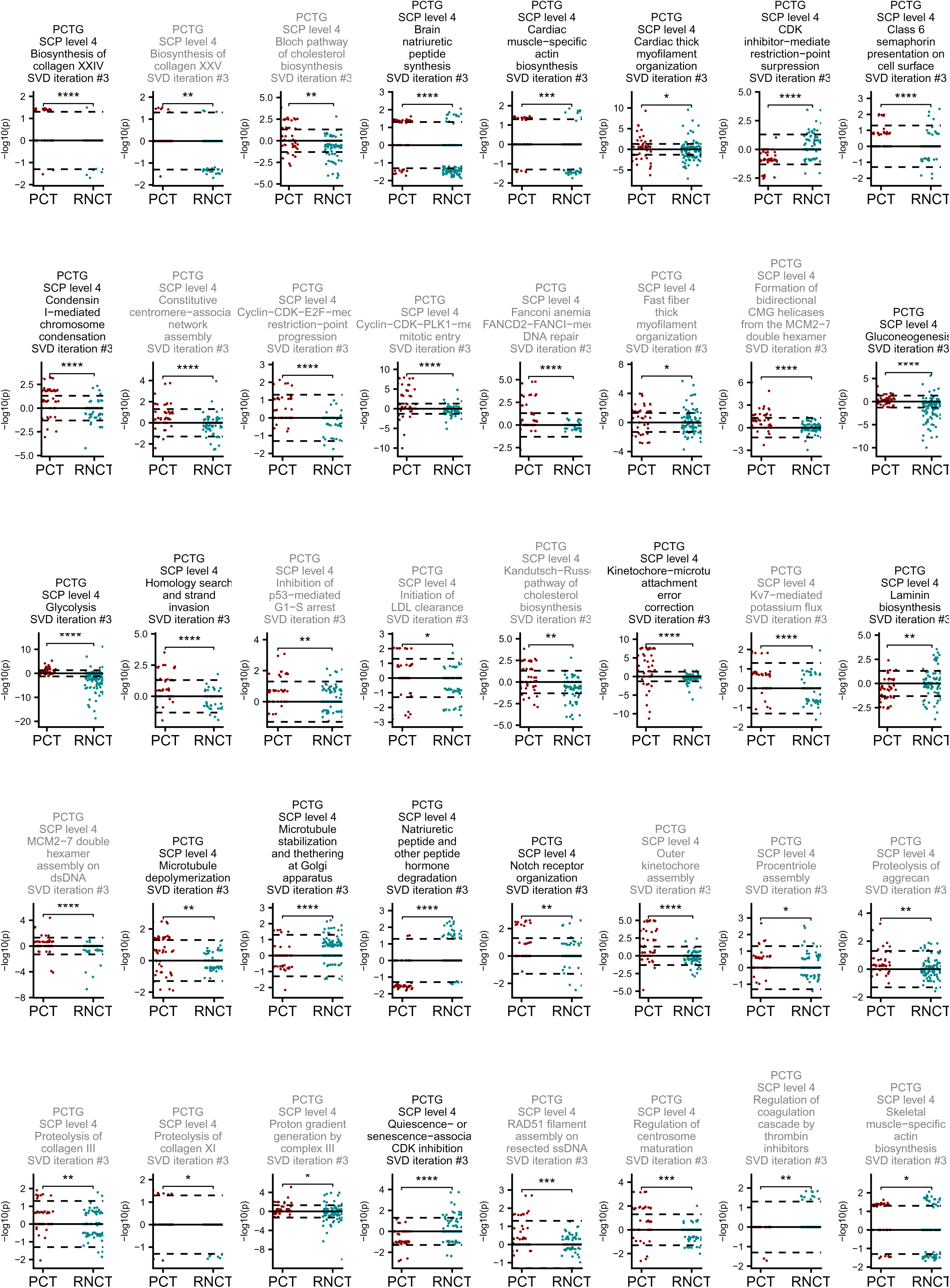
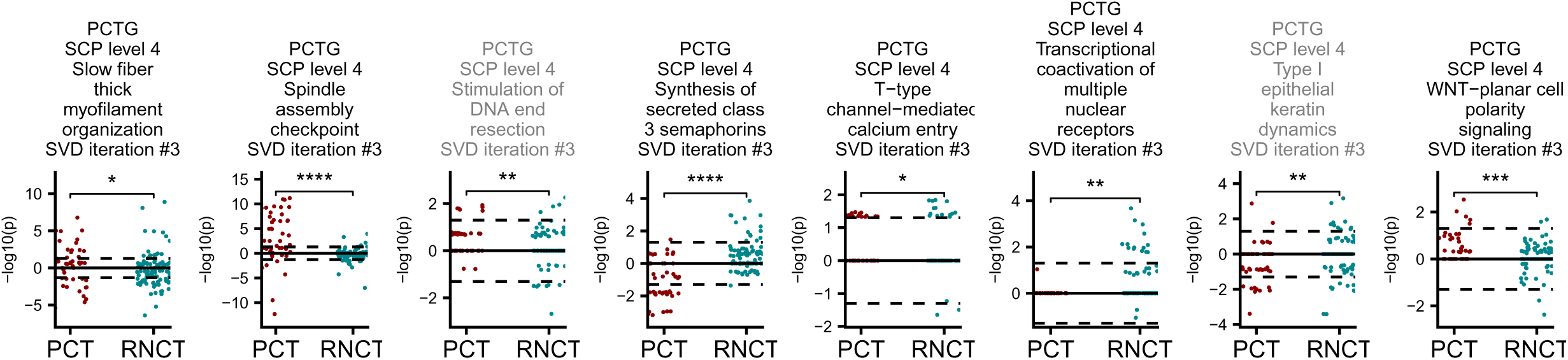
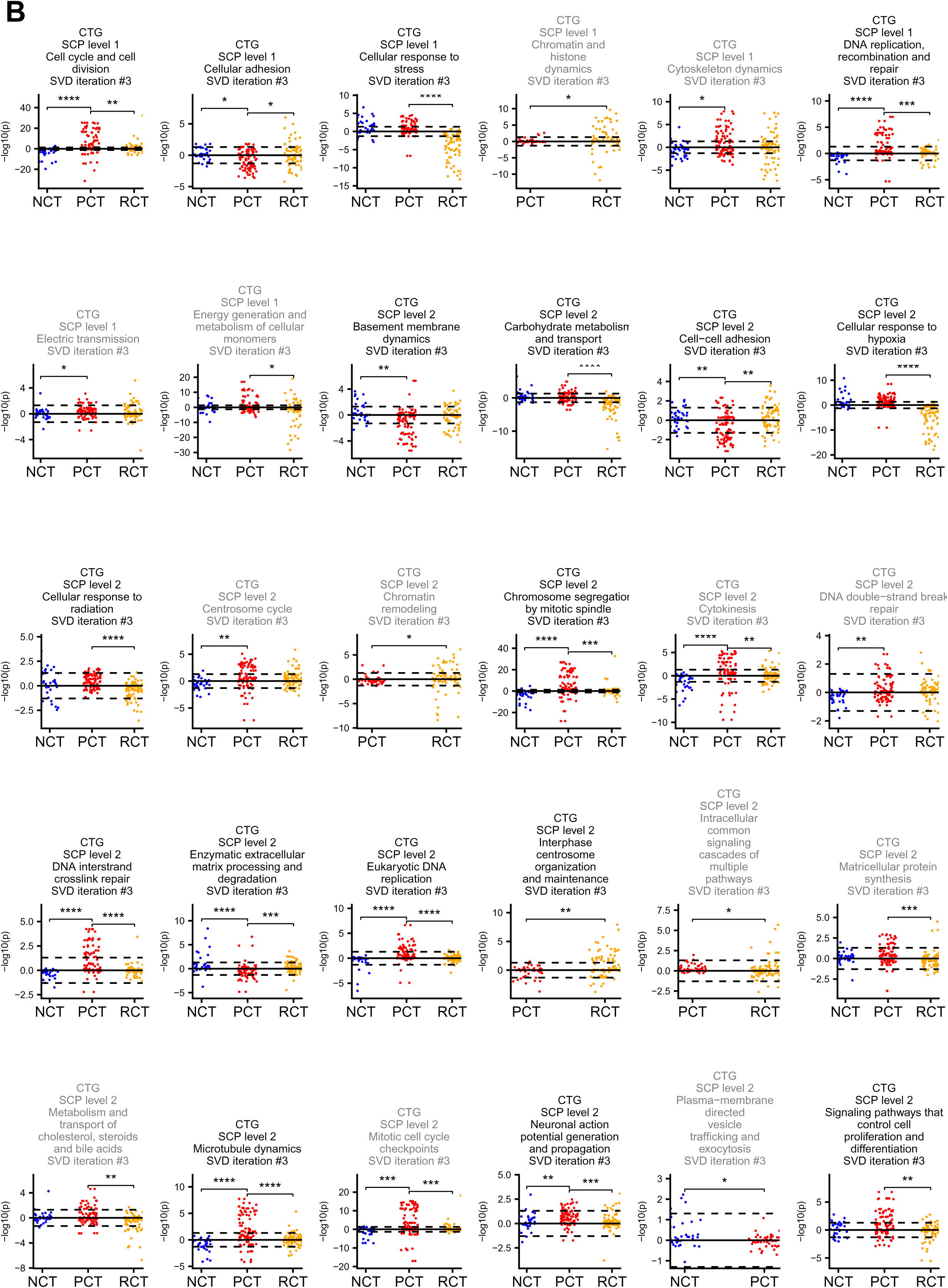
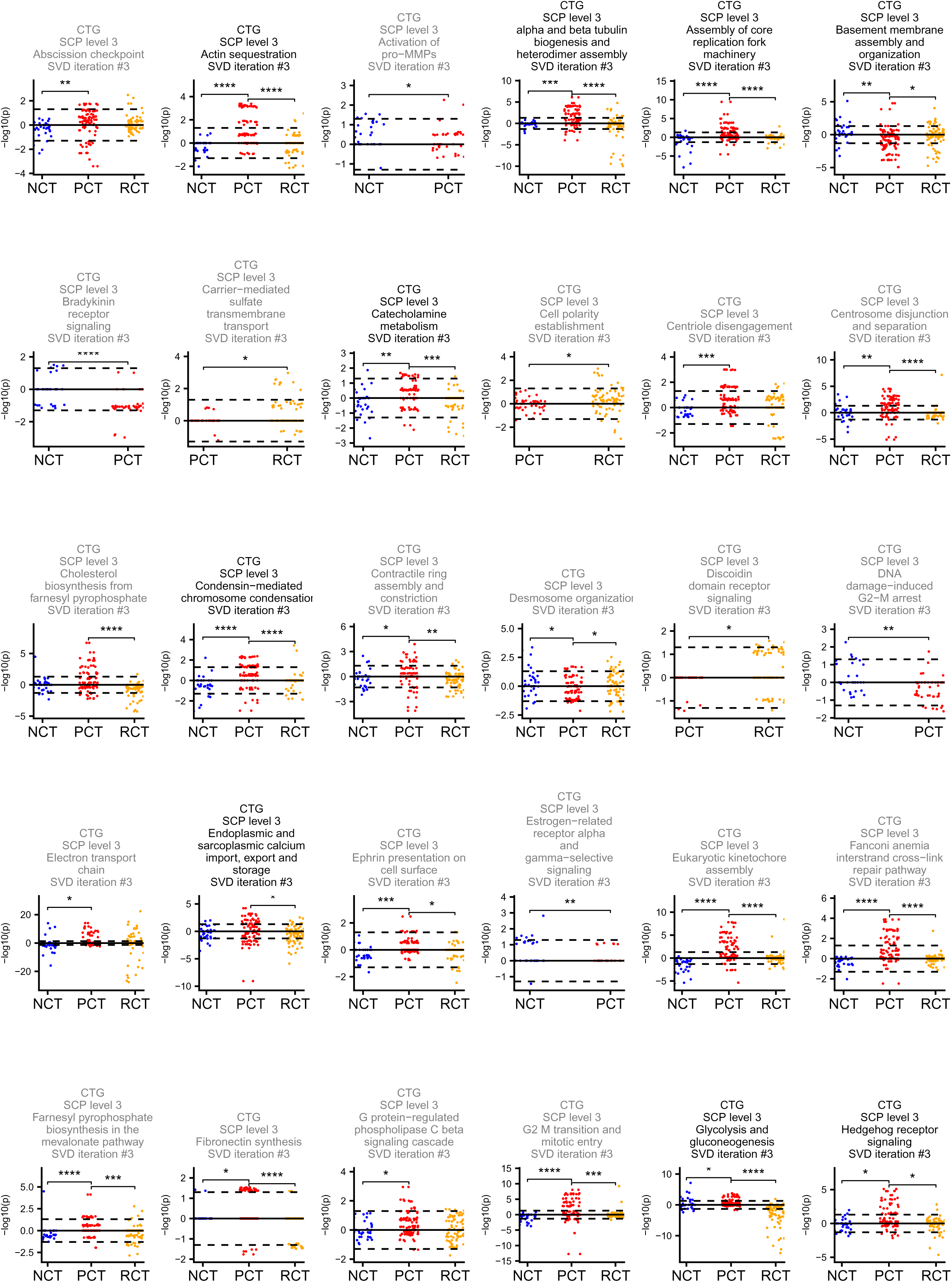
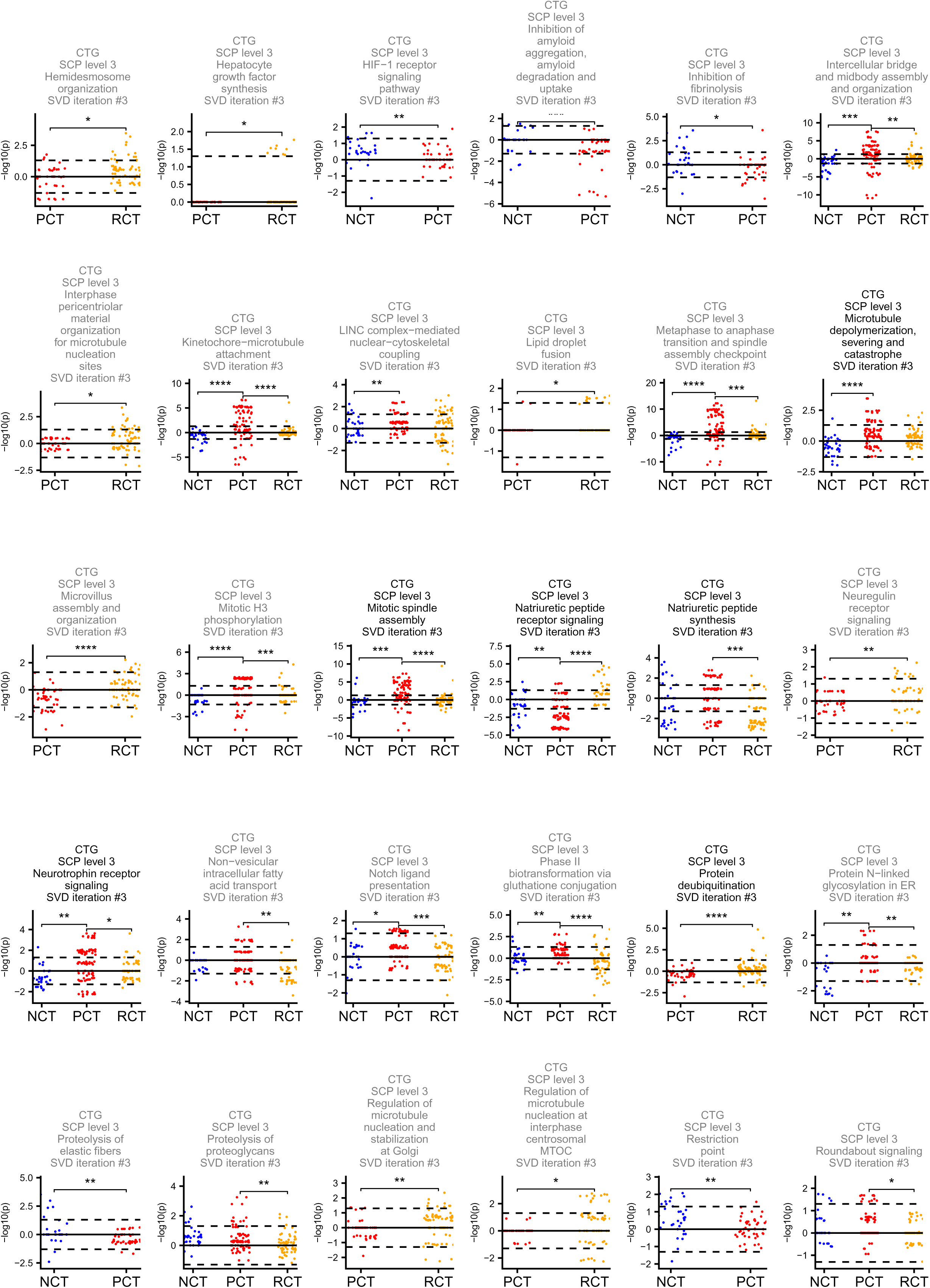
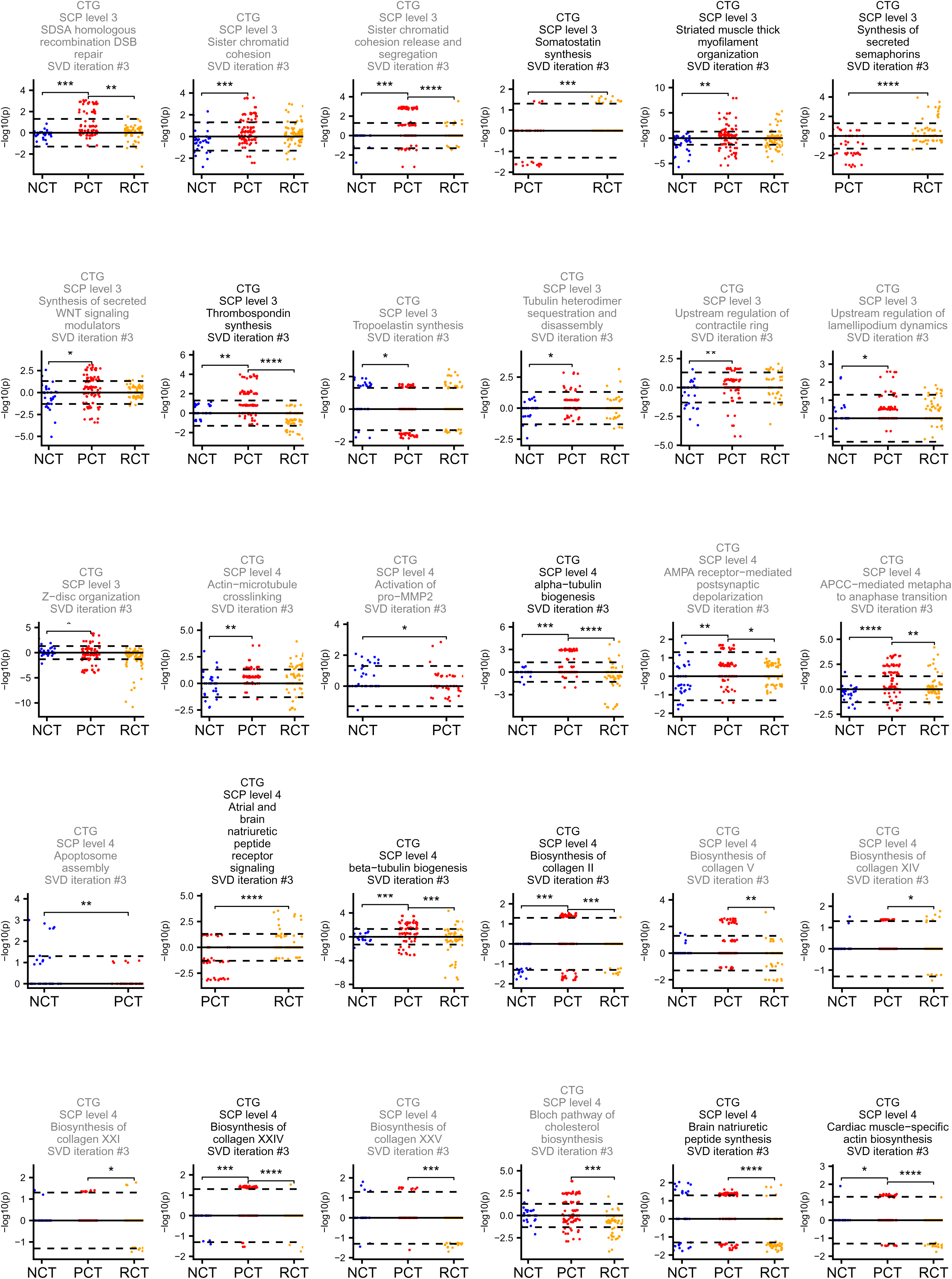
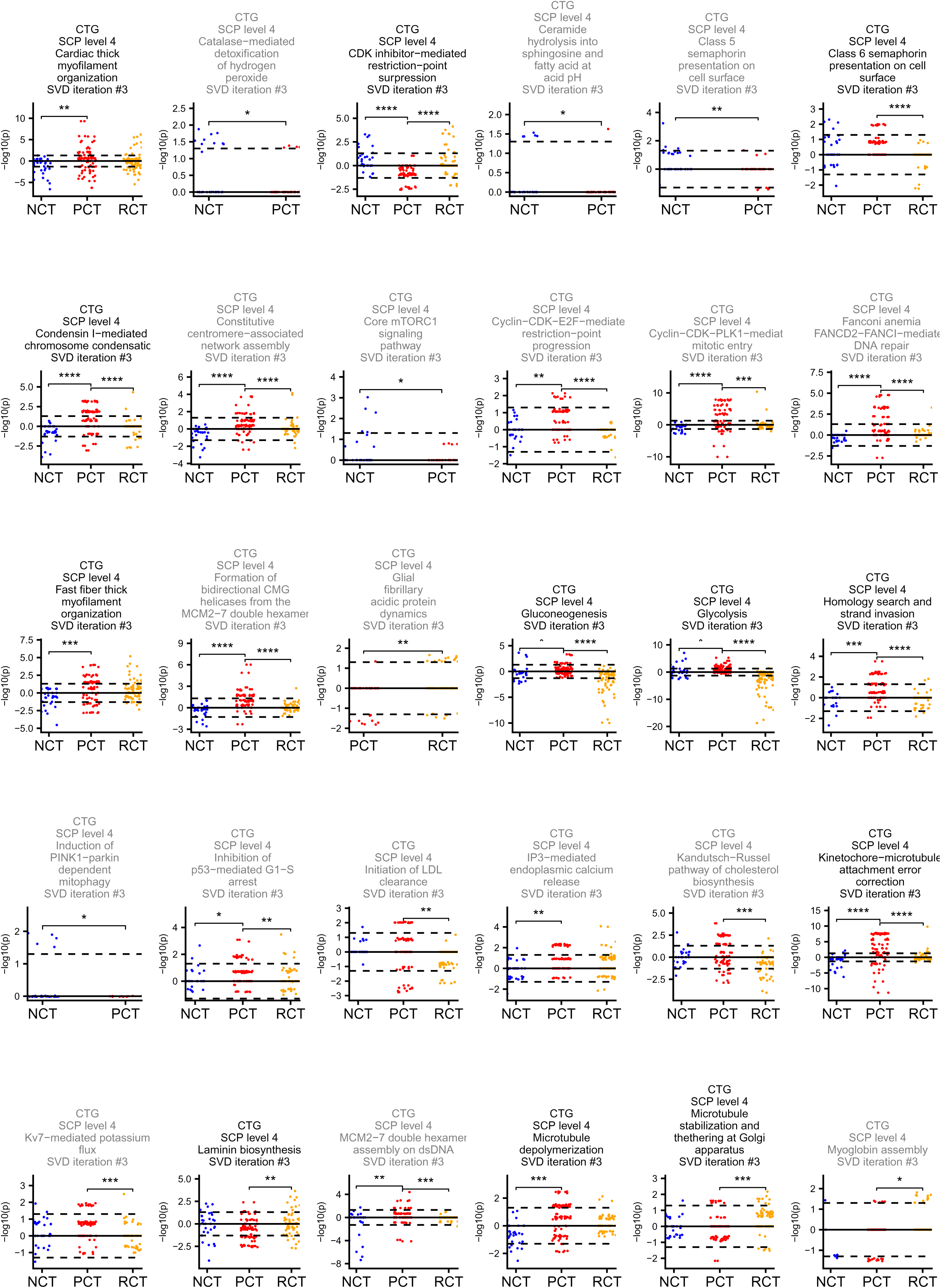
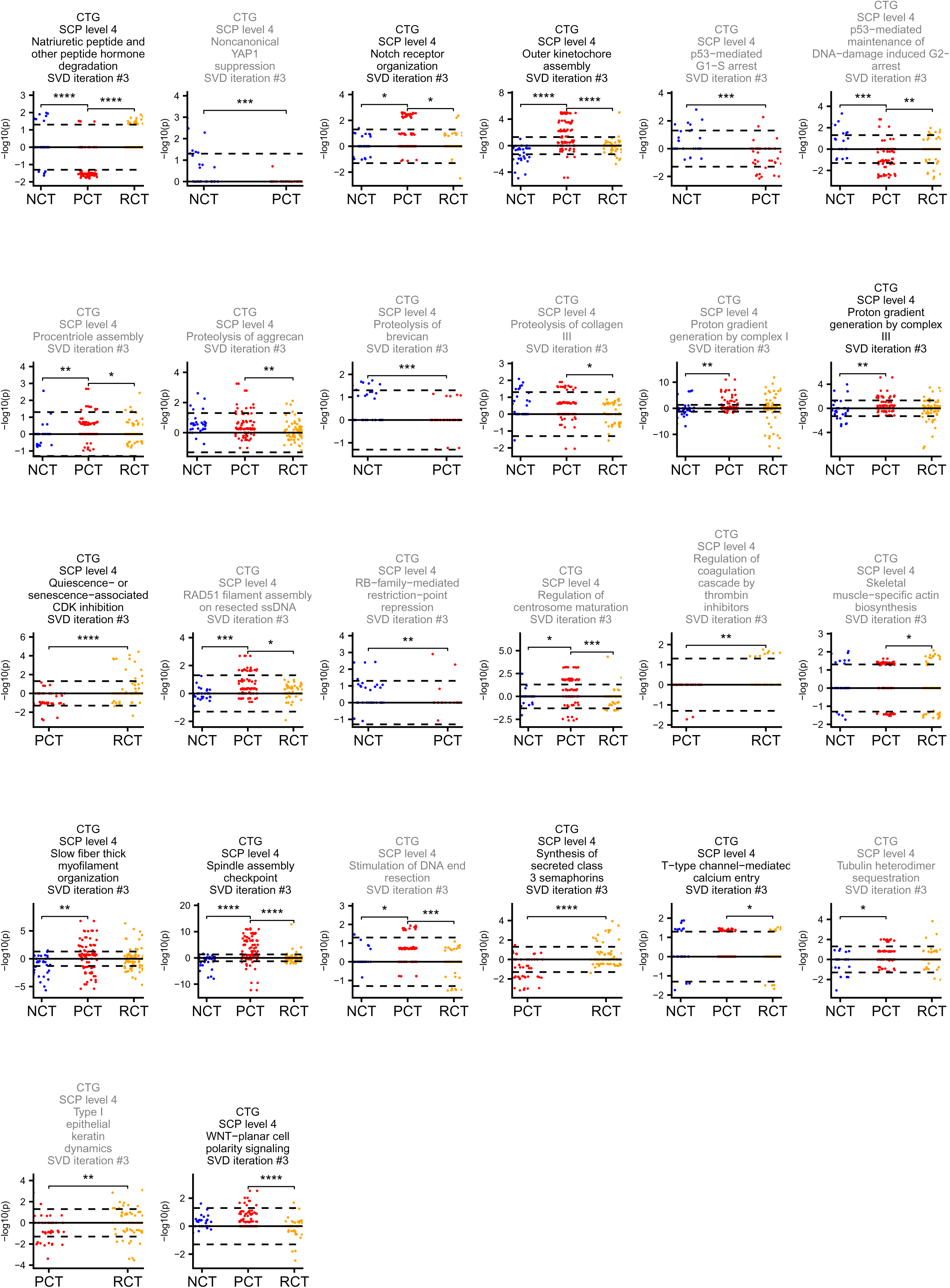
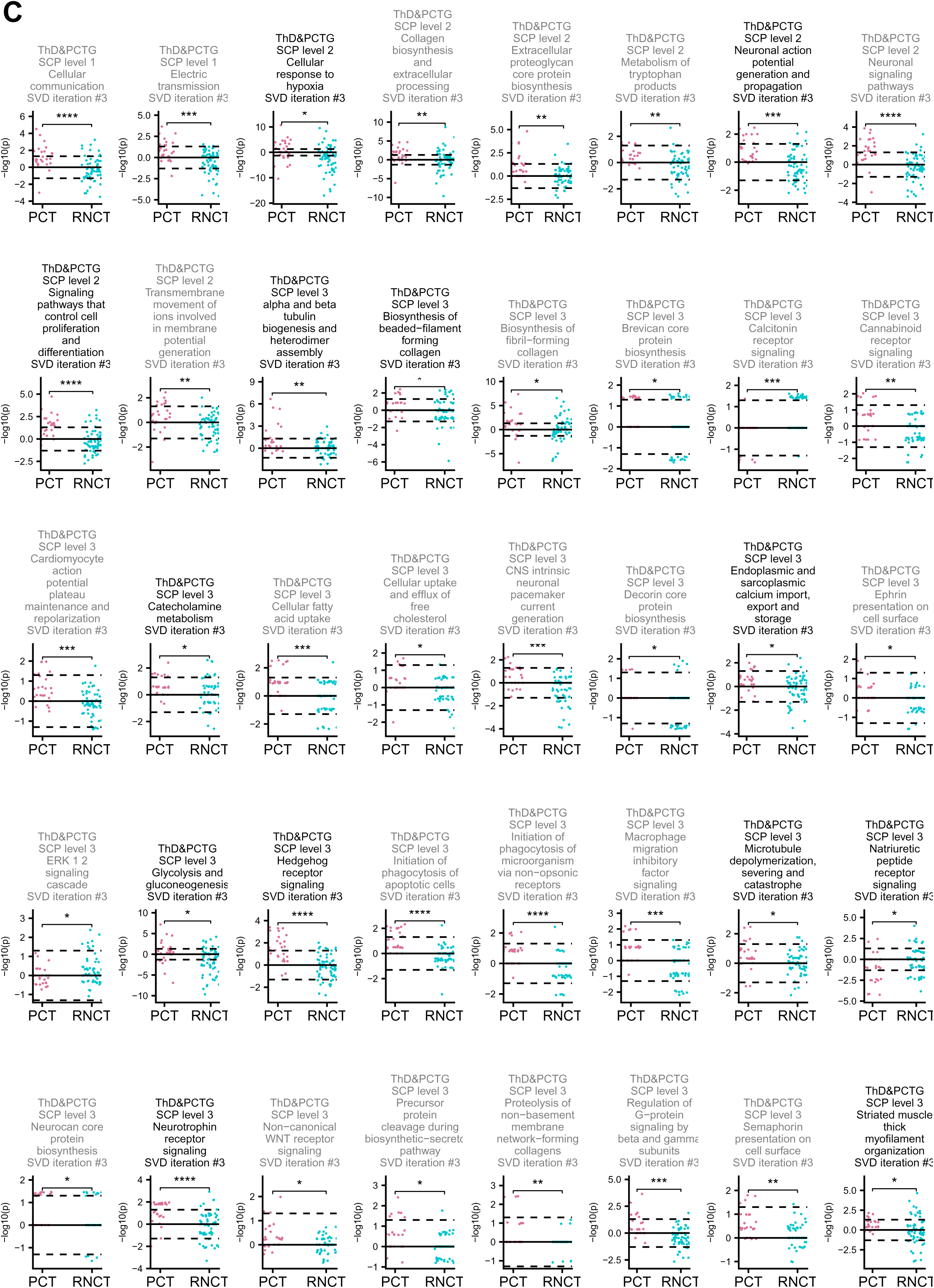
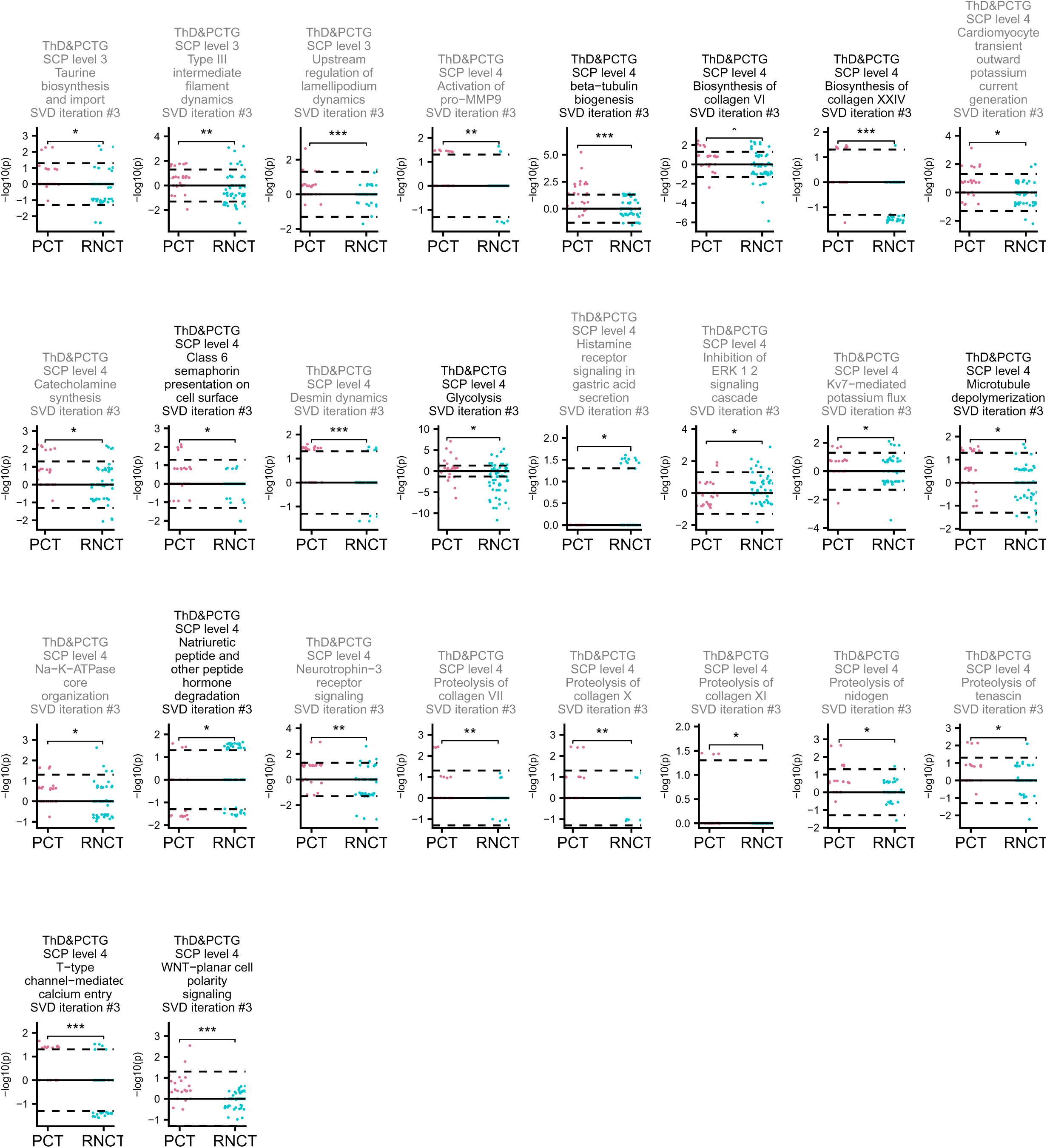
Pathways with induction level differences between persistent, reversible and no-CTRCD. -log_10_ P values (P ≤ 0.05) obtained by pathway enrichment analysis of downregulated final set-selective DEGs were subtracted from those obtained for upregulated final set-selective DEGs to calculate an overall induction level for each pathway. -log₁₀ P values of missing pathways were set to 0. Shown pathways documented significant variation in their induction levels in at least one comparison between persistent and non-persistent, reversible or no-CTRCD groups, as predicted by the Wilcoxon test and level- and comparison-selective Benjamini-Hochberg correction for the **(A)** PvsNP, **(B)** 3 CT groups and **(C)** T & PvsNP annotations. Only pathways that were significantly regulated in the same direction in at least 10% of all treatment and cell line combinations of at least one of the two compared CTRCD groups were tested. Black and gray headlines indicate pathways that were and were not consistently predicted as differentially induced, respectively. *, **, ***, ****: adjusted Wilcoxon P value ≤ 0.05, ≤0.01, ≤0.001, ≤0.0001. n.: no significance. Figure 4A shows representative level 3 pathways for the PvsNP annotation.

**Supplementary Figure 7.**
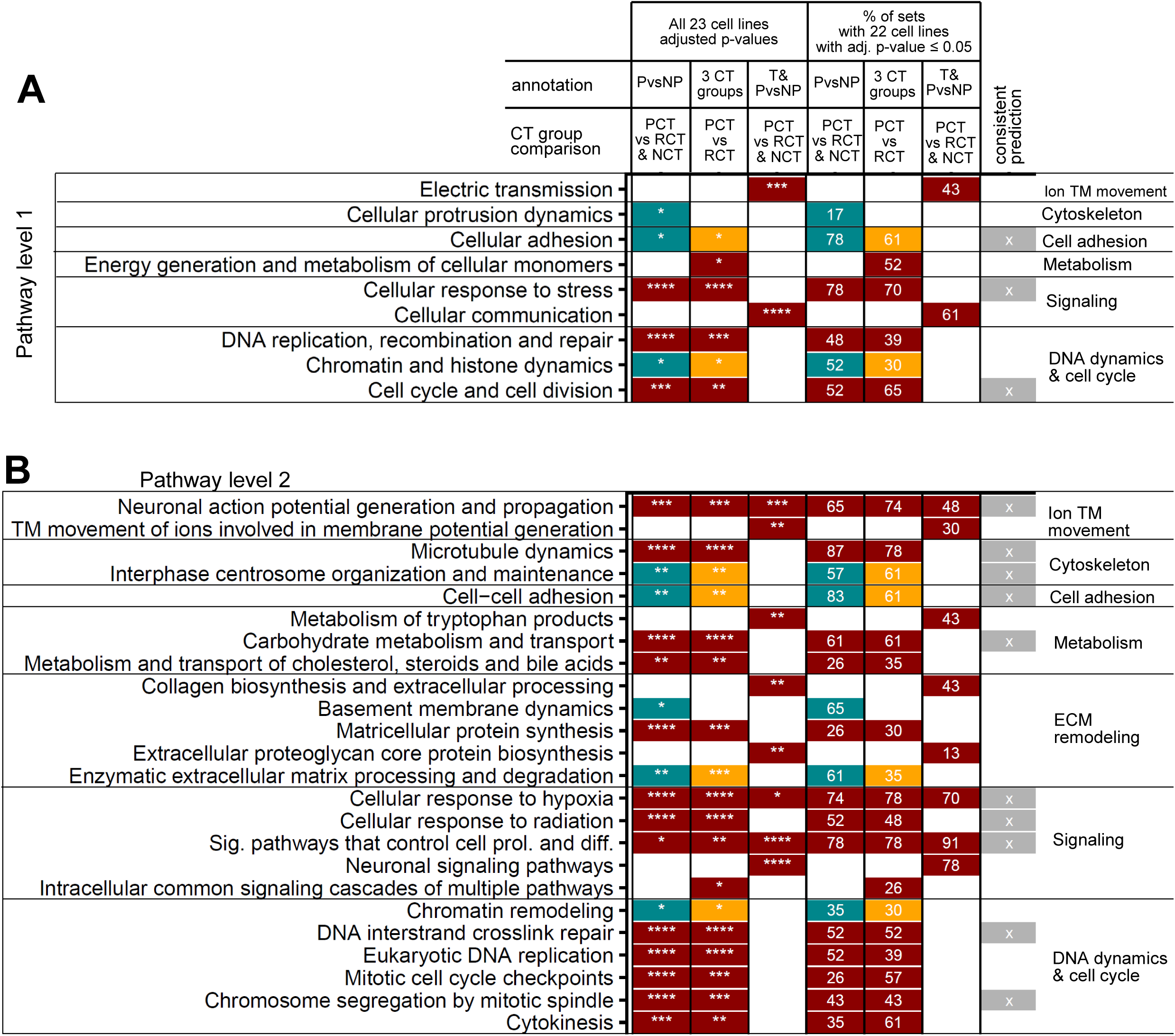
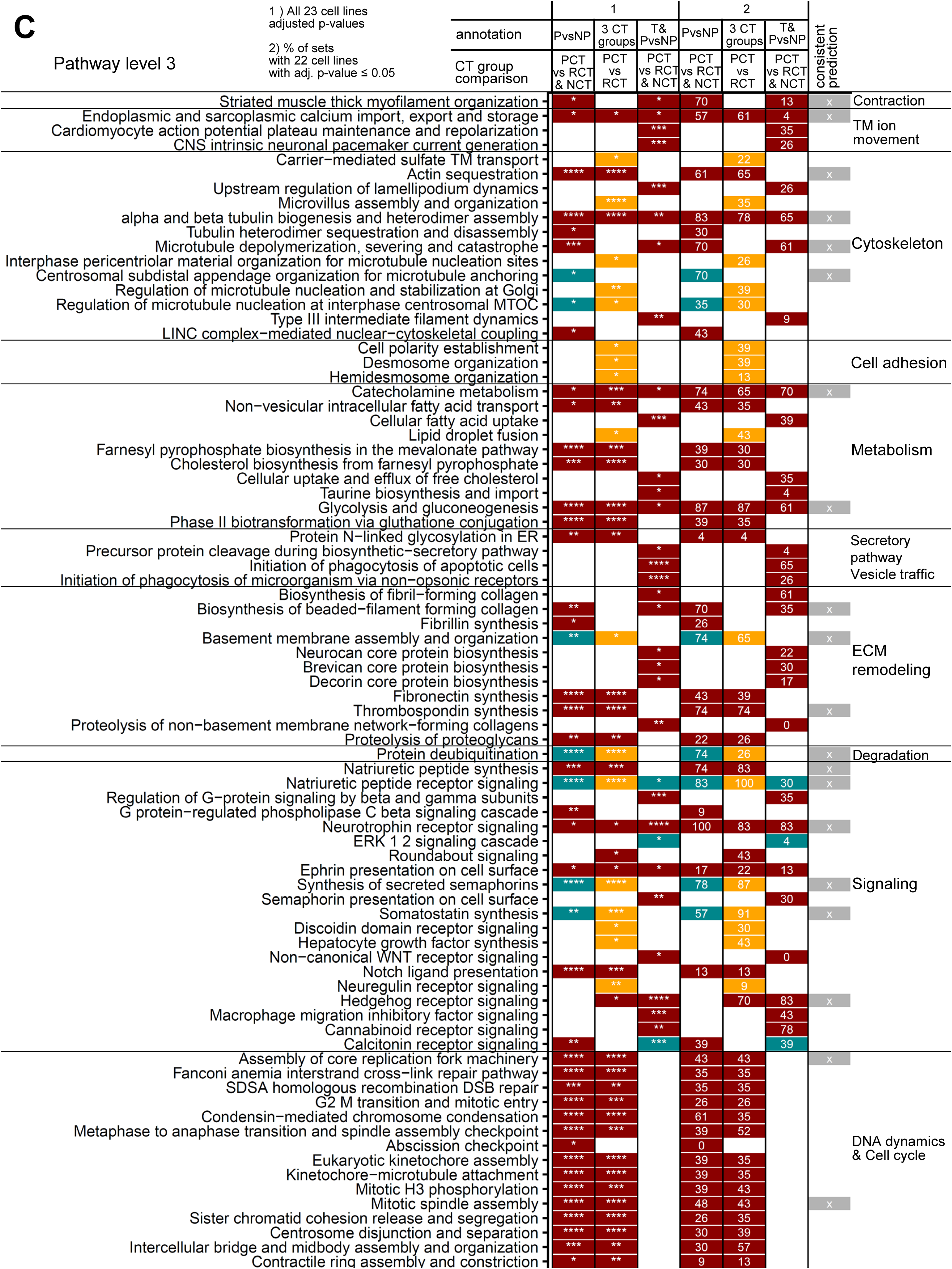
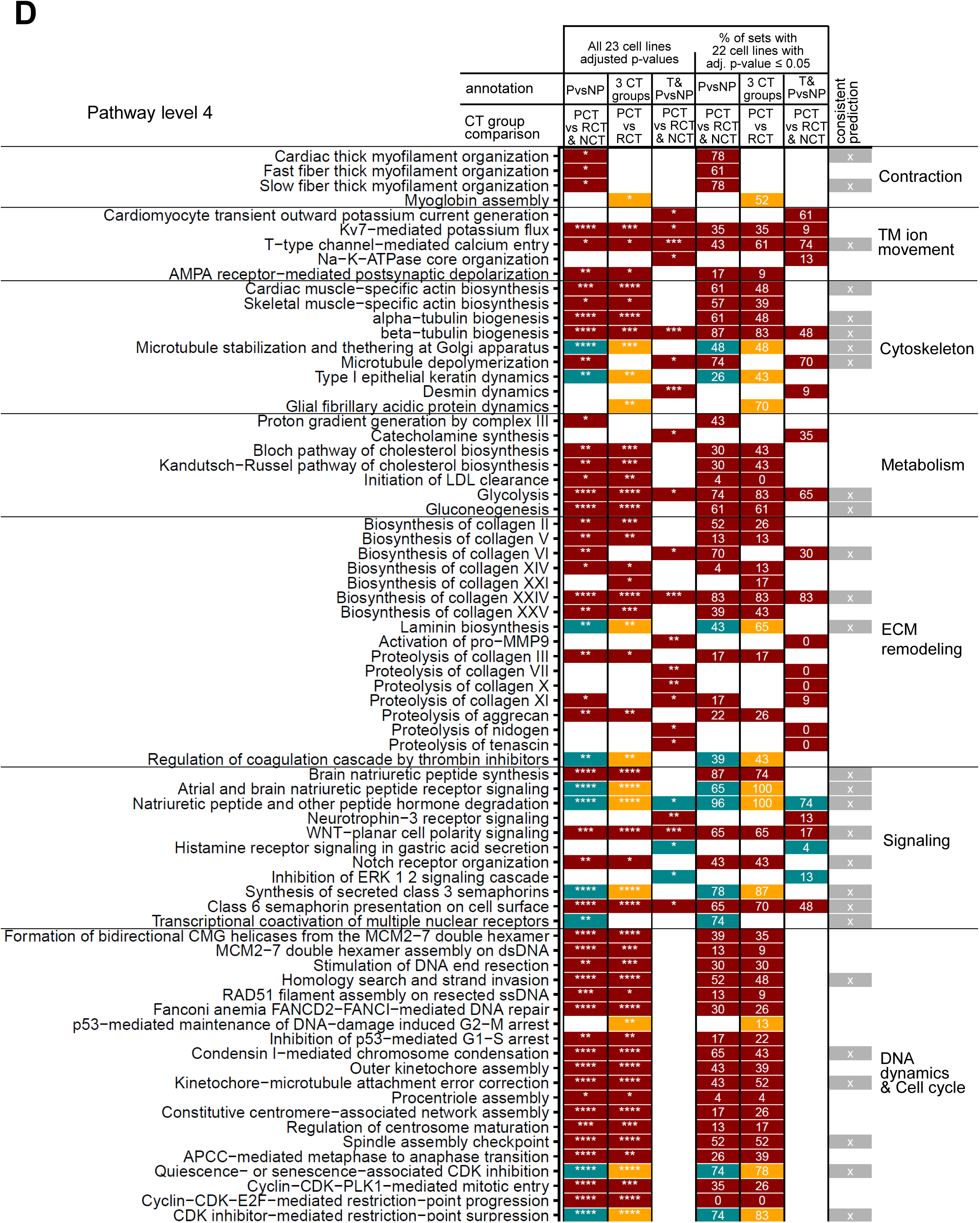
Pathways distinguishing persistent from non-persistent or reversible CTRCD. **(A)** Level 1, **(B)** level 2, **(C)** level 3 and **(D)** level 4 pathways with significantly different induction levels between persistent and non-persistent or reversible CTRCD groups that were predicted for the PvsNP, 3 CT groups or T & PvsNP annotation. The analysis was repeated with each of the 23 downsampled datasets obtained after removal of one cell line at a time. Percentage values document how often a pathway was predicted to show a significant variation between both groups (adjusted P ≤ 0.05). Colors label the condition with higher induction of a pathway for each pairwise comparison (dark red: persistent, turquoise: non-persistent, orange: reversible CTRCD). *, **, ***, ****: adjusted P value: ≤0.05, ≤0.01, ≤0.001, ≤0.0001. Consistently predicted pathways (labeled X) were identified based on the PvsNP annotation and in at least 70% of the related downsampled datasets or based on two annotations and in at least 50% of each set of related downsampled datasets. Consistently predicted level 3 pathways are also shown in Figure 4B.

**Supplementary Figure 8.**
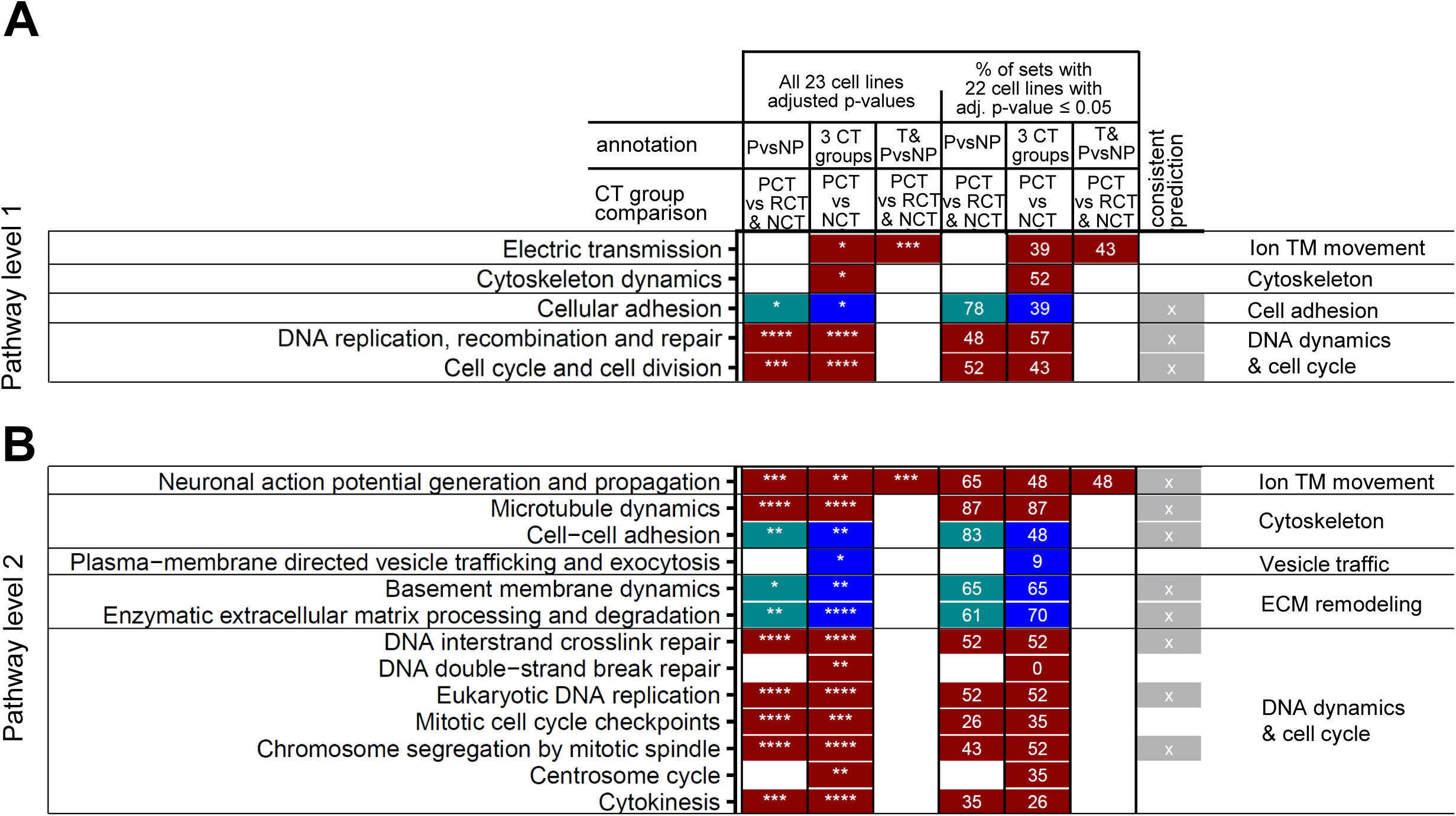
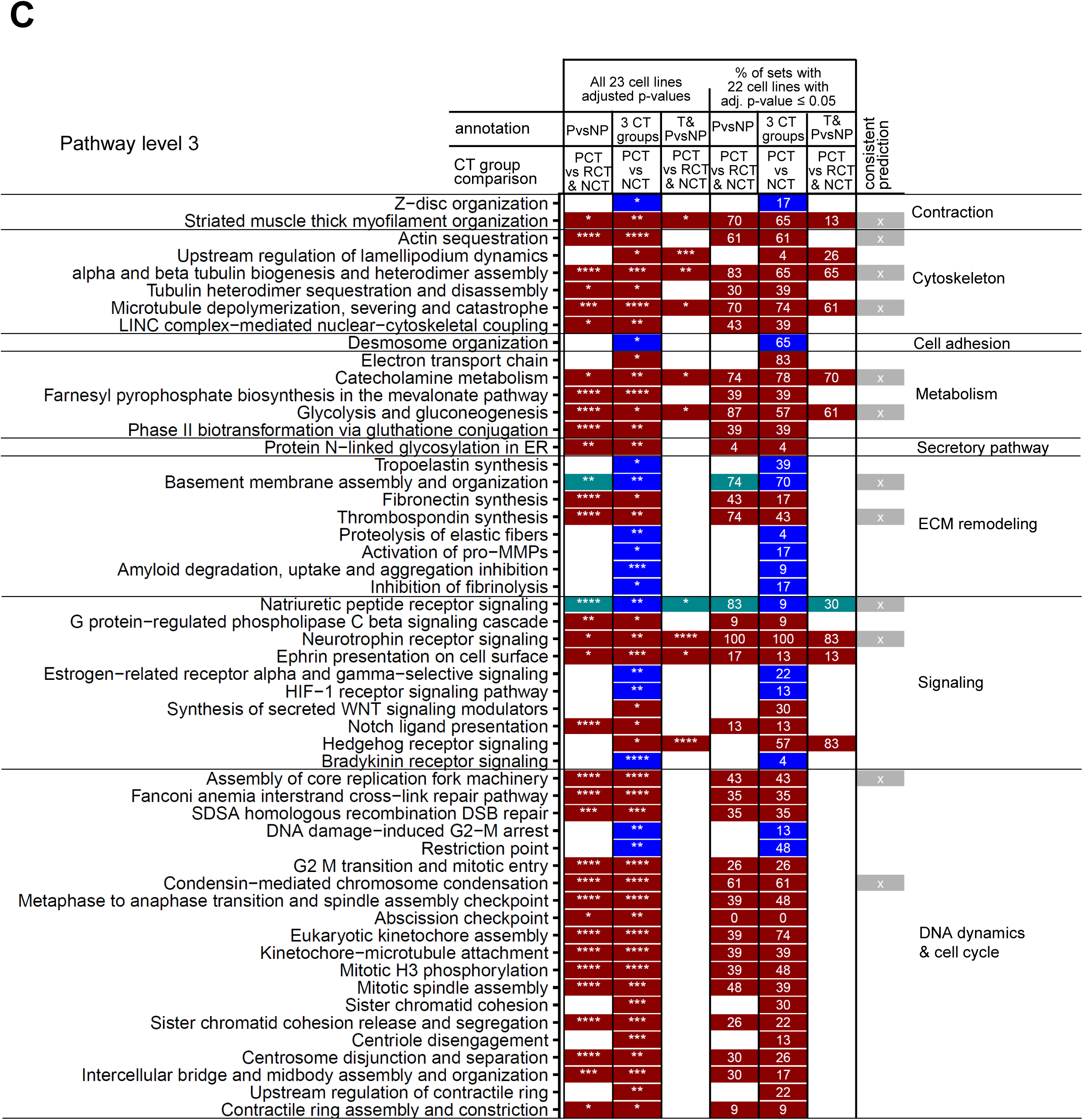
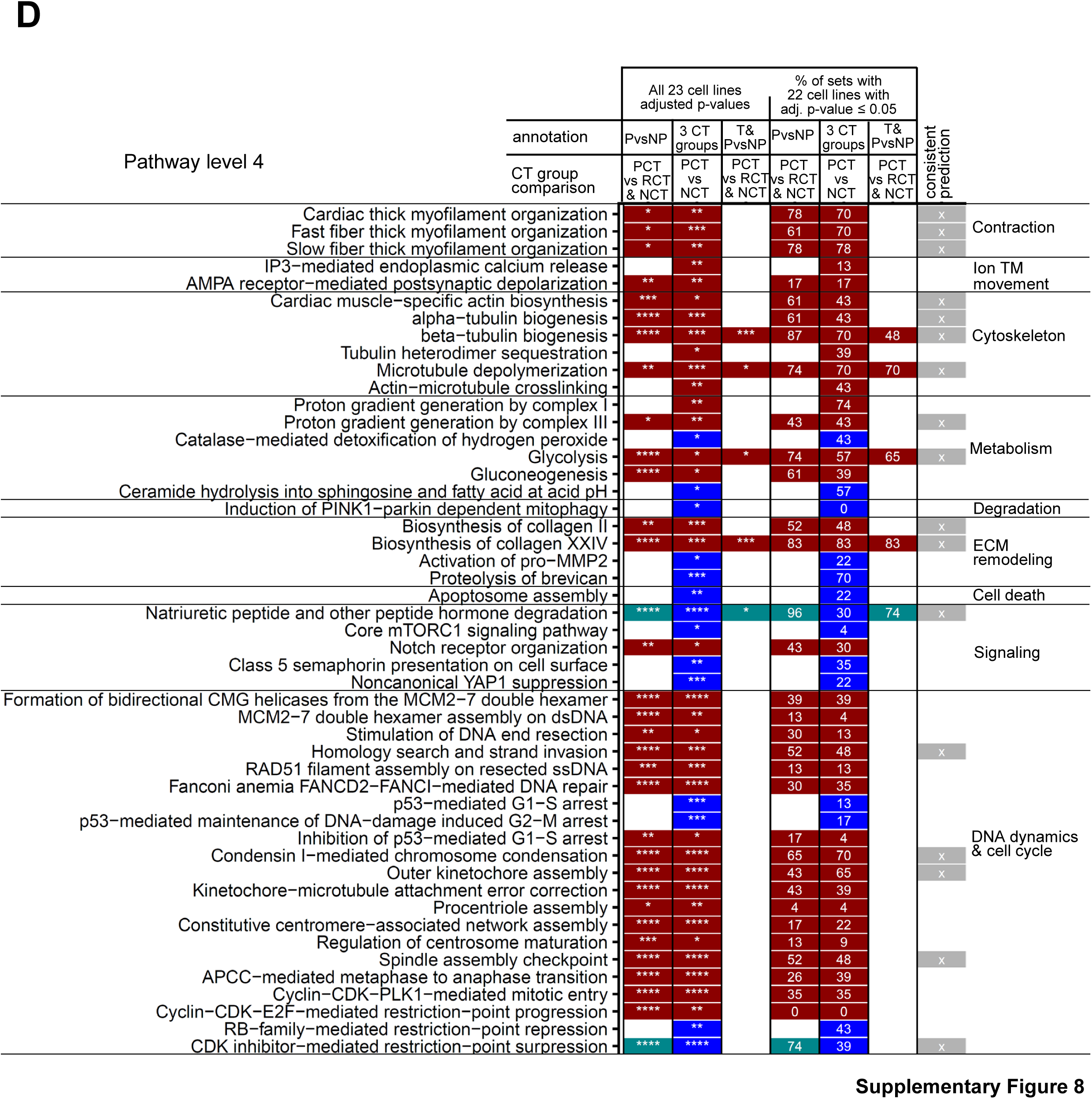
Pathways distinguishing persistent from no-CTRCD. **(A)** Level 1, **(B)** level 2, **(C)** level 3 and **(D)** level 4 pathways with significantly different induction levels between the persistent and no-CTRCD groups. Percentage values, symbols and consistency criteria as in Supplemental Figure 7; blue: higher induction in no-CTRCD lines.

**Supplementary Figure 9.**
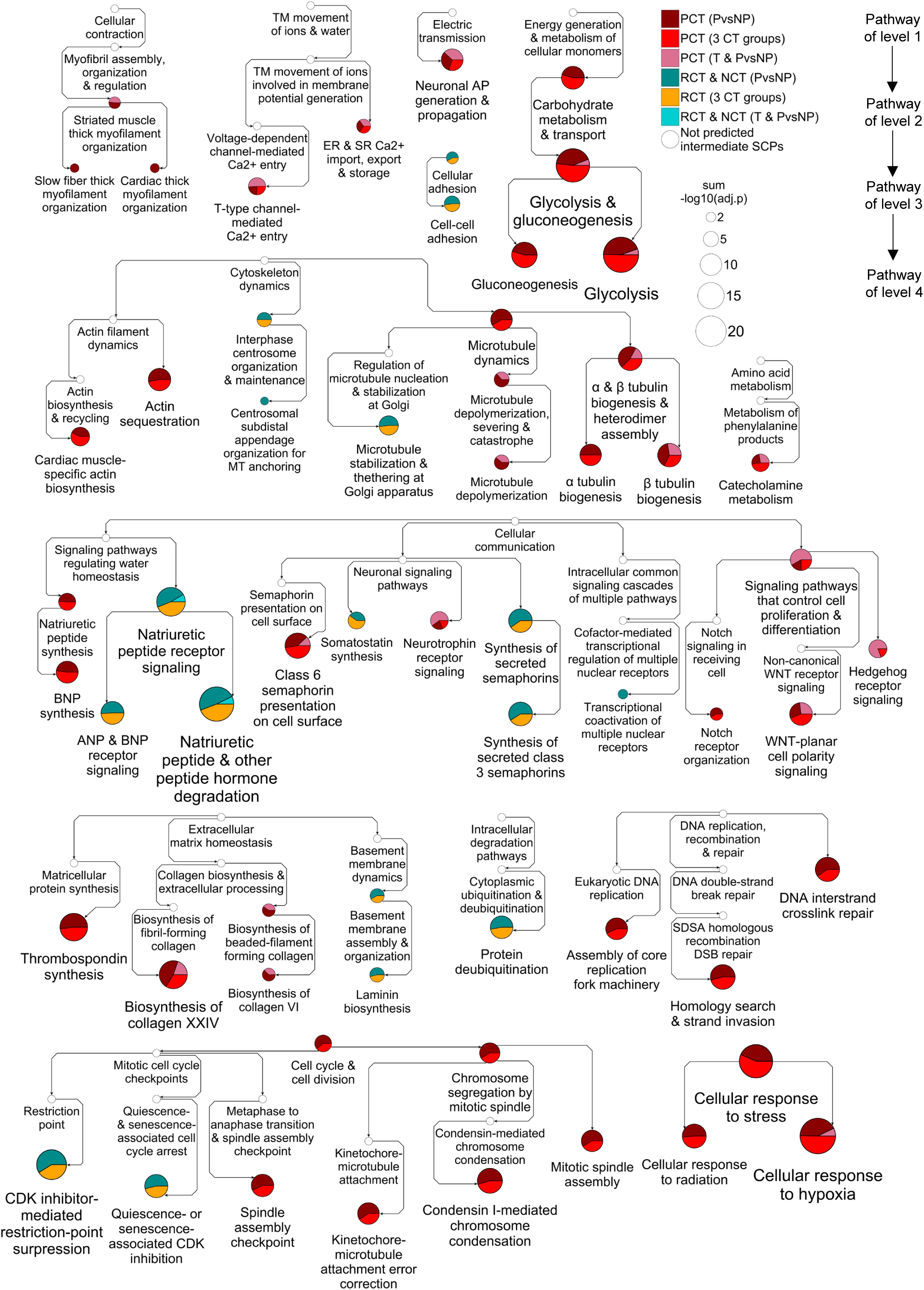
Hierarchical network of pathways distinguishing persistent from non-persistent or reversible CTRCD. Consistently predicted pathways with different induction levels between the persistent and non-persistent or reversible CTRCD groups were integrated into the MBCO v2 hierarchy. Pathway colors indicate the condition with higher induction levels and the underlying set annotation. Selected branches of this network are also shown in Figure 4C.

**Supplementary Figure 10.**
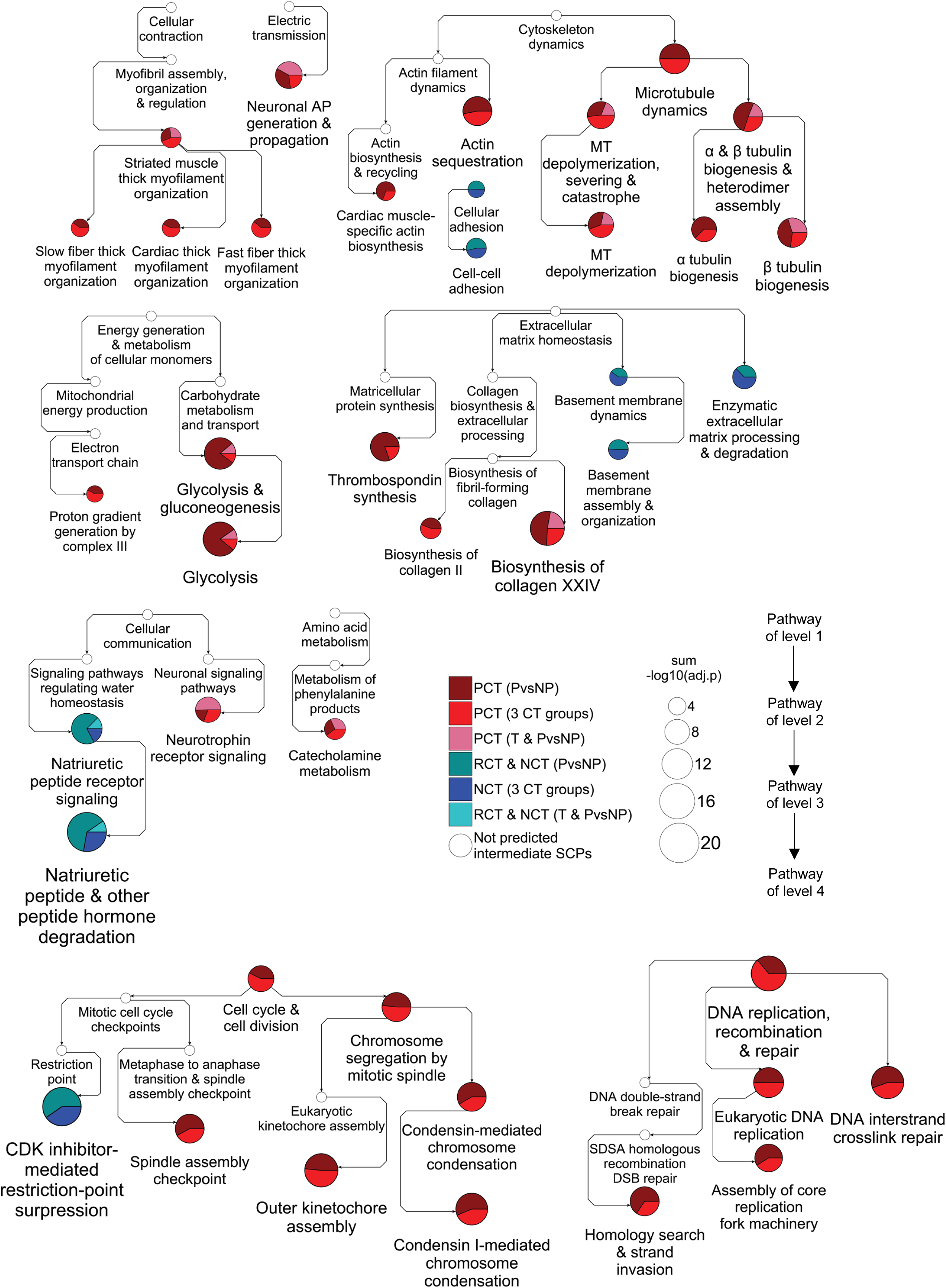
Hierarchical network of pathways distinguishing persistent from no-CTRCD. Consistently predicted pathways with different induction levels between the persistent and no-CTRCD groups were integrated into the MBCO v2 hierarchy. Pathway colors indicate the condition with higher induction levels and the underlying set annotation.

**Supplementary Figure 11.**
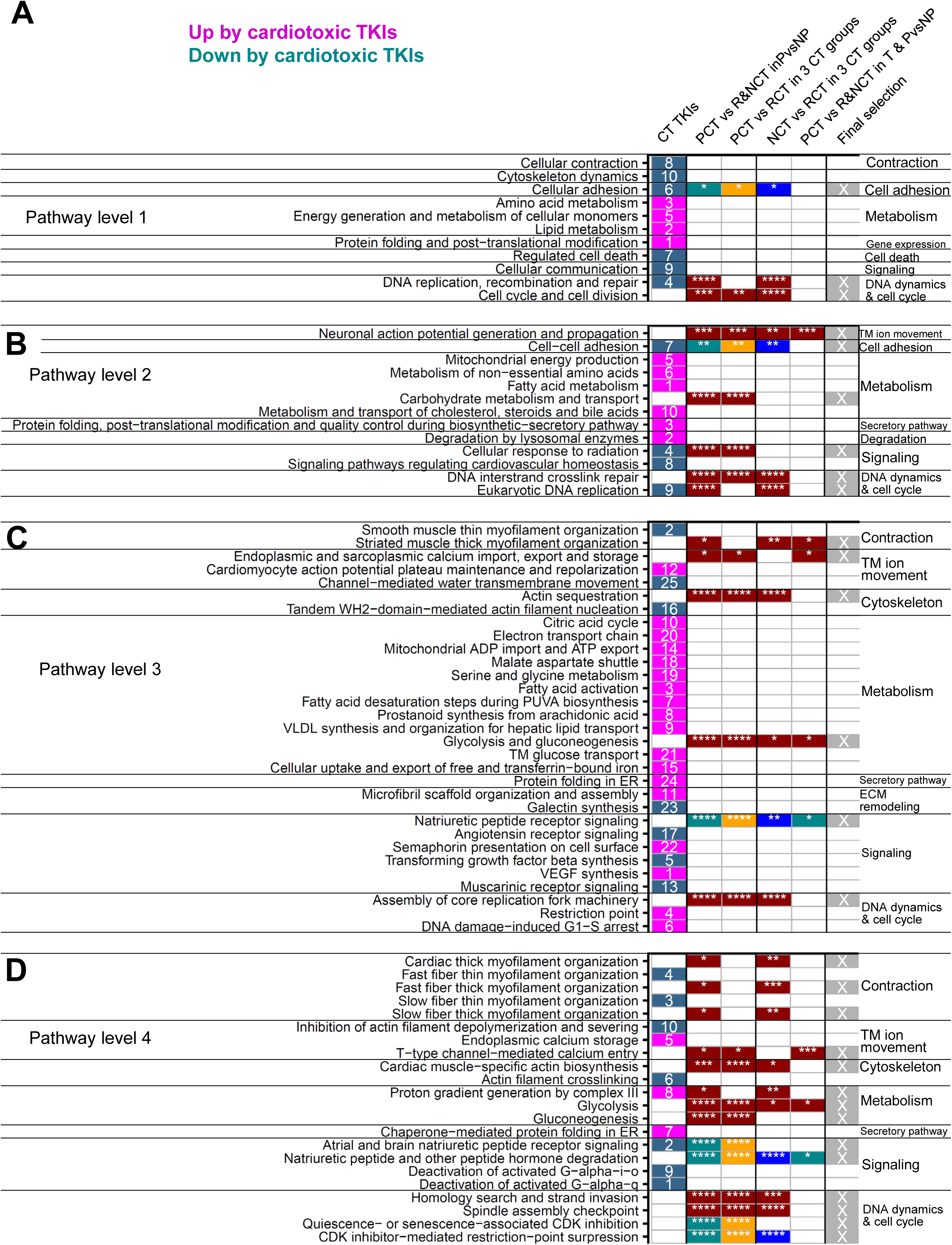
Mapping of cardiotoxic TKI pathways to pathways of this study. The top **(A)** 10 level 1, **(B)** 10 level 2, **(C)** 25 level 3 and **(D)** 10 level 4 MBCO v2 pathways that are up- or downregulated with higher ranked significance by cardiotoxic than by non-cardiotoxic tyrosine kinase inhibitors (TKIs) in healthy human iPSC derived cardiomyocytes were mapped to consistently predicted pathways identified in this study. All TKI pathways are shown. Shown pathways of this study were either identical or in family relationships (siblings, ancestors, descendants) with the TKI pathways. Numbers indicate selection ranks for TKI pathways. */**/***/****: adjusted Wilcoxon P value ≤ 0.05/0.01/0.001/0.0001. Dark red, turquoise, orange, blue: induced with to a greater extent in persistent, non-persistent, reversible or no-CTRCD lines, respectively. Magenta, gray-blue: Up- or down-regulated by cardiotoxic TKIs, respectively.

**Supplementary Figure 12.**
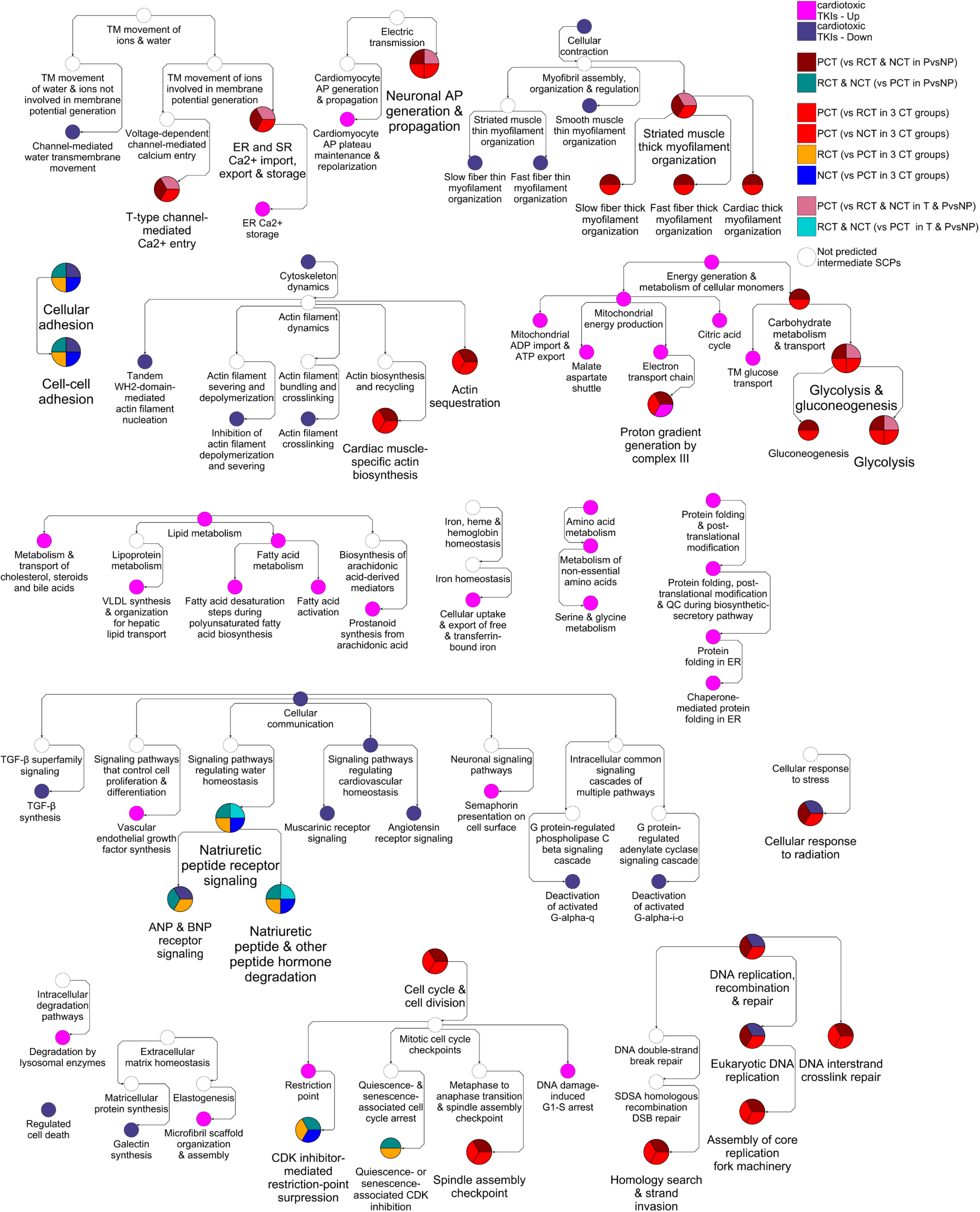
Integration of TKI pathways into the MBCO hierarchy. Integration of all TKI pathways and selected pathways identified in this study into the MBCO v2 hierarchy documents parent-child and sibling relationships. Selected branches of this network are also shown in Figure 5.

**Supplementary Figure 13.**
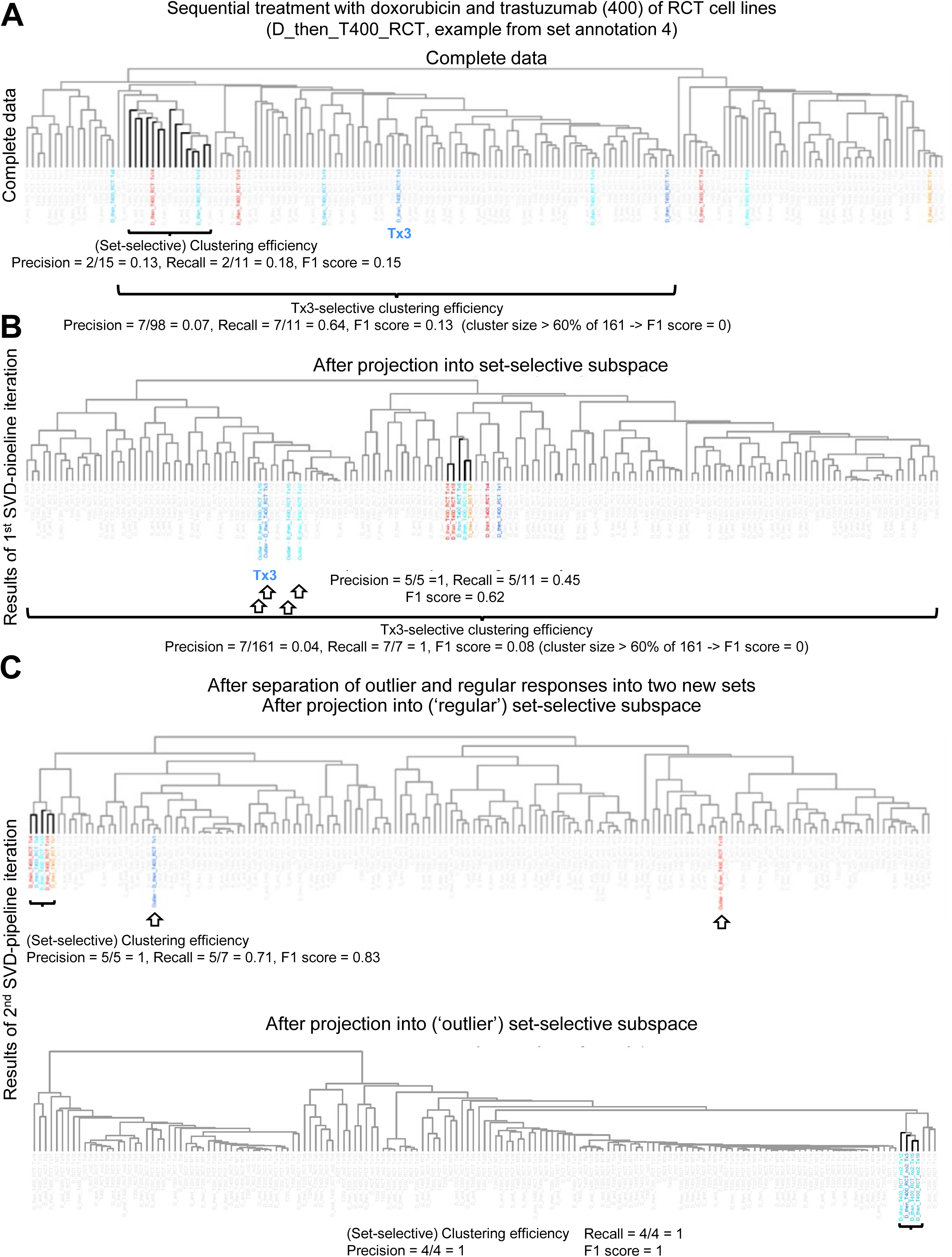
Example subspace projection across iterations of the SVD-pipeline. **(A)** 161 complete DEG lists were subjected to pairwise correlation analysis, followed by hierarchical clustering. To quantify the set-selective clustering efficiency (here shown for the set of reversible CTRCD lines sequentially treated with doxorubicin (D) and high-dose trastuzumab (T 400)) our pipeline cut the dendrogram at any possible height to identify any possible cluster. For each cluster, it calculated the fraction of DEG responses belonging to that set among all responses in that cluster (precision) and the fraction of all responses of that set that were in that cluster (recall). Both values allowed calculation of the F1 score (beta = 1). The highest F1 score (bold cluster) is the set-selective clustering efficiency in the complete data. For each DEG vector of a set, our algorithm additionally calculates the individual clustering efficiency, illustrated here for cell line Tx3, as the highest F1 score obtained for a cluster that contains the response in question and the set-selective F1 score cluster. The low set-selective clustering efficiency suggests that set-selective responses are overlaid by set-unrelated responses. Blue and red label cell lines obtained from patients with fast and slow recovery, respectively. **(B)** To extract set-selective response vectors our pipeline applies SVD and searches for set-selective subspaces with optimized clustering efficiencies and minimized numbers of outlier responses for all DEG responses of a set in question (see methods for details). Within the identified set-selective subspace for the example set, the clustering efficiency increased to 0.62. Arrows label outlier responses. **(C)** Regular and outlier responses were grouped into two new independent sets, followed by identification of subspaces for each new set using the same algorithm. The second iteration led to an additional increase of the clustering efficiencies. One outlier response was still identified for the ‘regular’ set. Because new sets require at least 4 members for any new set that constitute at least 20% of the parent set vectors, the algorithm did not generate a new set for that outlier response.

**Supplementary Figure 14.**
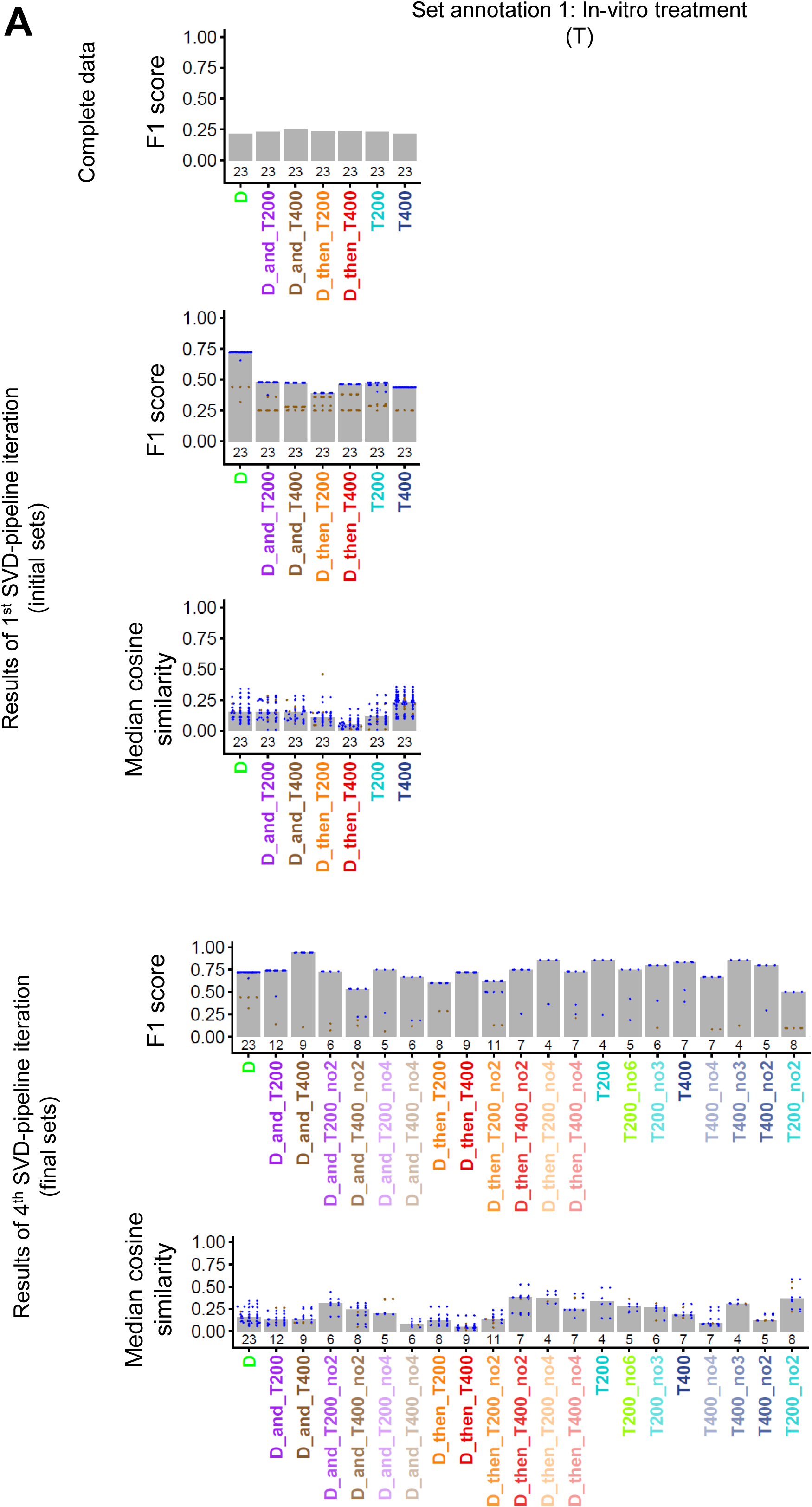
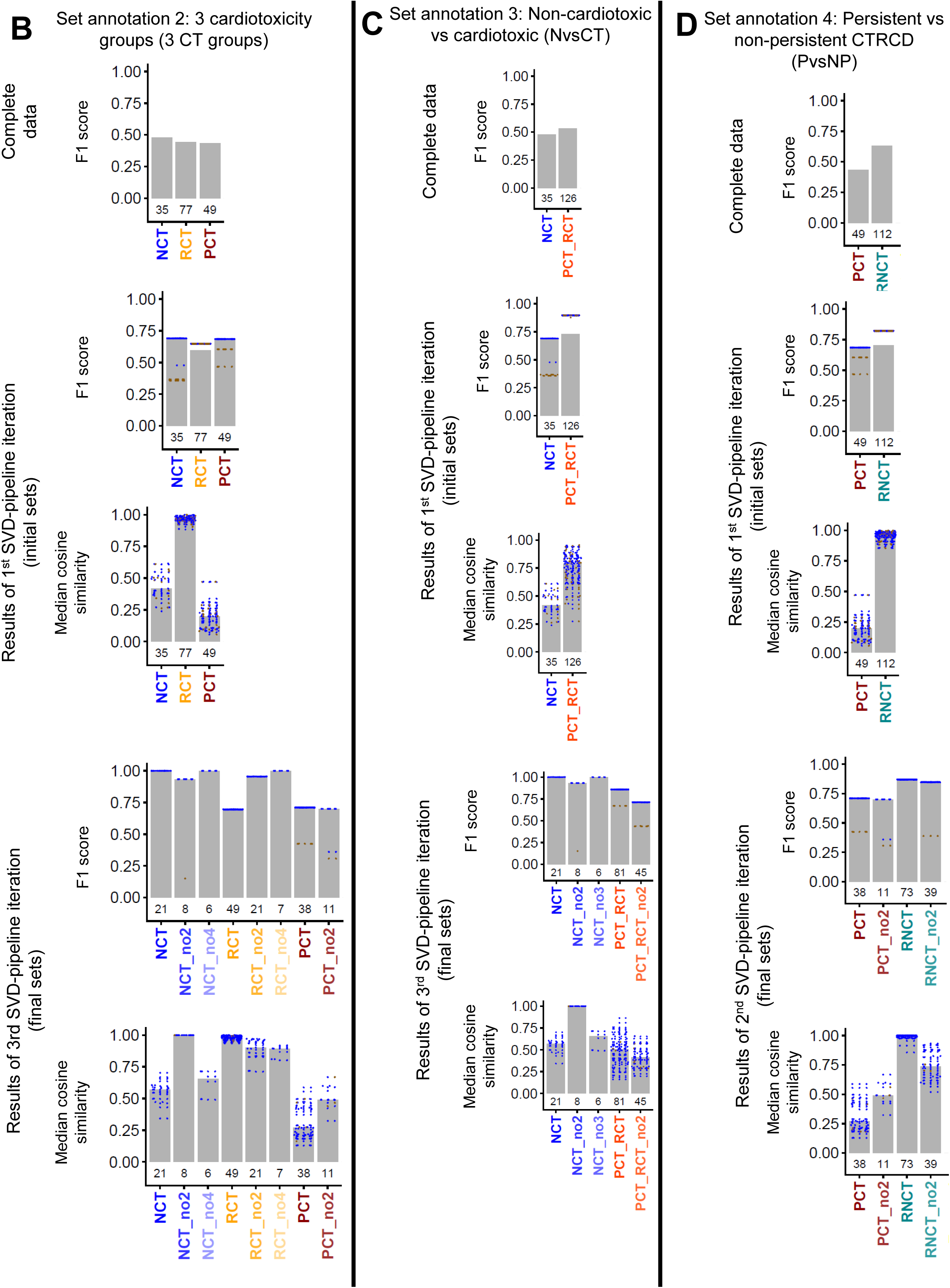
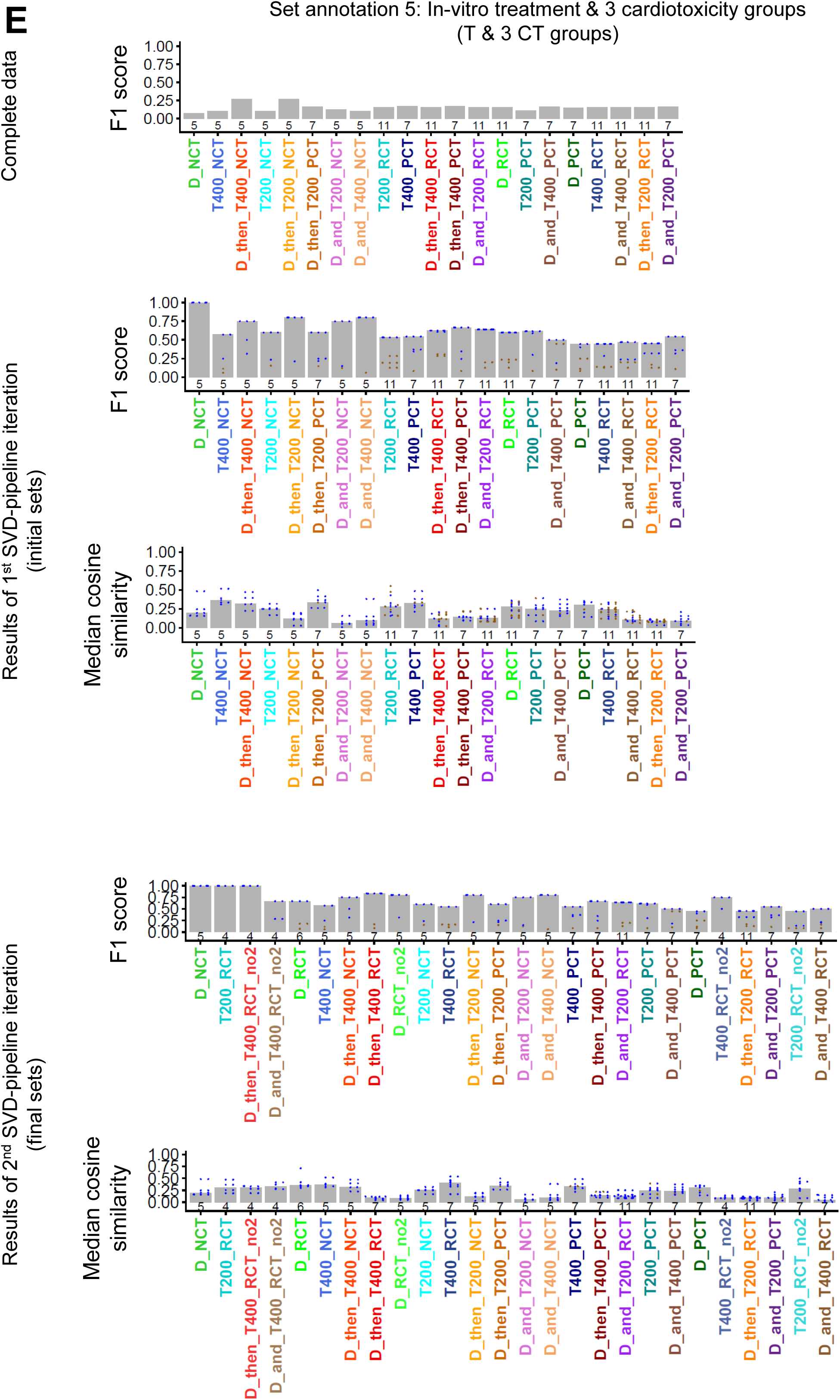
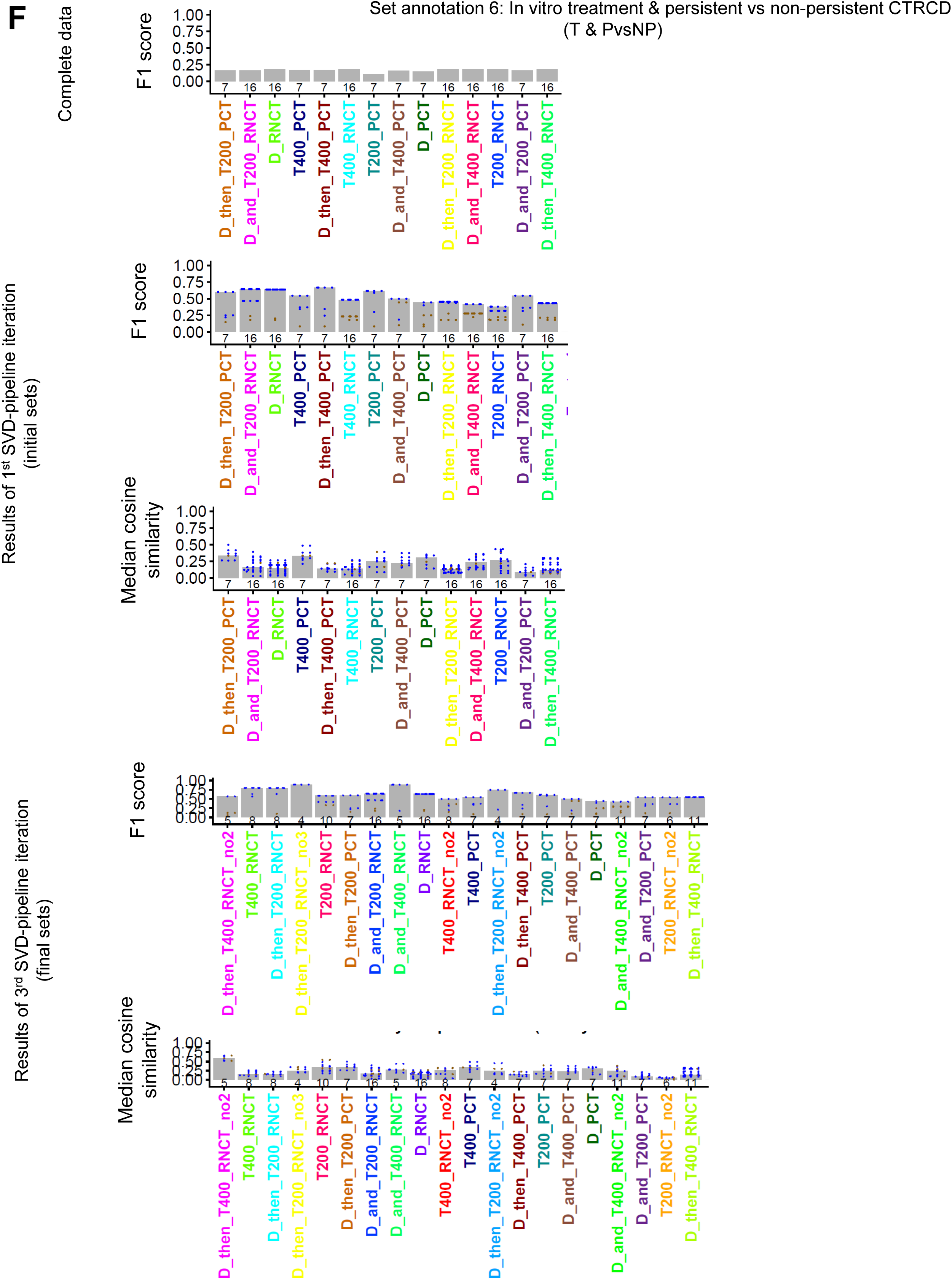
Clustering efficiency and information preservation. For the. **(A)** T, **(B)** 3 CT groups, **(C)** NvsCT, **(D)** PvsNP, **(E)** T & 3 CT groups and **(F)** T & PvsNP annotations, the SVD pipeline identified set-selective subspaces with minimal outlier responses and maximal clustering efficiencies. Gray bars show clustering efficiencies and the median cosine similarities between the DEG vectors of each set in the set-selective subspaces and the complete data. Blue and brown dots show the clustering efficiencies and cosine similarities obtained for each regular and outlier DEG vector annotated to the set in question. Horizontal blue lines show the median of the gray bars. Numbers of DEG responses within each set are added above the abscissa. Panel titles use earlier annotation names: CTG = 3 CT groups; NCTG = NvsCT; PCTG = PvsNP; T&CTG = T & 3 CT groups; T&PCTG = T & PvsNP; RNCT = reversible and no CTRCD. D: doxorubicin; T200, T400: low- and high-dose trastuzumab.

